# Multimodal Spatial Profiling Reveals Immune Suppression and Microenvironment Remodeling in Fallopian Tube Precursors to High-Grade Serous Ovarian Carcinoma

**DOI:** 10.1101/2024.09.25.615007

**Authors:** Tanjina Kader, Jia-Ren Lin, Clemens Hug, Shannon Coy, Yu-An Chen, Ino de Bruijn, Natalie Shih, Euihye Jung, Roxanne J. Pelletier, Mariana Lopez Leon, Gabriel Mingo, Dalia Khaled Omran, Jong Suk Lee, Clarence Yapp, Baby Anusha Satravada, Ritika Kundra, Yilin Xu, Sabrina Chan, Juliann B. Tefft, Jeremy Muhlich, Sarah Kim, Stefan M. Gysler, Judith Agudo, James R. Heath, Nikolaus Schultz, Charles Drescher, Peter K Sorger, Ronny Drapkin, Sandro Santagata

**Author notes:** These authors contributed equally (Tanjina Kader, Jia-Ren Lin, Clemens Hug, Ronny Drapkin, Sandro Santagata). **DECLARATION OF INTERESTS** PKS is a co-founder and member of the BOD of Glencoe Software and member of the SAB for RareCyte, NanoString, and Montai Health; he holds equity in Glencoe and RareCyte. PKS is a consultant for Merck. RD is a member of the SAB for Repare Therapeutics and is a consultant for Light Horse Therapeutics and Abbvie. The other authors declare no outside interests.

## Abstract

High-Grade Serous Ovarian Cancer (HGSOC) originates from fallopian tube (FT) precursors. However, the molecular changes that occur as precancerous lesions progress to HGSOC are not well understood. To address this, we integrated high-plex imaging and spatial transcriptomics to analyze human tissue samples at different stages of HGSOC development, including p53 signatures, serous tubal intraepithelial carcinomas (STIC), and invasive HGSOC. Our findings reveal immune modulating mechanisms within precursor epithelium, characterized by chromosomal instability, persistent interferon (IFN) signaling, and dysregulated innate and adaptive immunity. FT precursors display elevated expression of MHC-class I, including HLA-E, and IFN-stimulated genes, typically linked to later-stage tumorigenesis. These molecular alterations coincide with progressive shifts in the tumor microenvironment, transitioning from immune surveillance in early STICs to immune suppression in advanced STICs and cancer. These insights identify potential biomarkers and therapeutic targets for HGSOC interception and clarify the molecular transitions from precancer to cancer.

**STATEMENT OF SIGNIFICANCE:** This study maps the immune response in fallopian tube precursors of high-grade serous ovarian cancer, highlighting localized interferon signaling, CIN, and competing immune surveillance and suppression along the progression axis. It provides an explorable public spatial profiling atlas for investigating precancer mechanisms, biomarkers, and early detection and interception strategies.

## INTRODUCTION

High-Grade Serous Ovarian Carcinoma (HGSOC) is a highly aggressive gynecological cancer, causing over 200,000 deaths worldwide each year^1^. The absence of early-stage symptoms often results in diagnosis at advanced stages (AJCC Stage III & IV) when the cancer has already spread. The 5-year overall survival rate for patients with advanced disease is less than 30% and has remained unchanged for decades^2^. Standard treatment includes surgery to reduce tumor burden, followed by chemotherapy, but chemoresistance is common, with over 80% of stage III or IV patients experiencing relapse after initial treatment. This underscores the need for early detection and interception strategies to identify the disease at its earliest stages, reduce recurrence, and improve outcomes^3^. Women with *BRCA* gene mutations are at particularly high risk for developing HGSOC, leading some women to choose preventive surgery, such as bilateral salpingectomy-oophorectomy (BSO) to reduce their risk^3^.

Over the past two decades, studies have shown that HGSOC originates at the distal, fimbriated end of the fallopian tube (FT)^4–6^. Histopathologic evaluation and next-generation sequencing have recognized two types of precursor lesions for HGSOC: the p53 signature and serous tubal intraepithelial carcinomas (STIC). p53 signatures are benign-appearing stretches of secretory cells with *TP53* mutations that are non-proliferative and commonly located at the fimbriated end of the FT, regardless of genetic risk. The origin of these p53 signatures is linked to the ‘incessant ovulation hypothesis’, which proposes that continuous ovulation leads to repeated injury to cells near ruptured follicles, including those at the fimbriated end of the FT^7^. The repeated injury leads to genetic and epigenetic alterations, transforming p53 signatures into STIC and eventually into invasive cancer^8^. Incidental STICs discovered during risk-reduction surgeries have been associated with the subsequent development of peritoneal carcinomatosis^9^, suggesting that STIC cells may shed from the fimbria before invasion occurs (a precursor escape model^10^). However, the full progression of these precursors to invasive cancer remains complex and is not yet fully understood.

Genomic studies have demonstrated a clonal relationship between p53 signatures, STICs, and concurrent HGSOC, with each sharing identical *TP53* mutations^11,12,13^. Nearly half of HGSOCs exhibit defects in homologous recombination, resulting in genomic instability due to somatic mutations in the *BRCA1/2* genes, epigenetic silencing of *BRCA1/2* promoters, or mutations in other DNA repair factors^3^. HGSOC also commonly displays chromosomal instability (CIN), including breakage-fusion-bridge cycles that lead to the amplification of key oncogenes like *CCNE1*^14^. DNA methylation analyses further support the FT as the origin of HGSOC, as the methylation profile of HGSOC more closely resembles that of FT epithelium rather than ovarian surface epithelium^15,16^. In addition, HGSOC-specific hypermethylation is present exclusively in the FT epithelium of women with STIC lesions^17^. Early events in STICs include similar global patterns of copy number alterations, such as *CCNE1* amplification and higher ploidy^11–13,18–21^.

Despite progress in understanding these genetic features, the role of the immune microenvironment in HGSOC development and progression remains underexplored. Among the four subtypes of HGSOC, the immunoreactive subtype, characterized by CD8+ cytotoxic T cell (CTL) infiltration, is associated with a better prognosis^22,23^. By contrast, subtypes such as the C5/PRO subtype often exhibit immune deserts or CTL exclusion, which are linked to poorer outcomes^24^. Single-cell RNA sequencing of advanced HGSOC has identified several cancer-related pathways, including Janus kinase (JAK)-signal transducer and activator of transcription (STAT) signaling, the interferon (IFN) response, inflammatory pathways, and transforming growth factor β (TGF-β) signaling, all of which correlate with T cell infiltration levels^25^. The female reproductive tract (including the FTs) has a highly active immune system, rich in natural killer (NK) and T cells, which are likely involved in maintaining immune tolerance during pregnancy and monitoring microbial infections^26,27^. Thus, gaining a better understanding of how the immune microenvironment changes during HGSOC progression could unlock key insights for improving disease diagnosis and management.

In this study, we used a multi-omic approach to map the tumor-immune ecosystems during the development and progression of HGSOC. We leveraged several high-plex spatial analysis methods, such as cyclic immunofluorescence (CyCIF^28^), 3D CyCIF^29^, and spatial transcriptomics (whole transcriptome analysis, WTA)^30^ of human archival tissue specimens to map the spatial distributions, interactions, and molecular programs of different cell types. This data reveals key changes that occur in the tumor microenvironment (TME) as precancer STIC lesions progress to invasive HGSOC. We uncover temporal changes in molecular pathways, including activation of interferon (IFN) signaling, micronuclei (MN) formation and rupture, and cyclic GMP-AMP synthase (cGAS)-stimulator of interferon genes (STING) signaling. CyCIF imaging reveals dynamic shifts in immune cell populations and interactions over the progression axis. Early lesions are defined by immune surveillance, characterized by the presence of conventional dendritic cells Type 1 (cDC1), NK cells, and tissue-resident memory (T_RM_) CD8+ T cells. However, in advanced precursor lesions, we observe a significant decline in these immune cells, along with molecular evidence of immune dysfunction and immune editing. By combining spatially transcriptomics and high-plex imaging, this study highlights the dynamic interplay between immune activation, suppression, and molecular reprogramming during HGSOC progression. We have made our data publicly available and explorable through cBioPortal, providing a widely accessible resource for identifying therapeutic targets and informing early detection strategies.

## RESULTS

### Specimen Cohort

To investigate the molecular and spatial changes occurring in the fallopian tube (FT) during the early development of high-grade serous carcinoma (HGSOC), we analyzed 44 FT specimens with precursor lesions collected from 43 individuals obtained from a multi-center collaboration (**Fig. 1A-1C)**. The 44 specimens were collected from patients with various genotypes and disease presentations, which we categorized into two main groups based on the presence or absence of cancer. Specimens in Group 1 (n=24) contained invasive cancer and co-occurring serous tubal intraepithelial carcinoma (denoted as STIC.C). This included specimens from individuals with and without *BRCA* mutations (wild-type (WT) *BRCA* 15/24; germline (g) *BRCA* n=7/9; g*BRCA2* n=6/7; somatic 2/9) (**Fig. 1B, 1D**). Specimens in Group 2 (n=19) lacked invasive cancer but contained precursor lesions that were identified during risk-reducing BSO or opportunistic salpingectomy. Group 2 included patients with and without BRCA mutation, and specimens contained incidental p53 signatures (denoted as p53.I, n=10; g*BRCA*1 5/10; g*BRCA*2 5/10) and incidental STIC lesions (denoted as STIC.I, n=9; g*BRCA* 5/9; g*BRCA*2 1/5; WT *BRCA* 4/9) (**Fig. 1B-1D**). STIC.I likely represent early time points in clonal evolution, while STIC.C represents later points in the development and progression of STIC lesions^12,31,32^. Of the 44 specimens, all but nine had matched FT and/or fimbriae (Fim) within the same tissue section (**Fig. 1C**).

**Figure 1:**
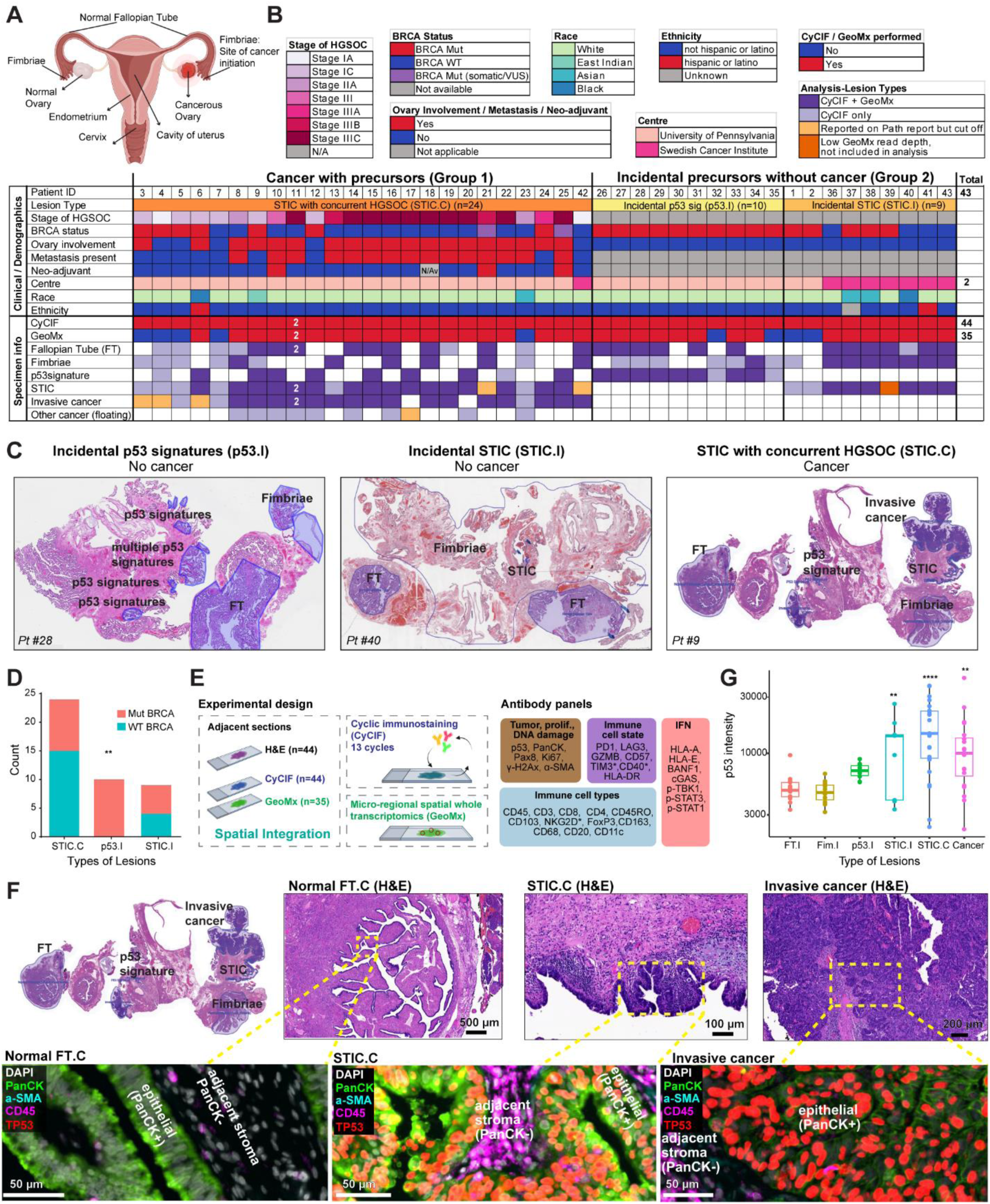
Overview of the cohort and experimental design. **A.** Anatomy of female reproductive tract showing Fallopian Tube (FT), distal end of the FT, Fimbriae and ovary. It is now generally accepted that the majority of High Grade Serous Ovarian carcinomas (HGSOC) arise from the secretory cells in the distal fimbriated end of FT. **B.** Detailed clinical annotation of the entire cohort of 43 patients, indicating 44 specimens run on tissue Cyclic Immunofluorescence (CyCIF) and 35 specimens on spatial whole transcriptomics (GeoMx) platform (NanoString). Clinical annotations are provided (HGSOC stage, BRCA mutation status, ovarian involvement, metastasis presence, and neoadjuvant chemotherapy), as are basic clinical details (race, ethnicity). For all samples, the type of lesion (i.e., histology) was recorded and the disappearance of lesions on subsequent sections from H&E was also noted. See **Supplementary File S1** for the complete table. **C.** Example of H&E from each subgroup of the cohort: Incidental p53 signatures (p53.I), incidental STIC (STIC.I) and STIC with concurrent cancer (STIC.C). **D.** Stacked bar plot comparing number of cases of *BRCA* mutant (Mut) and wild-type (WT) between incidental and cancer-associated precancer lesions, **p<0.01, Fisher exact test. **E.** Experimental design and spatial integration of histology-guided multiplex tissue imaging, CyCIF, and GeoMx. The adjacent sections (5 µM) were chosen for CyCIF and GeoMx. Region of Interests (ROIs) of GeoMx were then integrated into CyCIF images based on the X/Y coordinates of both sections (see **Supplementary Figure S2** and **Methods**). The antibody panel shows 31 antibodies used for analysis in this study; asterisks indicate antibodies that were run only on a subset of the specimens (n=26 out of 44). High-resolution 3D CyCIF was performed for one STIC.C case, shown in Figure 1F (patient ID 9, case RD-23-002). **F.** Example STIC with concurrent HGSOC (Case RD-23-002, patient ID 9, BRCA2 mutant, Stage IC HGSOC). H&E images (top row) show different histologies within the specimen: FT.C, STIC.C, and invasive cancer. The CyCIF images (bottom row) show increased p53 mutant epithelial cells with disease progression, as well as the epithelial compartment (PanCK+) and the adjacent stromal compartment (PanCK-). **G.** Box plot comparing p53 intensity level in epithelial cells across disease stages, as quantified from tissue imaging. Y axis is presented in log10 scale. Number of specimens per group as follows: FT.I (n=13), Fim.I (n=15), p53.I (n=10), STIC.I (n=9), STIC.C (n=23), and Cancer (n=20). The solid line indicates the median within the interquartile range, with whiskers extending to a maximum of 1.5 times the interquartile range beyond the box. Outliers shown. Black asterisks indicate significant differences in stages compared to the FT.I (**p<0.01, ****p<0.0001), as calculated with Linear Mixed Models (LMMs) with patient ID as random effect. LMMs were implemented in the lme4 R package (v 4.3.3). **A, E.** Created with BioRender.

Among the 24 patients with invasive cancer, eight had stage I disease (6/8 with gBRCA mutations). Five of these stage I patients had tumors restricted to the FT, with no involvement of the ovary or spread through the abdomen (peritoneal metastasis) (**Supplementary Fig. S1A, S1B**). The remaining 16 patients had stage II-III disease (13/16 *BRCA* WT). Only three cancer patients received neoadjuvant chemotherapy, while the rest had surgery without prior chemotherapy (treatment-naïve; **Fig. 1B**). Patients with invasive cancer co-occurring with STIC lesions (STIC.C) were much older with a median age (65.5 years, range 46-85) (**Supplementary Fig. S1C**) than individuals with incidental precursor lesions (p53.I or STIC.I; median age of 47 years, range 34-72). The younger age of individuals with incidental lesions reflects earlier timing of BSO in women with known *BRCA* mutations. Overall, this cohort represents the entire spectrum of HGSOC development, enabling the characterization of disease progression (**Fig. 1B**, **Supplementary File S1**).

### Precursor and Cancer Analysis with CyCIF and Spatial Transcriptomics

We analyzed tissue sections from all 44 specimens using hematoxylin and eosin (H&E) staining and CyCIF^28^, which was performed using a panel of 31 antibodies and quantified to reveal protein expression at the single-cell level^28,33^ (**Fig. 1E**). Multiple pathologists reviewed the H&E and CyCIF images to identify and classify precursor lesions and cancer (**Fig. 1B, 1F**). Compared to samples with incidental STIC lesions (STIC.I), STIC co-existing with cancer samples (STIC.C) displayed a greater number of discrete STIC lesions (median: 5 STIC.C *vs* 2 STIC.I) (**Supplementary Fig. S1D**) and larger lesion size, indicated by a higher number of epithelial cells per lesion (**Supplementary Fig. S1E**). As expected, CyCIF analysis revealed elevated p53 protein levels in p53.I, STIC.I, STIC.C, and cancer, compared to normal epithelium (FT.I) (**Fig. 1F, 1G**), consistent with the role of mutant p53 in HGSOC development^34,35^. In addition, we observed a progressive increase in proliferation (Ki67+ cells) and DNA damage (ɣ-H2Ax+ cells) with disease progression (**Supplementary Fig. S1F-S1I**). Interestingly, STIC.I lesions exhibited significant variability in both proliferation and DNA damage, suggesting the potential for further subtyping of these incidental lesions based on their molecular characteristics (**Supplementary Fig. S1F, S1G**).

Spatial transcriptomic analysis (GeoMx) was performed on an adjacent tissue section to measure gene expression across the entire transcriptome, within specified tissue regions (**Fig. 1E, Supplementary Fig. S2A**). The H&E and CyCIF images were used to select regions of interest (ROIs) for GeoMx analysis, which included normal epithelium (FT and/or Fim), precursor epithelium, regions of the tumor, and stroma adjacent to most of the ROI (**Supplementary Fig. S2A-S2B**; see Methods; ROI annotation in **Supplementary File S2**). Of the 44 specimens, 35 had sufficient material for GeoMx analysis (see Methods). Principal component analysis (PCA) of this spatial transcriptomics data revealed that epithelial and stroma ROIs segregated along the first principal component (PC1), while the second principal component (PC2) distinguished between incidental precursors and cancer ROIs, reflecting their biological differences (**Supplementary Fig. S2C, Supplementary Fig. S3**).

### Molecular Transitions During HGSOC Development

HGSOC development is a multi-stage process driven by genetic and molecular alterations influenced by selective pressures^3,32^. To understand the transitions between the stages of HGSOC development, we analyzed the differential gene expression patterns within the epithelium across various stages of the disease using spatial transcriptomics data (**Fig. 2**). Gene set enrichment analysis (GSEA)^36,37^ revealed prominent activation of the IFN pathway (both IFN-α and IFN-ɣ) and cell cycle regulator pathways in the epithelium during early HGSOC development. This activation was evident comparing normal epithelium of incidental FT (FT.I) and incidental Fimbriae (Fim.I) to p53.I (**Supplementary Fig. S4A**) and p53.I to STIC.I (**Fig. 2A, 2B**).

**Figure 2:**
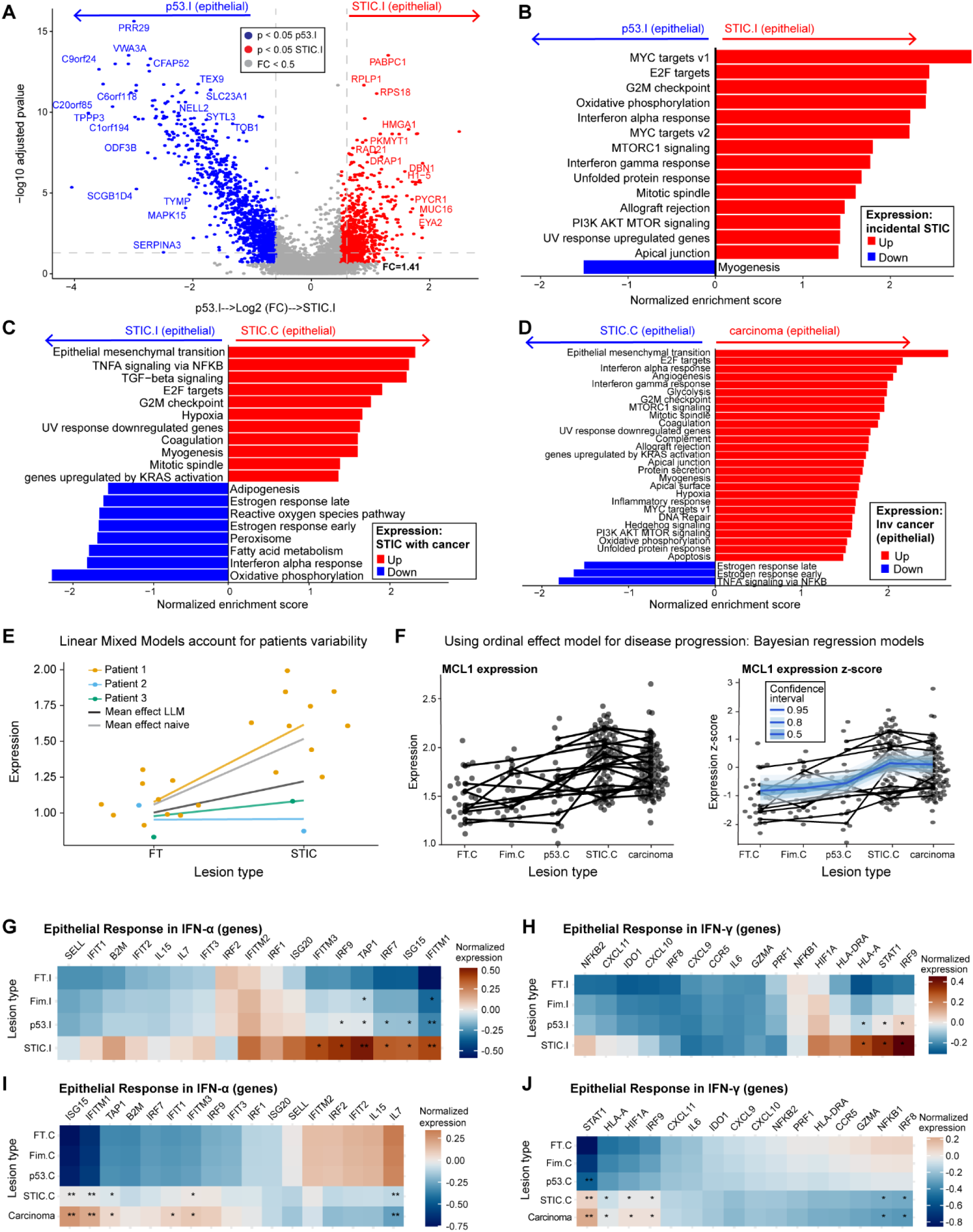
Molecular transitions during HGSOC development using spatial transcriptomics. **A.** Differential gene expression between epithelial segments of p53.I (n=39) and STIC.I (n=27). Linear Mixed Model (LMM) was applied for differential gene expression. LMM was performed with Benjamini-Hochberg (BH) correction using GeoMx DSP software (NanoString, v 3.1.0.221). Model formula: Lesions Type + (1| Scan_ID) whereby Scan_ID refers to the patient/slide ID. Only a subset of differentially expressed genes are shown. **B-D.** Gene Set Enrichment Analysis (GSEA) was performed based on the differential gene expression of the epithelia (as analyzed by GeoMx DSP software). MsigDB Cancer Hallmark gene sets associated with disease progression were compared between **(B)** p53.I (n=39) and STIC.I (n=27), **(C)** STIC.I (n=27) and STIC.C (n=96), and **(D)** STIC.C (n=96) and invasive carcinoma (n=105). Progression of STIC.I **(B)** was predominantly associated with interferon (IFN) and proliferative gene sets; STIC.C **(C)** was predominantly associated with EMT, TGF-β and hypoxia gene sets, and invasive carcinoma only events **(D)** were most likely to be associated with angiogenesis and Hedgehog signaling pathway. B-D. Pathway rankings were based on an adjusted p-value <0.05. **E-J.** Since LMMs and GSEA only allow us to look into pairwise comparison, we employed Bayesian regression modeling to model the progression from FT to cancer. Notably, Bayesian models differ from other methods since they take repeated sampling from the same patient into consideration. **E.** LMMs account for patient variability, demonstrated using a synthetic gene expression dataset across two disease stages. Repeated sampling from a single patient (Patient 1) introduces bias in estimating the mean effect. This bias is eliminated when patient variability is incorporated using LMMs. **F.** Bayesian ordinal regression models for gene expression across disease progression, exemplified using *MCL1*. While LMMs in the GeoMx DSP software allow only pairwise comparisons and a single random effect, Bayesian ordinal regression enables analysis across multiple disease stages. We implemented this approach using the "brms" R package, incorporating an ordinal monotonic constraint to model the stepwise sequence of lesions during disease progression^43^. GeoMx expression counts were Q3 normalized to account for sequencing depth (see Methods) and log transformed to stabilize variances. To account for differences in expression levels across different genes, the log-transformed values were further normalized by scaling to a mean of zero and variance of one (z-transform). We fitted one model per gene using the model specification gene_expression ∼ mo(stage) + (1 + mo(stage) | patient_id). A monotonic constraint was applied to enforce the assumption of an orderly sequence of these stages^43^. Repeat measurements from the same patients were accounted for by including patient-specific random intercepts and stage coefficients. In order to model the expression of gene sets, another random effect and its interaction with patient_id was included in the model (gene_expression ∼ mo(stage) + (1 + mo(stage) | patient_id * gene) (details are in **Methods**). **G-J.** In order to look into more details of IFN pathway and how it changes from normal FT to STIC.I or STIC.C to carcinoma, Bayesian ordinal regression model was applied to selected IFN-hallmark genes to compare expression changes in the epithelial and stromal compartments at different disease stages. Bayesian modeling allows to see the relative gene expression changes in the incidental or cancer group compared to the matched FT. Heatmaps show normalized expression of genes in the epithelia related to response in (G) IFN-α, (H) IFN-ɣ, from incidental group compared to their matched FT.I. Relative to the matched FT.I, upregulation of key genes induced by both IFN-α and IFN-ɣ activation, such as *STAT1, ISG15, IFITM1, IRF7, IRF9, HLA-A* at p53.I, indicating IFN activation at the very early stages of HGSOC progression. Heatmap showing normalized expression of genes in the epithelia related to response in (I) IFN-α, (J) IFN-ɣ, from cancer group compared to their matched FT.C. G-J. Columns correspond to individual genes, and rows correspond to types of lesions. Median of the posterior distribution was shown in heatmaps. Significance testing used the proportion of the 95% highest density interval (HDI) within the Region of Practical Equivalence (ROPE, 0.05 times the standard deviation). Comparisons with >95% of the HDI outside the ROPE were significant (*); >99% were very significant (**)^110^.

As the disease progressed, STIC.C lesions (regardless of BRCA status) displayed further enrichment of the IFN response as well as genes associated with TGF-β signaling, epithelial-to-mesenchymal transition (EMT), hypoxia, and tumor necrosis factor-α (TNF-α) signaling via NFκB pathway (**Fig. 2C, Supplementary Fig. S5)**. These pathways were further upregulated in established cancer (**Fig. 2D**), promoting invasion, motility, stress adaptation, and inflammation. The TGF-β pathway is known to induce EMT and interact with the PIK3K/mTOR or KRAS pathways^38^, both of which were activated before or during the STIC.C stage (**Fig. 2B, 2C**), potentially promoting growth even at early stages. Compared to precursor lesions, established cancer displayed enrichment for genes associated with hedgehog signaling and angiogenesis pathways (**Fig. 2D**). Analysis of the stromal compartment surrounding HGSOC precursors and cancer revealed a similar trend. In early stages, the stroma displayed enrichment for IFN response and IL6-JAK-STAT3 signaling pathways (**Supplementary Fig. S4B-S4E**), suggesting these pathways play a role during early stages of disease, likely through immune modulation and paracrine signaling.

### RNA Analysis Shows Early Persistent IFN-α/γ Activation, IRDS Emergence, and IFN-ε Signaling Downregulation in Later Stages

Since GSEA indicated early activation of the IFN response in HGSOC progression (starting at the p53.I stage), we performed a more detailed analysis of this pathway. We examined specific gene sets related to IFN-α, IFN-ɣ^36^, IFN-related DNA damage resistance signature (IRDS)^39,40^ and IFN-epsilon (ɛ)^41,42^ (**Supplementary File S3**). Due to the complex molecular relationships between disease stages, data variability, and patient-to-patient variation, we used a Bayesian ordinal regression model^43^ to analyze gene expression changes across disease stages (**Fig. 2E, 2F**).

The model confirmed a significant upregulation of key IFN-α/γ pathway genes in p53 signatures (p53.I), STIC lesions (STIC.I and STIC.C), and established cancer compared to matched normal epithelium (**Fig. 2G-2J**). This activation included significant upregulation of *STAT1* (an upstream activator of IFN signaling), and IFN-induced and regulatory factors (*IFITM1, IRF9, IRF7*), *with IFITM1* induced even in the Fim.I stage (**Fig. 2G, 2H**). IFN-stimulated genes (ISG; such as *ISG15*) were also elevated, along with genes involved in antigen processing and presentation (*TAP1, HLA-A*) (**Fig. 2G, 2H**). Prior studies have shown that elevated Type-I IFN activation results in the upregulation of classical major histocompatibility complex (MHC) genes, including Human Leukocyte Antigens (HLA) *HLA-A, HLA-B*, and *HLA-C.* The non-classical MHC-class gene HLA-E, often co-expressed with HLA-A upon IFN-ɣ activation^44–49^, plays additional immune regulatory roles. Upregulation of key IFN-α/γ pathway genes persisted in STIC.C and cancer cells (**Fig. 2I, 2J**). Furthermore, both classical (*HLA-A/B/C*) and non-classical (*HLA-E*) MHC-Class I antigen presentation molecules were upregulated in the epithelial compartment of p53.I and STIC.I lesions (**Supplementary Fig. S5B**). Collectively, these findings highlight a robust and sustained induction of IFN-α/γ pathway genes that persists throughout HGSOC development, underscoring a potential role in disease progression.

Recent studies have shown that the female reproductive tract epithelium, including the FT epithelium, constitutively expresses IFN-ε, a Type-I interferon, for immune defense^41,42,50,51^. Our data revealed a significant decrease in *IFNE* (i.e. IFN-ε transcript expression)*, IFNA2*, and *IFNA4* expression, in STIC.C and cancer epithelium compared to matched normal epithelium (**Supplementary Fig. S4F, S4G**). However, downregulation of IFN-ε signaling was not observed in incidental precursors, suggesting it occurs later in HGSOC progression, after the initial IFN-α and IFN-ɣ response.

Chronic activation of the IFN pathway, known as IRDS, has been linked to chemotherapy and radiotherapy resistance in various cancers^39,40,52^. Upregulated IRDS genes included *STAT1*, *MX-1*, and the anti-apoptotic BCl-2 family member *MCL-1* in STIC.C and cancer, with a trend toward increased expression in STIC.I (**Supplementary Fig. S4H, S4I**). Persistent IFN activation might contribute to the emergence of IRDS, particularly during the later stages of STIC clonal expansion. Overall, the data suggests dynamic shifts in IFN signaling throughout HGSOC progression. Early activation of IFN during the p53 signature stage persists and intensifies as the disease progresses, coinciding with the downregulation of IFN-ε signaling and the emergence of IRDS. These findings indicate a potential link between sustained IFN-α/ɣ activation, clonal selection, and the gradual accumulation of tumor-promoting and immune-suppressive pathways.

### Multiplexed Imaging Reveals Spatially Coordinated IFN Signaling in HGSOC Progression

CyCIF is a multiplexed imaging technique that quantifies protein marker expression at a single-cell level^53^. We used CyCIF to visualize IFN pathway activity within HGSOC precursors. This analysis revealed a stepwise increase in the number of epithelial cells expressing IFN pathway markers (e.g., p-TBK1, p-STATs, HLA-A/E) across disease stages (p53.I: median 22%, STIC.I: 33%, STIC.C: 43%, and cancer: 26%), regardless of BRCA mutation status (**Fig. 3A-3C, Supplementary Fig. S6-S8**). High-resolution 3D CyCIF imaging confirmed co-expression of the IFN inducible protein, MX-1, which formed multiple discrete puncta, and the antigen-presenting protein HLA-A within epithelial cells (PanCK+) in STIC.C and invasive cancer (**Fig. 3D, 3E**), reconfirming IFN induction in MHC-class I + epithelial cells.

**Figure 3:**
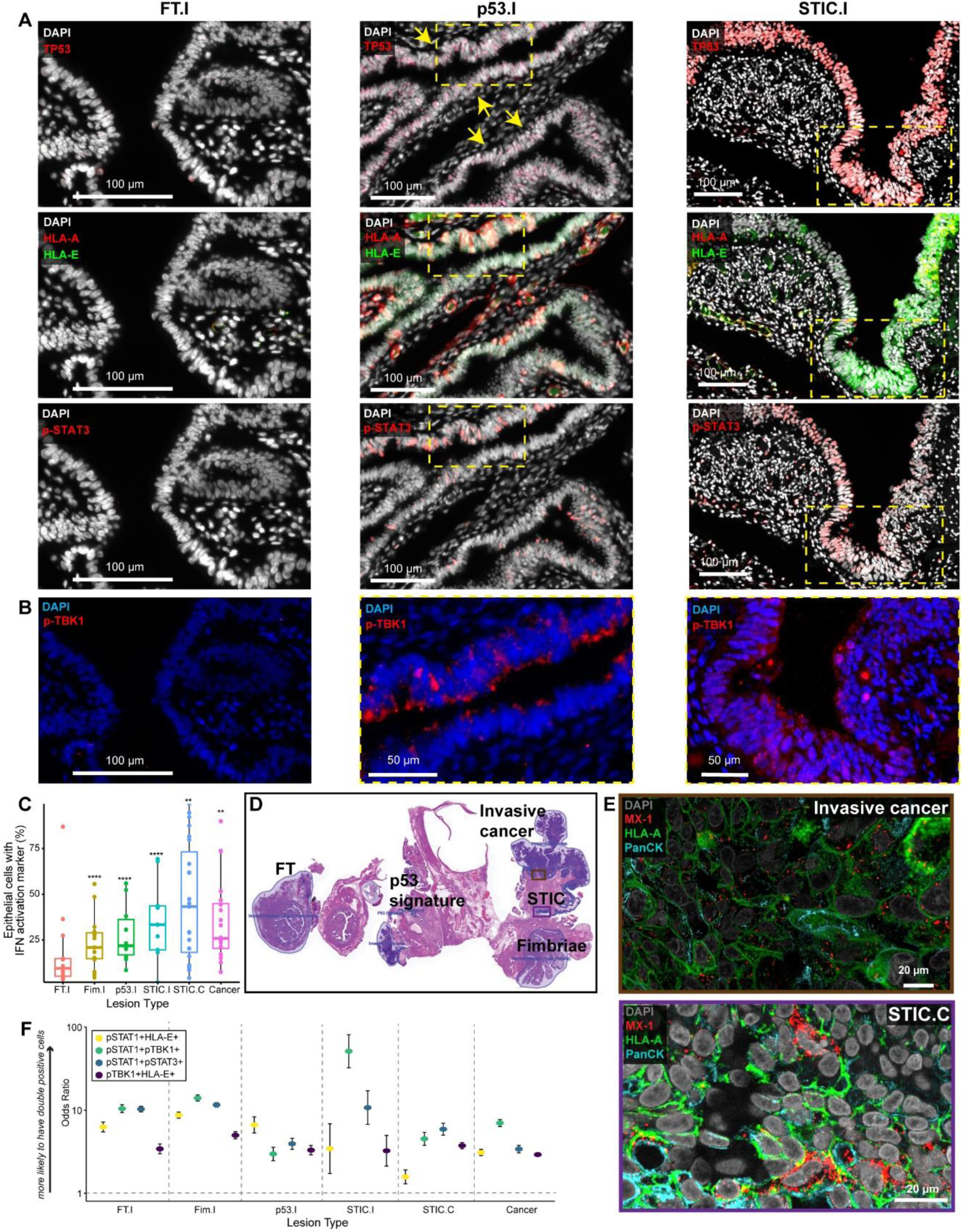
Multiplexed tissue imaging revealed spatially coordinated IFN signaling in HGSOC progression. **A.** CyCIF images showing markers related to downstream of IFN signaling pathway activation, such as overexpression of MHC-Class I (HLA-A and HLA-E) and p-STAT3 in both p53..I The representative STIC.I case is shown with matched FT.I (case CD302.04(939), patient ID 40, BRCA WT). The p53.I sample is from a different patient (case C21-22 patient ID 28, BRCA1 Mut). Yellow arrows on p53.I indicate the layer of epithelial cells representing “p53signatures”. **B.** CyCIF images showing p-TBK1 (cytosolic/ *punctate*) expression in regions that express HLA-A, HLA-E and p-STAT3 (FT.I identical ROI to **A**, p53.I and STIC.I ROIs outlined with yellow box in **A**), suggestive of one of the potential upstream mechanisms of IFN pathway activation. **C.** Box plot showing the percentage of epithelial cells with an IFN activation marker (p-TBK1+/p-STAT1+/HLA-E+/p-STAT3+). Quantification is based on single-cell quantification from tissue imaging data and suggests an increased number of cells with IFN activation marker with disease progression. The number of specimens for each lesion as follows: FT.I (n=13), Fim.I (n=15), p53.I (n=10), STIC.I (n=9), STIC.C (n=23), and Cancer (n=20). The solid line indicates the median within the interquartile range, with whiskers extending to a maximum of 1.5 times the interquartile range beyond the box. Black asterisks indicate significant differences in stages compared to the FT.I; **p<0.001, ****p<0.0001. Binomial Generalized Linear Mixed Models (GLMMs) taking patient ID as random effect. GLMMs was implemented in the lme4 R package (v 4.3.3). For each ROI the number of cells with the given phenotype (“successes”) and of all other phenotypes (“failures”) was modelled using the binomial distribution with a logit link function using the lme4 model formula cbind(n_success, n_failure) ∼ stage + (1 + stage | patient_id). See **Supplementary File S5, S6** for summary statistics. **D.** H&E image of WSI (Case RD-23-002, patient ID 9, BRCA2 mutant, Stage IC HGSOC) with ROIs for **E** indicated. **E.** 3D reconstruction using 3D CyCIF imaging of a case of STIC with concurrent HGSOC (patient ID 9, ROIs from **D**). MX-1, a well-known IFN-induced gene, shows *punctate* expression and is co-expressed with PanCK+ HLA-A+, suggesting IFN activation in both tumor and STIC.C epithelium. **F.** Plot of odds ratio (OR) for protein pairs (p-STAT1+ HLA-E+ or p-STAT1+ p-TBK1+ or p-STAT1+ p-STAT3+ or p-TBK1+ HLA-E+), showing the likelihood of co-expression of these marker pairs in epithelial cells across disease stages. Odds ratio (OR) with confidence interval (CI) (Y-axis) is >1, where OR<1 indicates a lower likelihood of co-expression. Y axis is presented in log(10) scale. After gating single cells (+ or – for a marker), a contingency table was generated with the total number of positive or negative cells for two markers of interest (such as p-STAT1 and HLA-E) (using R v 4.3.3). Next, GLMMs were performed to calculate the OR with CI and significance testing. All lesion types showed significance of having double positive cells (p<0.001). Fisher exact test was also performed, suggesting a similar OR to GLMMs. Summary statistics are in **Supplementary Files S5 and S6**.

Despite the observed increase in IFN marker expression with disease progression, there was significant inter-and intra-sample heterogeneity in the presence or absence of IFN pathway markers across all stages (**Supplementary Fig. S2A, S7, S9-S11**). For instance, within the same STIC.I lesion, we observed both HLA-E positive and HLA-E negative cells (**Supplementary Fig. S2A**). In cancer, we also noted variability in the intensity of IFN pathway marker expression, such as the presence of both HLA-E low and high-intensity tumor regions within the same patient (**Supplementary Fig. S10**). We leveraged this heterogeneity to explore the potential for localized and coordinated IFN signaling in HGSOC precursors. Visual inspection of CyCIF images suggested co-expression of IFN markers within individual cells (**Fig. 3A, 3B, 3E, Supplementary Fig. S6, S9**). This observation was further supported by pairwise correlation analysis, which demonstrated strong positive correlation between various pairs of IFN markers (e.g., pSTAT1+ HLA-E+; p-STAT1+ p-TBK1+; pSTAT1+ p-STAT3+; p-TBK1+ HLA-E+) across all disease stages (Odds Ratio (OR) 1.5-50, **Fig. 3F**). While the percentage of cells expressing IFN markers (e.g., HLA-E) was relatively low at early stages (p53.I median: 4%), it increased in STIC.I (16%), STIC.C (26%), and invasive cancer (18%) (**Supplementary Fig. S8C**). The high probability of IFN marker co-expression suggests coordinated and localized activation of the IFN pathway throughout HGSOC development. The progressive increase in IFN-positive cells within precursor lesions further implies clonal expansion driven by positive selection during disease progression.

### Interplay Between HLA-E Expression, IFN Pathway, and Immune Evasion Mechanisms

Loss of classical MHC-class I is a well-established mechanism of immune evasion in cancers, including advanced HGSOC^54,55^. However, tumors may become vulnerable to NK cell-mediated killing when non-classical MHC-class I HLA-E is lost^45,56,57^. Conversely, overexpression of HLA-E in various cancers, including cervical and ovarian cancer, allows tumor cells to evade NK cell surveillance^45,47,58–60^, likely contributing to the development of more aggressive tumor^48,58–60^. To investigate the relationship between HLA-E expression and transcriptional programs, we categorized the GeoMx ROIs based on HLA-E protein expression (imaged from serial tissue sections) into HLA-E positive or HLA-E negative groups (**Supplementary Fig. S2A, Supplementary File S2**). As expected, analysis of differentially expressed genes showed a significant upregulation of the *HLA-E* gene itself in ROIs positive for HLA-E protein (e.g., from STIC.I, **Supplementary Fig. S12A**). Interestingly, in early STIC lesions (STIC.I), HLA-E positive regions showed enriched expression of genes related to cell morphology and migration (*ARAP2*), antigen presentation (*HLA-A, HLA-B, HLA-DRA, HLA-DQB1, HLA-F, TAP, TAPBP*), coagulation (*F5*), and complement pathways (*CFB*, *CFH*, *CFI*, *C1S*, *C2*, *C4B*), suggesting a coordinated inflammatory and tissue remodeling response. Cell growth and maturation pathways were also induced, including the *MYC* oncogene, as were downstream IFN genes (*MX1*, *IRF1, DDX60*, *OAS3*, *IFI44*, *PSMB8*, *XAF1)*. Further analysis using Bayesian modeling with HLA-E status as covariate (**Supplementary Fig. S13**) showed a strong association of HLA-E positive epithelial ROIs with IFN responses (Type I and II; IFN-α/ɣ) and IRDS, even as early as the p53.I stage (**Supplementary Fig. S12B).** Differential gene expression and GSEA analysis further supported this association across disease stages (**Supplementary Fig. S12C-S12D**, **S14A-14D**). These findings suggest a complex interplay between HLA-E expression, IFN signaling and various immune evasion mechanisms, potentially supporting early tumor development and progression.

In contrast to IFN pathway activation, genes associated with other well-established tumor-promoting pathways, such as TGF-β, EMT, and those characteristic of the aggressive C5/PRO subtype of HGSOC^22,23^ (**Supplementary File S3**), emerged primarily at the STIC.C and cancer stages (**Supplementary Fig. S14E, S14F)**. Notably, these pathways were not restricted to HLA-E-positive epithelial areas. Further analysis of individual EMT-related (e.g., *CLDN6, CDH3, COL4A1, MMP14*, *MMP2*) confirmed that increased expression at later stages was not limited to HLA-E positive cells (**Supplementary Fig. S15**). This suggests that additional tumor-promoting mechanisms, independent of the IFN pathway, likely play important roles in shaping the HGSOC development.

### Micronuclear Rupture and cGAS Recruitment in HGSOC Progression

The presence of an IFN response and p-TBK1+ epithelial cells in early HGSOC precursors suggests activation of the cGAS-STING signaling pathway^61–63^. This pathway responds to cytosolic DNA from various sources, including DNA damage or CIN^62,64–66^. CIN, a hallmark of advanced HGSOC^3,67^, can lead to the formation and rupture of micronuclei (MN) containing mis-segregated chromosomes^63,68,69^. We used CyCIF to identify p53+ precursor lesions and assessed the presence and integrity of MN using barrier-to-autointegration-factor (BAF; *BANF1)*, a sensitive marker for cytosolic DNA^70,71^. Strikingly, BAF+ MN ruptures were observed as early as STIC.I lesions, with increasing frequency in invasive cancer. Visual inspection revealed that a subset of ruptured MN contained cGAS, and some also contained the DNA damage marker ɣ-H2Ax (**Fig. 4A, 4B, Supplementary Fig. S16**). High-resolution 3D CyCIF^29^ confirmed the co-localization of BAF, cGAS, and ɣ-H2Ax, with the highest frequency observed in invasive HGSOC (**Fig. 4C, 4D, Supplementary Video**; **Supplementary Fig. S16**). These findings suggest that CIN-induced MN ruptures occur unexpectedly early in HGSOC progression, potentially triggering IFN signaling via activation of the cGAS-STING pathway.

**Figure 4:**
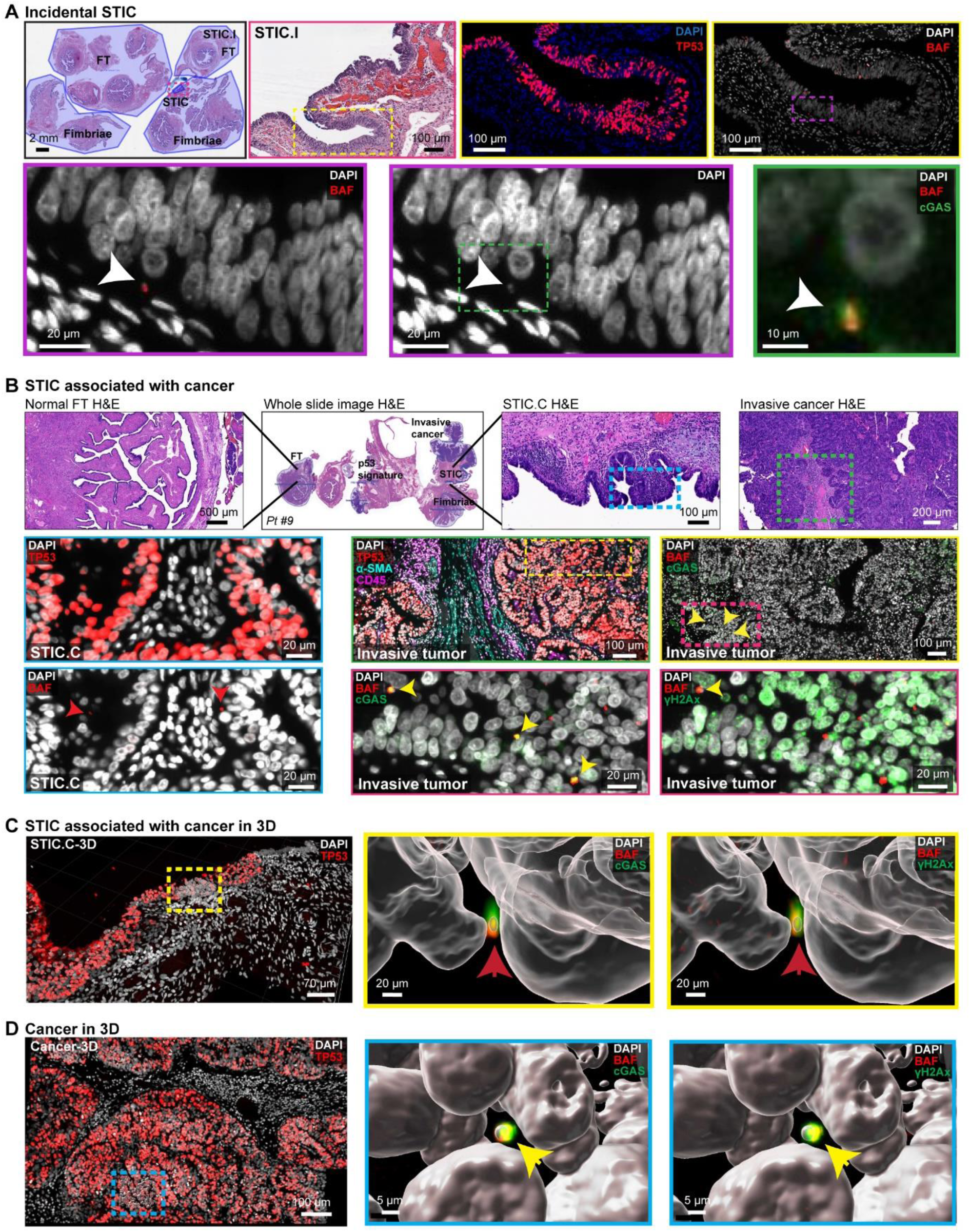
Tissue imaging revealed Micronuclear Rupture and cGAS Recruitment in HGSOC Progression. **A.** Top row: H&E of a representative case of STIC.I (case CD302.03(706), patient ID 38, BRCA1 Mut), with ROIs indicated. CyCIF imaging confirms TP53+ epithelial cells on STIC containing BAF positive staining, outlined with a purple box. Bottom: Co-localization of BAF+ staining with DNA (DAPI) indicates ruptured micronuclei (MN) (white arrowhead). Higher magnification panel (outlined with a green box) confirms cGAS colocalization with BAF+ MN and indicates cGAS binding to the ruptured micronuclear DNA. **B.** Top: H&E of a representative case of STIC with concurrent HGSOC, with ROIs for different histologies shown (also shown in Figure 1; Case RD-23-002, patient ID 9, BRCA2 mutant, Stage IC HGSOC). Cyan box on STIC.C H&E and brown box on invasive tumor H&E indicated ROI for panels in lower rows. Lower rows: CyCIF images of STIC.C (left column) showing BAF+ MN (red arrow heads). Additional CyCIF images of invasive cancer (right columns), showing a sharp increase of BAF+cGAS+ or BAF+ ɣ-H2Ax+ MN in the invasive component (yellow arrow heads). **C-D.** The same specimen as shown in panel **B** was imaged using 3D confocal microscopy to confirm the intact nuclei and MN rupture event(s). 3D reconstruction and surface rendering of the 20 µm thick specimen confirm the colocalization of BAF+ and cGAS+ MN as well as co-expression of BAF+ ɣ-H2Ax+ MN rupture events in STIC.C (**C**; red arrowheads outlined with a yellow box) and in the invasive cancer component (**D**; yellow arrowheads outlined with a green box). Co-localization of cGAS and BAF+ MN potentially indicates the sensing of MN-ruptured DNA by cGAS and activating the cGAS-STING pathway.

### Spatial Organization of Immune Cells in HGSOC Development

Activation of the cGAS-STING signaling pathway and the resulting IFN response play critical roles in shaping the spatial organization and function of the immune system within the TME^62,72^. To investigate how the spatial organization of immune cells changes during HGSOC development, we first used CyCIF data to quantify major immune cell types in precursor lesions and invasive cancer (**Fig. 5A-5C**). These cell types included: i) antigen-presenting cells (APC), such as conventional dendritic cells Type-1 (cDC1) and macrophage-derived APCs, ii) CD68+ macrophages (M1-like), iii) CD163+ macrophages (M2-like), iv) CD20+ B cells, v) CD4+ T cells, vi) CD8+ T lymphocytes (**Fig. 5A-5E**). A minor population of NK cells was also detected (**Supplementary Fig. S17A-S17F**).

**Figure 5:**
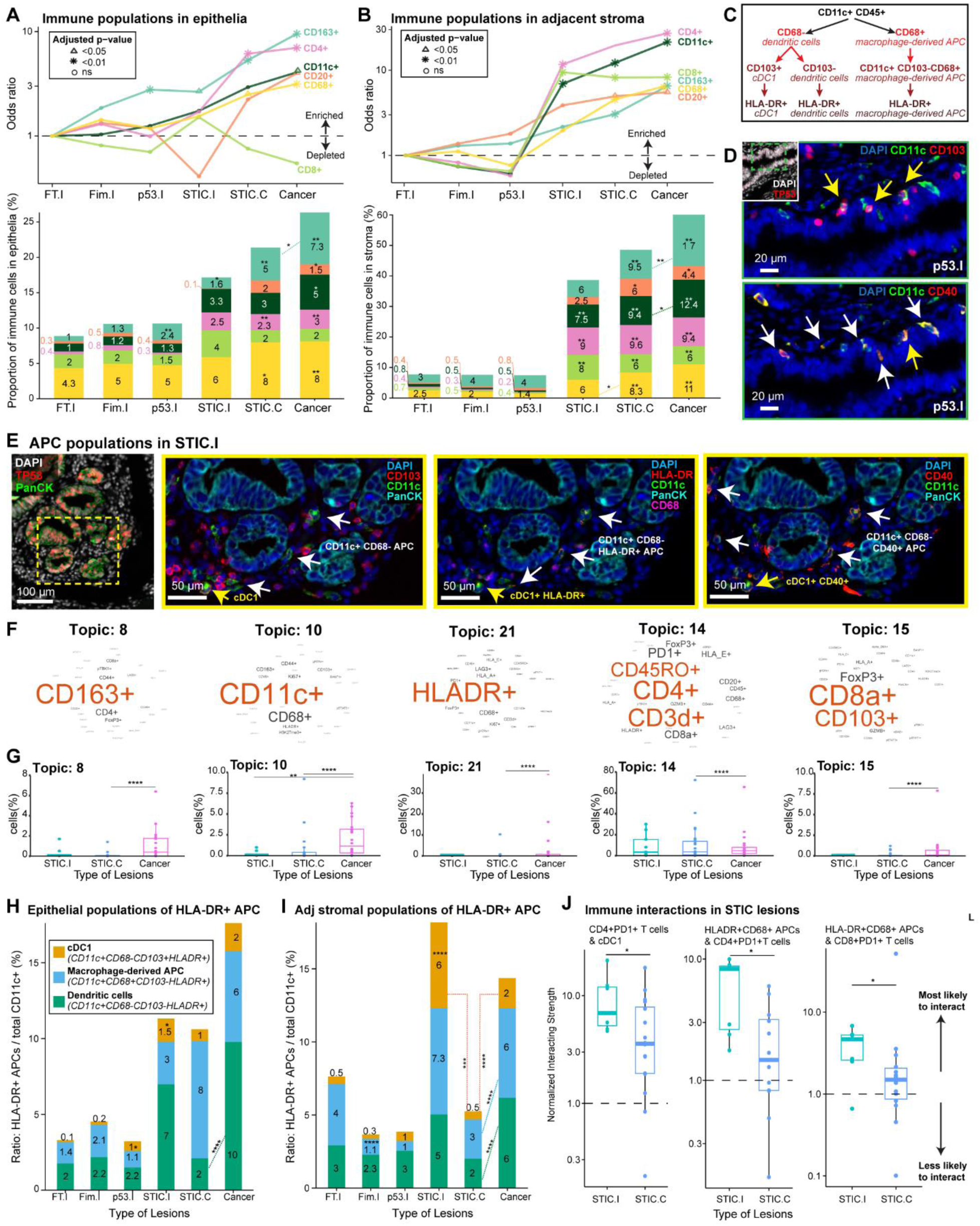
Immune composition suggests an active immune surveillance by the presence of activated antigen presenting cells (APC) at the early HGSOC progression. **A-B.** Top: Line plots depicting changes in the proportions of major immune cell types across disease stages. Data are derived from single-cell CyCIF analysis of (A) epithelial tissue and (B) adjacent stroma. The y-axis shows odds ratios relative to the proportions in FT.I on a log scale. Odds ratios and p-values were derived from binomial GLMMs, with patient ID and observation-level random effects. p-values were adjusted for multiple testing using the Benjamini–Hochberg (BH) procedure. Bottom: Stacked bar plots showing the average proportion of major immune cell types derived from single cell-CyCIF analysis across disease stages in epithelium **(A)** and in adjacent stroma **(B)**. Asterisks on bars indicate significant differences compared to the FT.I stage; asterisks between bars (with dashed lines) indicate significant comparisons between stages. *p<0.05, **p<0.01. Binomial GLMMs, with patient ID and observational level random effect. Average proportions were rounded up to the next whole number when applicable and shown for each cell type across lesion types. Number of specimens per group as follows: FT.I (n=13), Fim.I (n=15), p53.I (n=10), STIC.I (n=9), STIC.C (n=23), and Cancer (n=20). **C.** A schematic showing different subsets of CD11c+ population identified by tissue imaging and their definition within this manuscript. cDC1: conventional dendritic cells, APC: antigen-presenting cells. HLA-DR+ populations represent an activated APC state; the first step of activation of any antigen-presenting cells is expressing MHC-class II (HLA-DR). **D.** CyCIF image showing from a representative case of a p53.I, also shown in Figure 3 (case C21-22 patient ID 28, BRCA1 Mut). The green ROI on the inset indicates the layer of epithelial cells representing “p53signatures”. The epithelium of the “signatures” showed the presence of activated APCs, such as activated cDC1 and other CD11c+ dendritic cells. CD40+ CD11c+ indicates the presence of a co-stimulatory molecule, meaning activated dendritic cells, both cDC1 (shown with yellow arrows) and other dendritic cells (shown with white arrows). **E.** CyCIF image showing a representative case of a STIC.I, also shown in Figure 4 (STIC.I, case CD302.03(706), patient ID 38, BRCA1 Mut). An area outlined with a yellow box shows the presence of the overall APC population, including cDC1. Furthermore, these APCs are “activated” (i.e. presenting antigens (HLA-DR+)) or CD40+). The yellow arrow indicates cDC1, and the white arrows indicate other dendritic cells, excluding cDC1, in the adjacent stroma, close to the STIC.I epithelium. Overall, the presence of activated APCs, either dendritic cells (CD11c+ CD68-) or macrophage derived (CD11c+ CD68+) might be indicative of active immune surveillance at the early precursor lesions. **F.** Latent Dirichlet Allocation (LDA) neighborhood analysis for spatial topic analysis was performed from multiplex tissue imaging. Some topics indicating of cellular neighborhoods related to immune cell populations and states. The pooled frequencies of all samples, both incidental and cancer group were used to train the final LDA model. **G.** Box plots depicting the percentage of cells in each topic (described in F)across lesion stages (STIC.I, STIC.C, and Cancer). The number of specimens for each lesion as follows: STIC.I (n=9), STIC.C (n=23), and Cancer (n=20). The solid line indicates the median within the interquartile range, with whiskers extending to a maximum of 1.5 times the interquartile range beyond the box. Black asterisks indicate significant differences between groups; ****p<0.0001, ** p<0.01, using GLMMs and taking patient ID as random effect. **H.** Average proportion for each HLA-DR+ APC subset (as a fraction of the total CD11c+ population) across lesion types shown as a stacked bar plot, in the epithelia. **I.** Average proportion for each HLA-DR+ APC subset (as a fraction of the total CD11c+ population) across lesion types shown as a stacked bar plot, in the adjacent stroma. **H-I.** Average proportions for each subset across lesion types were shown, rounded to the nearest whole number when applicable. Number of specimens per group as follows: FT.I (n=13), Fim.I (n=15), p53.I (n=10), STIC.I (n=9), STIC.C (n=23), and Cancer (n=20). Asterisks on bars indicate significant differences in cell proportions compared to the FT.I stage; asterisks between bars (dashed lines) indicate significant differences between groups. *p<0.05, **p<0.01, ****p<0.0001, binomial GLMMs, taking patient ID as random effect. Average proportions were rounded up to the next whole number when applicable and shown for each cell type across lesion types. **J.** Box plots comparing normalized interaction strength between different cell types at STIC.I and STIC.C stages: cDC1and CD4+ PD1+ T cells (left), HLADR+ CD68+ APCs and CD4+ CD8+ PD1+ T cells (middle), and HLADR+ APCs and CD8+ PD1+ T cells (right). Scores >1 indicate more and <1 indicates fewer interactions between the two cell types in STIC.C compared to STIC.I than expected by chance. The interaction score was normalized against the distance of random sampling. Since at least five cells of both populations will have to be present for random sampling, the number of specimens analyzed from tissue imaging was STIC.I (n=7), STIC.C (n=16) (p=0.03) (cDC1 & activated CD4+); STIC.I (n=7), STIC.C (n=12) (p=0.02) (HLA-DR+ CD68+ APCs & activated CD4+) and STIC.I (n=7), STIC.C (n=14) (p=0.04) (HLA-DR+CD68+ & activated CD8+). Y axis is presented in log(10) scale. The solid line indicates the median within the interquartile range, with whiskers extending to a maximum of 1.5 times the interquartile range beyond the box. Black asterisks indicate significant differences between groups; *<0.05, Wilcoxon rank sum test.

We then applied latent Dirichlet allocation (LDA), a statistical method for topic modeling, to analyze the spatial patterns of the immune microenvironment. LDA has been successfully used to reveal recurrent cellular neighborhoods in both precancer and cancer tissues^53,73,74^. Based on cell type and activation markers, 4.22×10^7^ single cells from 44 specimens were classified into 21 distinct groups, revealing recurrent local cellular neighborhoods (‘topics’). These topics represent niches of specialized cell types or interacting cell types that may play a role in disease progression or response to therapy. Several notable recurrent neighborhoods were identified, including Topic 8, which predominantly comprised CD163+ (M2-like) macrophages, Topic 10, dominated by CD11c+ APCs, and Topic 21, characterized by HLA-DR+ (activated) APC cells (**Fig. 5F**). Each of these neighborhoods was significantly enriched in invasive cancer compared to STIC lesions (**Fig. 5G**). Topic 14 displayed high levels of CD4+ T cells, while Topic 15 contained both CD8+ T lymphocytes (including CD103+ T_RM_ cells) and FoxP3 T regulatory cells (Tregs). These T-cell rich neighborhoods were also more prevalent in invasive cancer compared to STIC lesions (**Fig. 5F, 5G**). These findings suggest that HGSOC progression involves dynamic reorganization of immune cell spatial distribution, potentially influencing cell type interactions, immune signaling, and the overall anti-tumor immune response. We next sought to use our CyCIF and GeoMx datasets to better understand how the organization of specific immune cell subtypes changes during different stages of HGSOC progression.

### cDC1 and NK Cells Decrease while Macrophages Increase in Later Stages of HGSOC Development

We first focused on two cell types that are critical organizers of the initial immune response to tumors: cDC1 (CD11c+, CD103+, CD68-)^56,75,76^ and NK cells (NKG2D+ CD3-). Intra-tumoral NK cells produce chemokines that recruit cDC1 into the TME, while cDC1 cells transport tumor antigens to lymph nodes to promote cytotoxic T lymphocyte (CTL) responses. cDC1 cells have superior antigen processing and presentation capacities, making them highly effective in activating and recruiting CD8+ CTLs^75,77–80^. Using CyCIF imaging, we observed that although the total population of CD11c+ APCs (which includes cDC1 cells) increased with disease progression (**Fig. 5A, 5B**), the number of HLA-DR+ cDC1 cells (indicating antigen presenting function) rose significantly in the early stages, particularly in the epithelium (**Fig. 5C, 5H)**. These cells were 10-fold higher in p53.I epithelium and 15-fold higher in STIC.I epithelium compared to normal epithelium but remained relatively stable with disease progression. In the stroma, cDC1 populations expressing HLA-DR increased 12-fold in STIC.I compared to normal tissues but decreased substantially in more advanced lesions (12-fold decrease in STIC.C and 3-fold in established cancers compared to STIC.I) (**Fig. 5I**). This suggests that the antigen-presenting function provided by cDC1 decreases as precursor lesions progress. Similarly, NK cells, which were present at low levels in normal tissue and p53.I precursors (median 0.1%), became nearly undetectable in later stages of the disease (STIC.I, STIC.C, and cancer; median: 0.02%) (**Supplementary Fig. S17**), indicating a further decline in anti-tumor immunity.

To validate these observations, we performed Bayesian modeling of the spatial transcriptomics data to analyze gene sets specific to cDC1 and NK cells^56,75,76^ (**Supplementary File S3**). This analysis confirmed the CyCIF findings, showing that cDC1 and NK cells are present in early precursors but decrease with disease progression, particularly in STIC.C and tumor stages. Later-stage lesions showed reduced gene expression related to cDC1 function and the NK-cDC1 axis activity. These genes included *CLEC9A, BATF3, CLNK, XCL1, XCR1*, *IL-15,* and *IL-12*. In addition, genes associated with NK cell receptors, such as *KLRK1, NCR1, CD226* (which encode NKG2D, and activation receptors NKp46 and DNAM1, respectively), also declined. There was also a decrease in genes related to NK inhibitory receptors, including *KLRD1* (CD94), *KLRC2* (NKG2A/C), and *KLRG1,* as well as *KIR*s (**Supplementary Fig. S18A, S18B**). This data indicates progressive dysfunction of NK and T cells in STIC.C and tumor epithelium, supported by decreased expression of genes related to cytotoxic activity, such as *PRF1, NKG7, GZMB, GZMK, GZMH,* and *GZMA* (**Supplementary Fig. S18C, S18D**). These findings suggest a suppression of NK cell and cDC1 activity during HGSOC development.

While NK cells and cDC1 decline during HGSOC development, tissue imaging revealed a stepwise increase in macrophage populations with disease progression, as suggested by LDA analysis Topic 8 (**Fig. 5F-G**). Both CD68+ M1-like macrophages and CD163+ M2-like macrophages became more abundant, reaching their highest levels in invasive cancer (**Fig. 5A, 5B, Supplementary Fig. S19A, 19B**).

Interestingly, more than 50% of CD68+ cells co-expressed CD11c, suggesting they function as APCs (i.e., macrophage-derived APCs^78^) (**Supplementary Fig. S19C, S19D**). These macrophage-derived APCs frequently co-expressed HLA-DR^81^ and/or CD40 which are markers of antigen presentation and T cell co-stimulation (**Fig. 5D, 5E, 5H, 5I**, **Supplementary Fig. S19**). For comparison, less than 1% of epithelial cells in precursor lesions or tumor expressed HLA-DR (**Supplementary Fig. S20**). Spatial transcriptomics data further supported the increase in APCs with disease progression. While multiple cell types can produce various chemokines and cytokines (e.g., CXCL9, CCL3 and TNF), the data showed an increasing trend of gene expression associated with both M1 and M2 macrophages^82^ in the cancer group, including stabilin-1 (*STAB1)*, *AXL*, *IL-10* and *CD163* (**Supplementary Fig. S21**). Moreover, there was an increase in the expression of MHC-class II (*HLA-DRA, HLA-DMA, and HLA-DRB1*) in the stroma of precursors and invasive cancer (**Supplementary Fig. S18E, S18F**). Taken together, these findings suggest an increase in macrophage-derived APCs that express HLA-DR, especially in cancer, indicating that these cells may play a role in presenting tumor antigens to CD4+ T cells.

### Shifting Functional Landscape of CD4+ T Cells in HGSOC Development

The observed changes in APC composition during HGSOC progression – a decrease in cDC1 and an increase in macrophage-derived APCs – led us to characterize CD4+ T cells, critical partners with APCs in orchestrating adaptive immune responses. CyCIF analysis revealed a progressive increase in CD4+ T cell infiltration throughout the disease course (**Fig. 5A, 5B, Supplementary Fig. S22A, S22B**). However, both activation and dysfunction markers were observed in these CD4+ T cells. Over 35% of CD4+ T cells expressed either HLA-DR or PD1, suggesting potential antigen presentation and activation, starting from the STIC.I stage (**Supplementary Fig. S22**). At the same time, the presence of numerous regulatory T cells (CD4+ FoxP3+ Tregs: ∼14%) and CD4+ LAG3+ (∼14%) indicated the development of an immunosuppressive microenvironment and potential dysfunction of antigen presentation (**Supplementary Fig. S22A, S22B**).

CD4+ T cell activation relies on antigen presentation by MHC-class II molecules, primarily expressed by APCs. While direct interactions between CD4+ T cells, HLA-DR expressing APCs, and CD8 T cells were observed, there was evidence of a dampening effect – some APCs co-expressed HLA-DR+/CD40+ along with TIM3, an inhibitory receptor that modulates the function of both lymphoid cells and APCs, suggesting a reduced capacity for antigen presentation^83–86^ (**Supplementary Fig. S23**). Quantitative analysis comparing the cell-cell interactions in STIC.I and STIC.C lesions showed decreased proximity (normalized interaction score) between activated CD4+ T cells, CD8+ T cells, and cDC1/macrophage-derived APCs expressing HLA-DR in STIC.C lesions (**Fig. 5J**). Thus, while CD4 T cells infiltrate HGSOC precursor lesions, their capacity for activation appears to be counterbalanced by the emergence of suppressive mechanisms, potentially contributing to disease progression.

### CD8 T cell Dynamics and Dysfunction in Early HGSOC Development

We also observed changes in the numbers, activation states, and localization of CD8+ T cells within the tissues during HGSOC development. Compared to normal tissue, CD8+ T cells increased significantly in the epithelium of STIC.I lesions (2-fold) and even more dramatically in the stroma (10-fold) (**Fig. 5A, 5B**). However, this increase was followed by a gradual decline in later disease stages (STIC.C and cancer), with a sharper decrease in the epithelium compared to the stroma (2-fold vs. 1.5-fold).

Many different CD8+ T cell subsets were present, including tissue-resident memory T cells (T_RM_; CD8+ CD103+ CD45RO+), a specialized subset of CD8+ T cells adapted for localized immune surveillance within specific tissues^87,88^, and conventional cytotoxic CD8+ T lymphocytes (CTL: CD8+ CD103-CD45RO-). We used markers such as Ki67, PD1, GMZB, TIM3, and LAG3 to assess their functional states (**Fig. 6A-6D**). T_RM_ cells increased 1.5-fold in p53.I epithelium compared to normal epithelium, but gradually decreased with disease progression (**Fig. 6A, 6E, 6F**). CTL numbers started to rise in p53.I, with the most striking increase observed in STIC.I epithelium and stroma, followed by a gradual decrease in STIC.C (4-fold) and cancer (2-fold), particularly in the epithelium (**Fig. 6A, 6B**).

**Figure 6:**
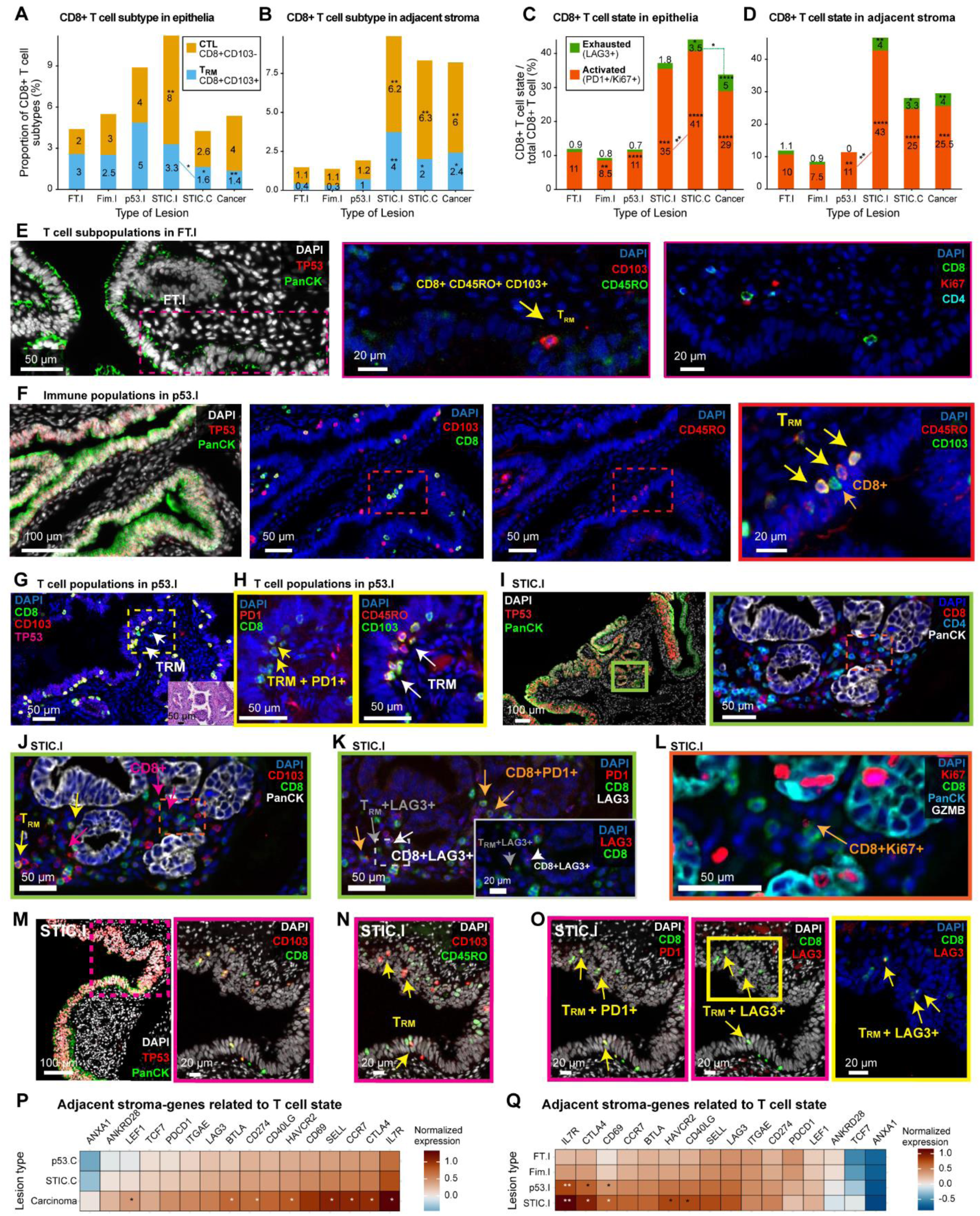
Immune editing and T cell dysfunction at early stage of HGSOC development. **A-B.** Stacked bar plots showing summary of major T cell subtypes from single cell-CyCIF analysis in the epithelia (A) and in the adjacent stroma (B) across disease stages. A-B. The average proportion of T cell subtypes shown is CD8+ CD103+ T cells, indicating tissue-resident memory T cells (also express CD45RO) (T_RM_) as well as CD8+ CD103-T (Cytotoxic T: CTL) cells. Number of specimens per group as follows: FT.I (n=13), Fim.I (n=15), p53.I (n=10), STIC.I (n=9), STIC.C (n=23), and Cancer (n=20). Black asterisks indicate significant differences in stages compared to the FT.I; *p<0.05, **p<0.01. Binomial GLMMs, with patient ID and observational level random effect. Colored asterisks with dashed lines were also shown based on GLMM output, only when significant comparing between groups (for eg. STIC.I *vs* STIC.C). **C-D.** Stacked bar plot showing the proportion of CD8+ T cell states in total CD8+ T cells from CyCIF analysis in the epithelia (C) or the adjacent stroma (D). T cell states are defined as follows: activated CD8+ T cells: Ki67/PD1+ and LAG3- and exhausted CD8+ T cells: PD1+LAG3+/LAG3+. Number of specimens per group as follows: FT.I (n=13), Fim.I (n=15), p53.I (n=10), STIC.I (n=9), STIC.C (n=23), and Cancer (n=20). Black asterisks indicate significant differences in stages compared to the FT.I; *p<0.05, ***p<0.001, ****p<0.0001, binomial GLMMs. Colored asterisks with dashed lines were also shown based on GLMM output, only when significant comparing between groups (for eg. STIC.I *vs* STIC.C or p53.I *vs* STIC.I). **E.** CyCIF image showing from a representative FT.I from a matched STIC.I, also shown in Figure 3 (case CD302.04(939), patient ID 40, BRCA WT). The Yellow arrows indicate a T_RM_ (CD103+ CD45RO+ CD8+). **F.** CyCIF image showing from a representative p53.I, also shown in Figure 3 (case C21-22 patient ID 28, BRCA1 Mut). Red box indicates the layer of epithelial cells representing a “p53 signatures”, which show the presence of T_RM_ (yellow arrows) and CD8+ 103-T (CTL) (orange arrows) in higher magnification (left). **G-H.** CyCIF image of another representative p53.I (case C21-80, patient ID 35, BRCA2 Mut) with the presence of T_RM_ (white arrows), expressing the activation marker, PD1 (yellow arrows). **I-J.** CyCIF image showing a representative STIC.I, also shown in Figure 5 (STIC.I, case CD302.03(706), patient ID 38, BRCA1 Mut). Both CTL (magenta arrows) and T_RM_ (yellow arrows) are present, namely in the adjacent stroma. **K.** CyCIF image of the same ROI as J, showing the presence of these T cells with PD1 (orange arrows) or LAG3 (*punctate* expression) (white arrows for cytotoxic T, grey arrows for T_RM_) co-expression, indicating potentially CD8+ T cell exhaustion in STIC.I. Inset shows higher magnification of gray ROI. **L.** CyCIF image showing proliferative CD8+ T cell on the same STIC.I (orange arrow), ROI indicated in **I** and **J**. **M-N.** CyCIF image showing from another representative STIC.I, also shown in Figure 3 (case CD302.04(939), patient ID 40, BRCA WT). TP53 positive epithelium and ROI shown in (**M**) with the presence of T_RM_ (yellow arrows) in the epithelium of STIC.I (**N**). **O.** CyCIF image of same ROI as **N**, showing exhausted intra-epithelial T_RM_ in STIC.I, co-expressing both PD1+ LAG3+ (yellow arrows). **P-Q.** Heat maps showing normalized expression upregulation of selected genes related to T cell state, including naïve, dysfunction, and memory T cell. They include *HAVCR2* (encodes for TIM3), *CTLA4* observed in the STIC.I stroma or tumor stroma. Here, *PDCD1* (encodes for PD1), *ITGAE* (encodes for CD103), *TCF7* (encodes for TCF1), *CD274* (encodes for PDL1). Columns correspond to genes, and rows correspond to lesion stages. The heatmaps display the median of the posterior distribution derived from our Ordinal Bayesian model, relating log- and z-transformed gene expression to disease stage. Asterisks show significant changes in gene expression compared to the baseline stage (p53.C in (P) and FT.I in (Q)) Significance was determined using the proportion of the 95% highest density interval (HDI) within the Region of Practical Equivalence (ROPE, defined as 0.05 times the standard deviation). Comparisons with >95% of the HDI outside the ROPE were significant (*); >99% were very significant (**). Overall, exhausted CD8+ T cells, were observed even in incidental STIC.

Although the total number of CD8+ T cells decreased in later stages of HGSOC progression, the remaining CD8+ T cells were more likely to be either activated (PD1+ or Ki67+) or exhausted (LAG3+ or PD1+LAG3+), indicating a balance between active surveillance and T cell dysfunction (**Fig. 6C, 6D, 6G-6O**; **Supplementary Fig. S24, S25A-S25D**). While 11% of CD8+ T cells in p53.I showed signs of activation, this fraction rose significantly in STIC.I, STIC.C, and cancer (25-43%). Similarly, exhaustion markers increased in the epithelia compartments (**Fig. 6C**), with the proportion of exhausted CD8+ T cells rising 3- to 7-fold (∼2% were LAG3+ T_RM_ CD8 T cells in STIC.I, STIC.C, and cancer; 1% were LAG3+ CTL CD8+ T cells in STIC.C and cancer) (**Supplementary Fig. S24A**). In the stroma of STIC.I, STIC.C and cancer, ∼1-2% of CD8+ T cells were LAG3+ T_RM_ and ∼2% were LAG3+ CTL (**Fig.6D, Supplementary Fig. S24B**). The increased expression of RNA for *CTLA4* and *HAVCR2* (TIM3) (**Figure 6P-Q**) and GZMB protein and RNA (**Supplementary Figure S25E, S25F, S18G, S18H**) provided further evidence of activation and exhaustion in both the epithelium and stroma of HGSOC lesions (**Supplementary Fig. S26**). This data indicates that there is active immune surveillance by both T_RM_ and CTL populations, as well as conditions supporting immune editing and immune selection in HGSOC precursors.

## DISCUSSION

We have developed a comprehensive resource of multiplexed imaging and spatial transcriptomics data to study HGSOC development, from precancer lesions (incidental p53 signatures and incidental STICs) to invasive cancer. Our analysis reveals that localized IFN signaling and CIN are early events that increase with progression, accompanied by significant immune microenvironment reorganization. Activated T_RM_ and components of the NK-cDC1 axis are already present at the p53 signature stage, indicating early immune surveillance at the inception of HGSOC development in the fallopian tube. This immune response intensifies with increased activated CTL activity in incidental STICs. However, during the transition from incidental STIC to cancer-associated STIC, we observe a significant decrease in the NK-cDC1 axis. This decline coincides with an increase in activated CD8+ T cells, exclusion and dysfunction of these cells, reduced CD4+ T cell-APC interactions, and an increase in Tregs. These findings highlight a dynamic competition between immune surveillance and immune suppression, where initial immune responses in early lesions are progressively countered by immune-suppressive mechanisms as HGSOC progresses (**Fig. 7**).

**Figure 7:**
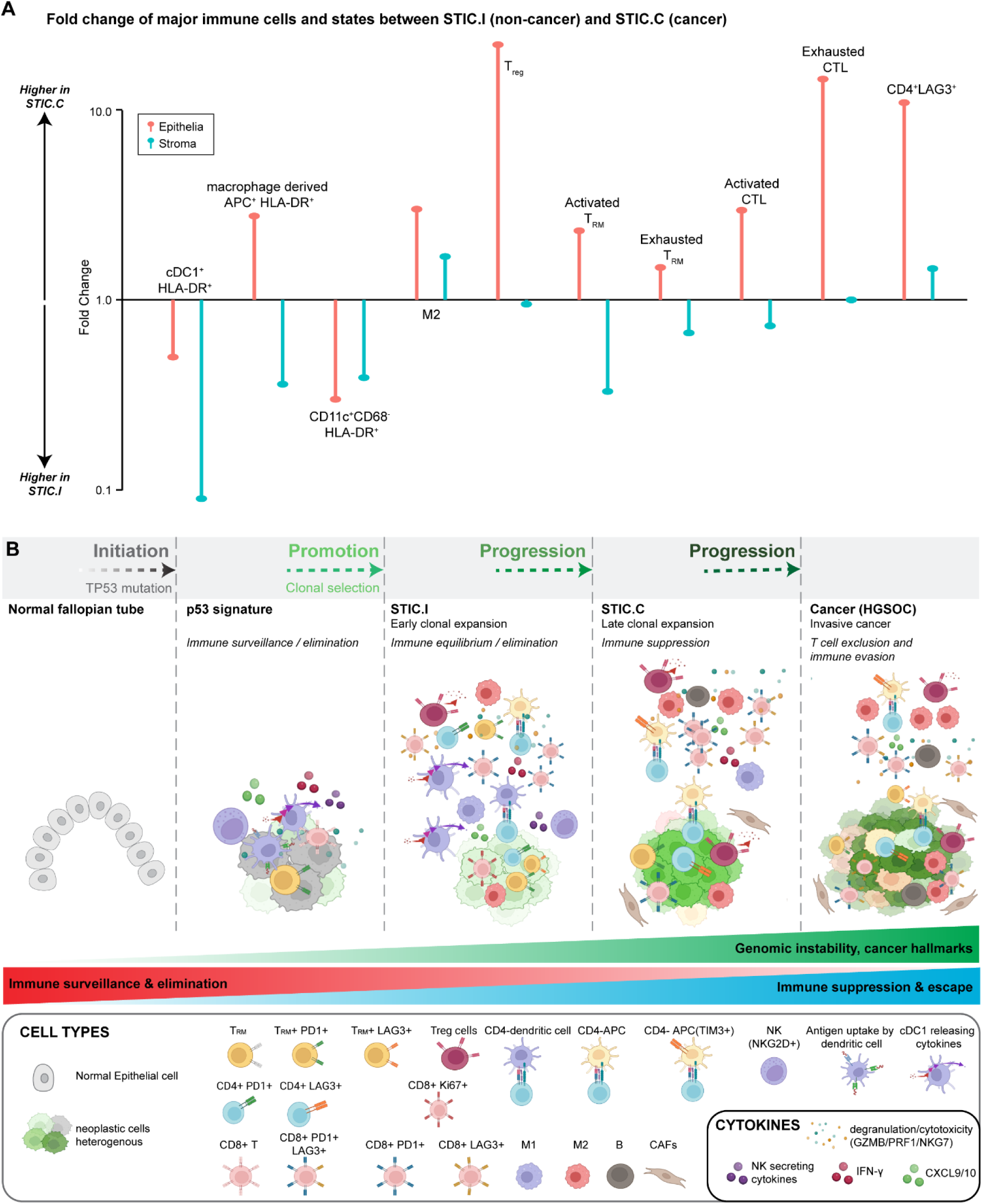
Evolution of the precancer ecosystem during HGSOC progression. **A.** Lollipop plot shows the relative difference (fold difference) of major immune cells and state between STIC.I and STIC.C. Epithelial and stroma regions are both shown. The fold difference is calculated from the average proportion of each cell state (proportion of STIC.C/STIC.I). Fold difference of 1 indicates no change, >1 indicates that cell type is higher in STIC.C and <1 indicates higher in STIC.I. **B.** This schematic illustrates the stages of HGSOC progression, highlighting the temporal evolution of cancer hallmarks and immune-precancer/cancer interactions. Cancer arises from genetic alterations, where mutations and aneuploidy, and other cancer hallmarks, under positive selection, lead to tumor development. However, cells with oncogenic mutations can remain latent for decades, and most never progress to malignancy. Evidence suggests that cancer originates from these “phenotypically normal” clones with driver mutations, followed by a clonal expansion. Along the progression axis, these clonally expanded cells accumulate additional mutations and encounter a gradual decline in anti-tumor responses, with an increase in immune-suppressive and dysfunctional immune cell subtypes. Our multimodal precancer atlas profiling shows how the TME evolves over time. Early in precursor development, cytotoxic immune responses emerge despite lower genomic instability compared to advanced tumors. Innate immune mechanisms (NK-cDC1-CTL axis) and tissue-resident memory T cells (T_RM_) play a key role in controlling p53 signature outgrowth. Aneuploidy or extrinsic factors enhance immune surveillance, removing precancer cells before disease outgrowth. During early STIC clonal expansion, we observe an initial immune response characterized by IFN activation, an increase in activated cDC1 and activated APCs, and the presence of NK cell secreted chemokines (which further attracts cDC1), indicating active immune surveillance. Interactions between APCs and activated CD4+ and CD8+ T cells provide further evidence of immune engagement. However, suppressive immune populations, such as M2-like macrophages and Tregs, emerge during this stage; STIC in this equilibrium phase simultaneously experience a cytotoxic and immunosuppressive environment with activation of tumor-promoting pathways. Late-stage clonal expansion has fewer CD8+ T cells with less interaction with APC and CD4+ and CD8+ T cells, more exhausted CD8+CTL and CD4+ (LAG3+), almost no NK and cDC1 cells, and increased suppressive APC. Overall, the transition from STIC to tumor is driven by hallmark mechanisms, such as TGF-β, which exclude CTLs, altered cytokines and fibroblast profiles, and induction of EMT and migration programs. The dotted arrows represent hypothetical timing, with the transition from p53 signature to early STIC taking longer than the progression from early to late STIC.

The molecular heterogeneity observed in precancer lesions offers insights into the interactions between the immune response and precursor phenotypes. Early localized IFN activation in p53 signatures and STICs, combined with the gradual accumulation of tumorigenic programs downstream of cGAS-STING or IFN pathways – such as NFκB, IRDS or IL-6 induced STAT3 pathways^40,72,89^ – may drive immune editing and selection in precursor populations. Traditionally this heterogeneity and extensive genomic instability (including MN rupture and catastrophic genomic events that drive genome evolution, immune evasion, and disease spread^40,61,62,69^) have been viewed as hallmarks of later tumor evolution and chemoresistance^3,14,25,54,90^. However, our data suggest that these processes are likely influential much earlier in HGSOC development. This phenotypic variability in STIC lesions, including differences in proliferation and DNA damage, may inform diagnostic risk stratification, by identifying STIC lesions with increased potential for malignant transformation and dissemination^91,92^. This variability coupled with known genetic alterations in HGSOC, such as 6p loss leading to MHC-class I loss and 6q loss affecting *IFNGR1* (Interferon Gamma Receptor 1)^25,54,90,93^, highlights the potential for intricate interplay between genetic mutations, precursor cell phenotypes, and localized immune interactions during cancer development^94^.

Immune evasion is a critical factor in HGSOC progression. Along the progression axis, IFN signaling coincides with HLA-A expression, responsible for antigen presentation to CTLs. However, upregulation of HLA-E expression may represent a key immune suppression mechanism during STIC clonal expansion. This aligns with findings in other cancers, where high HLA-E expression inhibits NK cell activity by engaging NKG2A receptors, thereby suppressing NK cell-mediated cytotoxicity^45,47,58–60^. The observed decline in the NK-cDC1 axis and dysfunction in adaptive immunity suggests that targeting NK cell reactivation using therapies like humanized anti-NKG2A antibodies (e.g., Monalizumab^58^) could be a promising strategy for early intervention, particularly in high-risk patients with incidental STICs linked to peritoneal cancer development^9^.

Immune surveillance, driven by activated NK cells, cDC1, and T_RM_ cells, along with elevated *IFNG* and *TNF*^75,87,88^ in the adjacent stroma, likely restrains the progression of early lesions, such as p53 signatures. This environment, sustained by feedback established between these immune cells and recruited CTLs, may limit abnormal growth and prevent cancer progression. However, for clonal selection and expansion to occur, additional factors are likely required^95–97^ over the course of decades^7,8,10,11,13^, such as repeated ovulatory-related stress leading to epithelial damage, increased cell proliferation, and genomic instability^8,98^. A shift in the IFN response, from early immune surveillance to chronic stimulation, may further promote IRDS and IFN-ɛ-suppression^40^, supporting DNA damage tolerance and immune suppression. Furthermore, macrophage-derived APCs, while increasing in prevalence in later stages, show reduced interaction with CD4+ T cells, indicating a functional shift that further promotes the transition from activation to immune suppression^79,81^, facilitating the progression to more advanced stages of disease.

Our findings align with emerging concepts of precancer hallmarks, including the failure of the NK-CTL axis, evolving immune suppression mechanisms, and microenvironment remodeling that promotes immune evasion^73,99^. These mechanisms are likely critical drivers of HGSOC development. In addition to offering insights into the temporal and spatial dynamics of the immune landscape and molecular transitions throughout HGSOC progression, this study also provides an accessible and large-scale public resource for further exploration of the interactions between transcriptional changes, cellular phenotypes, and spatial organization of precancer ecosystems. The integration of our spatial transcriptomics and imaging data into cBioPortal will enable future studies to uncover additional mechanisms driving cancer development and progression, ultimately advancing strategies for prevention, diagnosis, and treatment^100^.

### Limitation of the study

While this study provides insights into the immune landscape of HGSOC development and represents the largest available dataset of transcriptional and multiplexed imaging data of precancer, increasing the sample size will further enhance our understanding of the immune response across the HGSOC spectrum. In addition, 25% of STIC lesions associated with cancer have been reported to be disseminated cancer cells, reflecting the complexity of studying HGSOC progression in clinical settings^12^. Expanding the cohort to include more incidental STIC lesions would allow for the discovery of additional mechanisms relevant to disease progression and improve the identification of markers linked to clinical outcomes, offering a more complete understanding of HGSOC progression.

## METHODS

### Patient specimens and experimental design

In total, 43 patients were identified from the University of Pennsylvania’s (UPenn) and Swedish Cancer Institute, Seattle hospital databases, which met our criteria of the presence of either incidental STIC lesions (n=9), incidental p53 signatures (n=10), or STIC with concurrent carcinoma (n=24). After institutional review board approval, serial sections of 5uM thickness were processed. H&E stain was performed from the same block for CyCIF and GeoMx analysis (R.D., N.S., S.C) to confirm the diagnosis. One specimen per patient was processed, except one, patient 11 (bilateral STIC with concurrent carcinoma whereby both specimens were processed). In total, 44 FFPE specimens were collected, and all were processed for multi-plex imaging using CyCIF. For micro-region whole transcriptomics (GeoMx), 35/44 specimens were available and processed with more than 600 regions of interest (ROI) (n= initial 603 ROI collection), including normal FT/Fimbriae, precancer lesions and/or cancer. Tissue processing for both techniques are in **Supplementary Methods**.

### 2D Cyclic Immunofluorescence

Protocol for CyCIF was performed as described as Lin *et al* ^28,53^. The detailed protocol is available in protocols.io (https://doi.org/10.17504/protocols.io.bjiukkew). In brief, the slides were baked at 55-60^ο^C for 55 min prior to shipping. Then upon receiving the slides, the BOND RX Automated IHC/ISH Stainer was used to bake FFPE slides at 60 ^ο^C for 15 minutes, to dewax the sections using the Bond Dewax solution at 72 ^ο^C, and for antigen retrieval using Epitope Retrieval 1 (Leica^TM^) solution at 100 ^ο^C for 20 minutes. Slides underwent multiple cycles of antibody incubation, imaging, and fluorophore inactivation. All antibodies were incubated overnight at 4 ^ο^C in dark. Coverslips were wet mounted using 200 µL of 50% Glycerol/PBS before imaging. Images were acquired using a 20x objective (0.75 NA) on a CyteFinder slide scanning fluorescence microscope (RareCyte Inc. Seattle WA). Fluorophores were inactivated using a 4.5% H_2_O_2_, 24mM NaOH/PBS solution and an LED light source for 1 hour. The detail of the antibody panel used in this study were detailed in Supplementary File S4.

### Image processing and quality control

Image processing and analysis was performed with the Docker-based NextFlow pipeline MCMICRO^33^ and with customized scripts in MATLAB^53^ and R (v 4.3.3) as described previously. Briefly, raw images were stitched and registered from the different tiles and cycles after the acquisition using the ASHLAR^101^ module within the MCMICRO pipeline. After the registration step, the OME.TIFF files from each slide were passed through the quantification module of MCMICRO. Overall, UNMICST2^102^ was used for segmentation and quantification to generate single cell data. Details can be found at www.cycif.org. Quality Control (QC) of the single-cell data includes removing cycles where tissue loss was observed, as previously published to have the final single-cell feature table^53^.

### Cell type identification

All samples and markers were gated independently using an open-source “gator” viewing and analysis tool as well as binary gating as described previously^53^. The details of gator can be found at https://github.com/labsyspharm/minerva_analysis/wiki/Gating. After generating the initial gate, visual inspection and adjustment was made to the final gating table to incorporate with single cell feature table. For cell type and state identification, pre-existing knowledge based on literature was used as described^53^.

### Cell population proportions

After calculating the average proportion of cells expressing single/double/tripe markers for cell phenotyping on MATLAB, downstream quantification across HGSOC stages, including statistical analysis, was performed in R (v 4.3.3). To test whether proportions of cells with a given phenotype among the whole populations of cells differed between disease stages, we used binomial Generalized Linear Mixed Models (GLMMs) implemented in the lme4 R package (v 1.1-34)^103^. For each ROI, the number of cells with the given phenotype (“successes”) and of all other phenotypes (“failures”) were modelled using the binomial distribution with a logit link function using the lme4 model formula cbind(n_success, n_failure) ∼ stage + (1 + stage | patient_id). ROIs were not independent since they were derived from the same patients more than once. To account for this, covariance and patient specific effects were modeled by including random intercepts and stage coefficients for each patient. Post-hoc contrasts between stages were performed using the emmeans R package with Benjamini-Hochberg (BH) correction for multiple testing. Due to the overdispersion of some immune populations, we used binomial GLMMs implemented in the glmmTMB R package (v 1.1.9)^104^ for six major subtypes of immune populations. For each ROI, the number of cells with the given phenotype (“successes”) and of all other phenotypes (“failures”) were modelled using the binomial distribution with a logit link function using the glmmTMB model formula cbind(n_success, n_failure) ∼ stage + (1 | patient_id) + (1 | observation_id). ROIs were not independent since they were derived from the same patients more than once. To account for this, covariance and patient specific effects were modeled by including random intercepts and stage coefficients for each patient. Overdispersion was controlled for by including an observation-level random effect (OLRE)^105^. This model specification minimized the Akaike information criterion (AIC) compared to other alternative specifications, including negative binomial, beta-binomial, and binomial models without OLRE. The data summary and statistics for all CyCIF analysis are in **Supplementary File S5 and S6**.

### Neighborhood analysis (LDA) and cell to cell interactions

LDA analysis for spatial topic analysis was performed using MATLAB *fitlda* function as described by.^53^. The pooled frequencies of all samples were used to train the final LDA model and 21 topics were isolated. To inspect cell to cell interactions, especially between two types of immune cells, a cell type dependent interaction score was generated^106^ by a custom MATLAB script, whereby >1 indicates a close proximity between two cell types and vice versa. The score was normalized against the distance of random sampling of two cell types of interest and then compared to different stages of HGSOC progression. To allow for the permutation step, only samples with both cell types >5 cells were included with further manual inspection of each cell type/sample for each stage of HGSOC progression. The details of cell types and size comparing STIC.I vs STIC.C is in **Supplementary File S5**.

### 3D Cyclic Immunofluorescence and image processing

One of the cases of STIC co-existing with cancer was processed (20 µm thickness) for super high-resolution imaging as detailed here^29^ using Zeiss LSM980 confocal microscope (**Supplementary Method**). The staining protocol is similar to standard 2D CyCIF with overnight antibodies incubation. The antibody is detailed in **Supplementary File S4**.

### Annotation, selection of regions of interest, and protocol for micro-region spatial transcriptomics

For the micro-region spatial transcriptomic profiling, we have used the GeoMx platform with the whole transcriptome (WTA) probe sets (NanoString, Seattle, USA) as previously published^73^. The ROIs were annotated by a board-certified pathologist (S.C) based on H&E and visualizing images from CyCIF (Cycle 1-6). Since we wanted to integrate RNA and protein expression, we have annotated all ROIs based on the presence or absence of *HLA-E* expression in both epithelium and epithelium-stroma boundary (i.e. adjacent stroma). Unlike the cancer group, adjacent stromal compartments of normal FT or Fimbriae were collected for all incidental cases when available (**Supplementary Fig S2**). NanoString GeoMx gene expression analysis utilizing the whole transcriptome (WTA) probe set was performed using previously described methods^73^. WTA probe set is in **Supplementary File S7**, keeping in mind that few probes were generic, meaning NanoString could not design those probes for specific receptors, such as *KLRC2* and *KLRC1*, such that both receptors are displayed by “*KLRC2*”. Briefly, a 5μm section was dewaxed and stained overnight with antibodies targeting epithelial (pan-cytokeratin) and immune cells (CD45), defining cell morphology and highlighting regions of interest. The section was hybridized with the WTA probes before being loaded into the instrument. In total, 603 ROIs were initially selected for collection and library preparation. Followed by QC, 542 ROIs representing different stages of the disease progression were used for downstream analysis (**Supplementary Methods**).

### Data processing and quality control for GeoMx data

All sample processing and sequencing were performed by the Dana Farber Sequencing or HMS facility. The quality control (QC) and the Quartile-3 (Q3) normalization of the initial data set were performed as suggested by NanoString using GeoMx DSP software, NanoString (v 3.1.0.221). The detailed steps of QCs are mentioned in **Supplementary Methods**. The ROI and annotation of these regions after the QC are detailed in **Supplementary File S2**. The ROI numbers for each lesion type are depicted in **Supplementary Figure S2**. The Count matrix file after the QC is in **Supplementary File S8**.

### Spatial Integration of CyCIF and GeoMx data

To establish the mapping between GeoMx ROIs and CyCIF data, the DNA channel in both whole-slide images was used. The process begins by deriving a global affine transformation between the downsized whole-slide images of the GeoMx and CyCIF datasets. In the second step, we query 2D image patches at full resolution, centered around the ROI centroids in the GeoMx image and their corresponding locations in the CyCIF image, based on the global affine transformation. These pairs of image patches are then used for a second round of affine registration. Both rounds of affine registration are conducted using ORB (Oriented FAST and Rotated BRIEF)^107^ feature detection and matching techniques.

### Differential gene expression and Gene Set Enrichment Analysis (GSEA)

To identify sets of genes that were highly or lowly expressed, differential gene expression was performed using GeoMx DSP software, NanoString (v 3.1.0.221) using the Q3 normalized counts. DSP software was used for Linear Mixed Models (LMM) with Benjamini-Hochberg (BH) correction to perform differential gene expression. Model formula: Lesions Type + (1| Scan_ID) whereby Scan_ID refers to the patient/slide ID. LMM is designed to handle data with repeated measurements from the same sampling unit and here Scan_ID was chosen as a random effect. DSP software’s custom R script provided by NanoString was used to visualize the data for differential gene expression as a volcano plot.

The output of differentially expressed genes was exported from the DSP software (i.e., a ranked list of differentially expressed genes between two sets of experimental conditions, such as p53.I (epithelial) *vs* STIC.I (epithelial), shown in **Fig. 2A, 2B**). Then gene set enrichment (GSEA) analysis using MsigDB^36^ Cancer Hallmark pathways and reactome pathways^37^ were performed. MsigDB GSEA was performed in R (version 4.3.3) using msigdbr (hallmark gene set, category == H) and fgsea packages. GSEA on reactome database was performed using GeoMx DSP software and visualized using R. The visualization of the GSEA was adapted^108^.

### To model the progression of HGSOC using published gene set by Bayesian Regression Modelling

To study the relationship between gene expression and cancer lesion progression we employed a Bayesian ordinal regression model implemented by the brms R package^109^. We preprocessed GeoMx expression counts by Q3 normalized to account for sequencing depth, followed by log10 transformation to stabilize variances. To account for differences in expression levels across genes, we further normalized the log-transformed values by scaling to a mean of zero and variance of one (z-transform). We fitted one model per gene using the model specification gene_expression ∼ mo(stage) + (1 + mo(stage) | patient_id) (**Fig. 2E, 2F, Supplementary Fig. S13**). The lesion stage was used as ordinal predictor mo(stage) with a monotonic constraint to enforce the assumption of an orderly sequence of stages. We accounted for repeat measurements from the same patients by including patient-specific random intercepts and stage coefficients. To model expression of gene sets, we modified the model to include another random effect and its interaction with patient_id (gene_expression ∼ mo(stage) + (1 + mo(stage) | patient_id * gene). This modification allows every patient and gene to have a different expression baseline and expression changes at each disease stage. The ‘*’ operator between patient and gene enables genes to behave differently in every patient (i.e., interaction). As a result, we can look at the expression along with the progression axis as a gene set, such as the IRDS gene set shown in **Supplementary Fig. S13E**. To investigate the effect of HLA-E expression on other genes, we further modified the model to include a fixed effect coefficient serving as binary indicator for presence or absence of HLA-E in the CyCIF image of the ROI (gene_expression ∼ mo(stage) * hlae_e + (1 + mo(stage) * hlae_e | patient_id * gene). We tested this modified model first to confirm whether HLA-E expression in protein and RNA levels match. **Supplementary Fig. S13C** indicates that HLA-E RNA expression is higher in clones that are HLA-E+ by CyCIF, including an increased trend observed from FT.C to cancer. For significance testing, we used the proportion of the 95% highest density interval (HDI) within the Region of Practical Equivalence (ROPE, 0.05 times the standard deviation). Comparisons with >95% of the HDI outside the ROPE were significant (*), while those with >99% were considered very significant (**)^110^. In most cases, matched FT was chosen as a reference, meaning p53.I and STIC.I were compared to FT.I, and STIC.C and cancer were compared to FT.C. One exception was the stromal component from the cancer group, where we used p53.C as a reference due to the unavailability of normal FT or Fimbriae from this group.

### Statistical analysis

All statistical significance is considered p<0.05 unless stated otherwise. Other statistical analyses, such as t-test were performed using R (version 4.3.3) or Graph Pad Prism (v 10.0.2 (232)).

### Schematic diagrams

Schematics in Figure 1 and Figure 7 were made with BioRender.

## Supporting information

cohort description

GeoMx ROI annotations

gene sets

antibody panels for CyCIF

data summary

Stat summary

WTA probe details

Q3 normalised count matrix file

Supp Video of micronuclei rupture

## Data and software availability

Both CyCIF images and GeoMx data will be available through https://cbioportal.org/study/summary?id=ovary_geomx_gray_foundation_2024.GeoMx data is available as Count Matrix as a **Supplementary File S8** with this manuscript. New Codes for this manuscript will be available prior to publication at https://github.com/labsyspharm/stic-ms-2024.

## ACKNOWLEDGMENTS

We thank the MicRoN core facility at HMS for providing access to an LSM980 Airyscan 2 microscope.

## AUTHOR CONTRIBUTIONS

TK, JL, SCoy, RJP, MLL, JSL, YX, and CY performed experiments, spatial transcriptomics and imaging. TK, JL, CH, and YC performed data analysis.

JL, YC, and JM developed data visualization and management approaches.

SK, SMG, NS, GM, EJ, DKO, JRH, CD, and RD provided and annotated reagents.

TK, JL, JBT, PKS, RD, and SS wrote the paper and all authors reviewed drafts and the final manuscript.

TK, JL and CH prepared the figures.

NS, SChan, EJ, CD, RD, and SS supervised clinical biospecimen collection and management.

IdB, BAS, RK and NSchultz supported data visualization using cBioPortal.

PKS, RD, and SS supervised the overall research.

