## Supplementary material for "Multimodal Spatial Profiling Reveals Immune Suppression and Microenvironment Remodeling in Fallopian Tube Precursors to High-Grade Serous Ovarian Carcinoma": Supp Figures

**Supp Figure S1 (Related to Figure 1)**

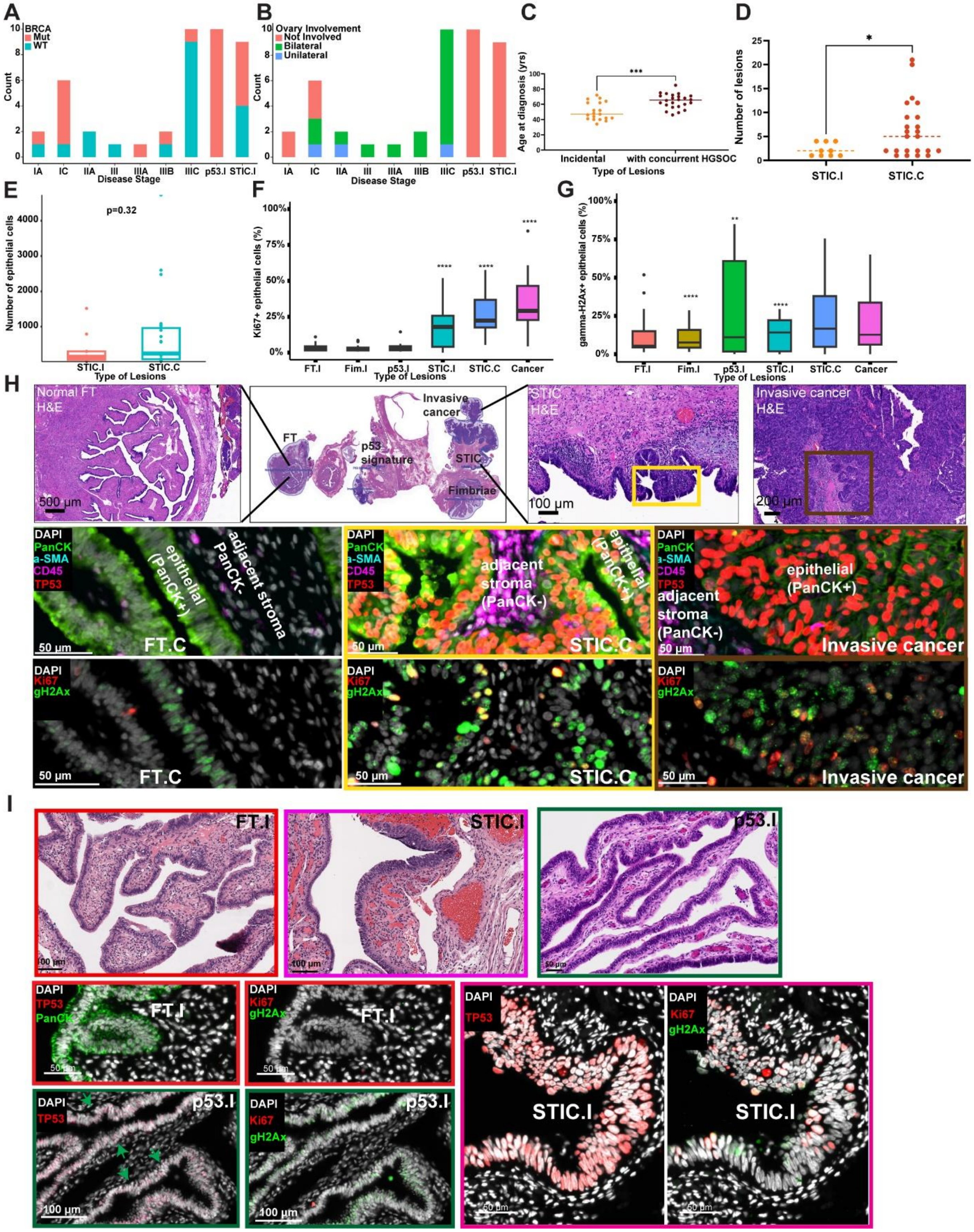

**Supp Figure S1. CyCIF images suggested higher proliferation with HGSOC progression:**
A. Breakdown of BRCA status across disease stages including carcinoma (Stage IA- Stage IIIC).

B. Description of ovary involvement across disease stages including carcinoma (Stage IA- Stage IIIC).

C. comparison of age of the patients between incidental (n=19) and cancer(n=24) groups. Solid line represents median, ***p<0.001, Mann-Whitney t-test.

D. The comparison of the number of STIC lesions between STIC.I (n=9) and STIC.C (n=23). The dotted line represents median, *p<0.05, Mann-Whitney t-test.

E. Box plot comparing the number of epithelial cells between STIC.I (n=9) and STIC.C (n=23). The median number of cells in STIC.I are 148 *vs* 242 in STIC.C (p=0.32, unpaired t test with BH correction (using R v 4.3.3).

F-G. Box plots comparing (F) Ki67+ and (G) ɣ-H2Ax+ epithelial cells across disease stages, from tissue imaging, expressed as a percentage. Number of specimens per group as follows: FT.I (n=13), Fim.I (n=15), p53.I (n=10), STIC.I (n=9), STIC.C (n=23), and Cancer (n=20). The solid line indicates the median within the interquartile range, with whiskers extending to a maximum of 1.5 times the interquartile range beyond the box (E-G). Outliers were shown as dots. Black asterisks indicate significant differences in stages compared to the FT.I; **p<0.01, ****p<0.0001. Binomial Generalized Linear Mixed Models (GLMMs) taking patient ID as random effect, implemented in the lme4 R package (v 4.3.3).

H. An example of a case of STIC with concurrent HGSOC (Case RD-23-002, patient ID 9, BRCA2 mutant, Stage IC HGSOC), also shown in Figure 1F. The H&E specimen (top) shows different histology on the specimen, such as FT, STIC, and invasive cancer. The CyCIF image shows increased p53 mutant epithelial cells (middle panel), Ki67+, and ɣ-H2Ax+ epithelial cells (bottom panel) with disease progression. Insert from Panel H for STIC.C, outlined with a yellow box and invasive cancer, outlined with a brown box. CyCIF image also depicts the epithelial compartment (PanCK+) and the adjacent stromal compartment (PanCK-) to the epithelia for all histology.

I. Top: H&Es of an example of a STIC.I case (outlined with a magenta box) with matched FT.I (outlined with a red box) (case CD302.04(939), patient ID 40, BRCA WT, STIC.I) and an example of a p53.I case (outlined in a green box) (case C21-22 patient ID 28, BRCA1 Mut, p53.I). Bottom: CyCIF images showing p53 mutant, Ki67+ and ɣ-H2Ax+ epithelial cells. Green arrows of p53.I indicate the “p53 signatures” regions of the case.

**Supp Figure S2 (Related to Figure 1)**

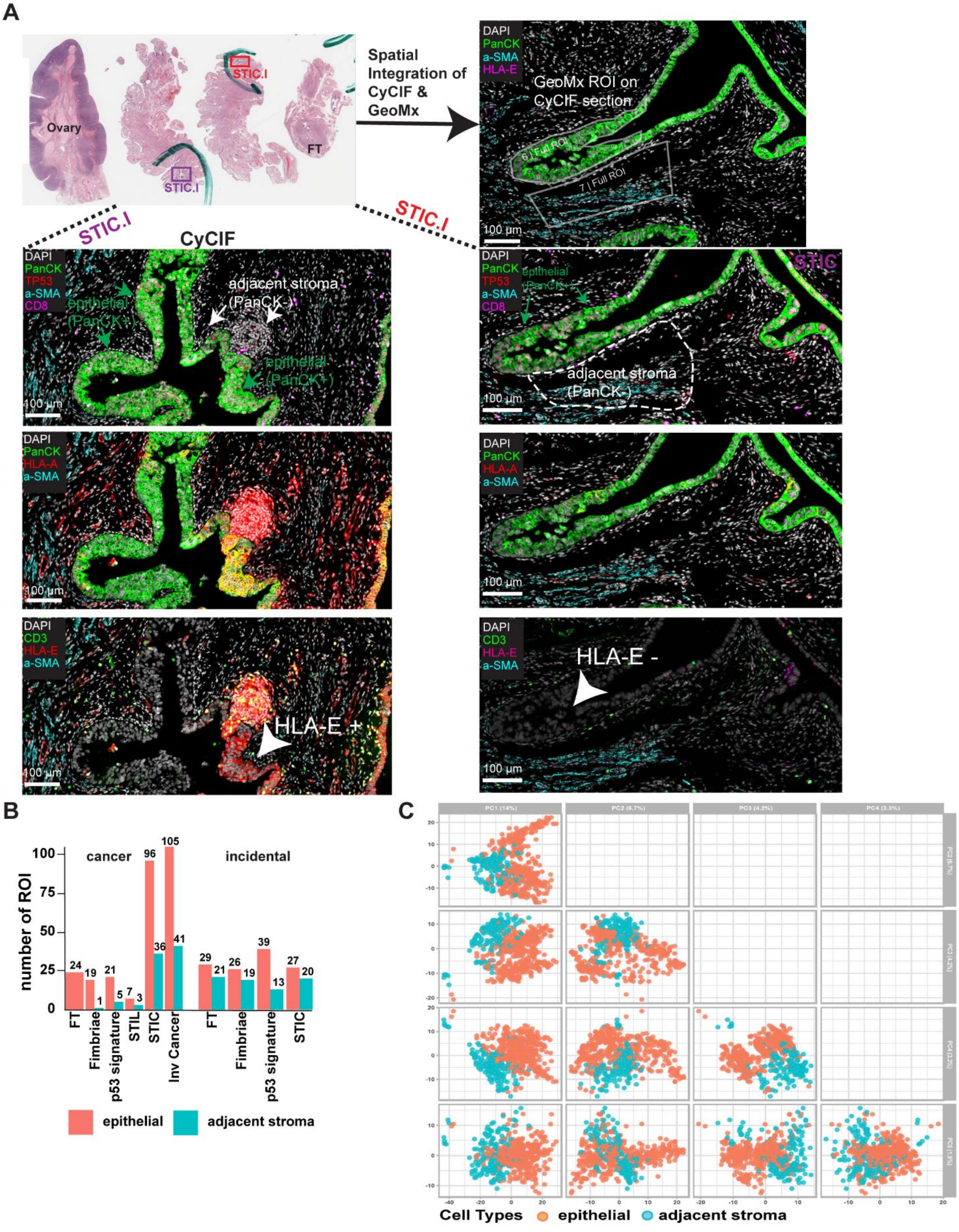

**Supp Figure S2. Spatial integration of tissue imaging and spatial whole transcriptome (GeoMx) and ROI selection:**

A. Experimental design for histology-guided (H&E) GeoMx ROI collection and spatial integration of GeoMx ROI on CyCIF specimen. The H&E on top showing a STIC.I case having 2 regions of STIC lesions, outlined with a red and purple box (case CD302.8(7923), patient ID 43, BRCA WT STIC.I). The epithelial compartment was shown with green arrows, and the adjacent stroma to the epithelia was shown either with a white arrow (left) or a white dashed box (right) for both STIC lesions, purple and red, respectively. For example, ROI 6 indicates the epithelial (PanCK+) (green arrow) and ROI 7 indicates adjacent stroma (PanCK-) (outlined with white dashed box) of STIC (outlined with a red box on H&E). CyCIF image from this specimen also shows the heterogeneity of STIC.I lesion from the same patient in terms of the expression of MHC-class I. STIC.I, outlined with a red box on H&E has no overexpression of MHC-class I (both HLA-A and HLA-E), shown by CyCIF image on the right. On the left, another STIC.I lesion (outlined with a purple box on H&E) has overexpression of MHC-Class I on some epithelial and adjacent stroma compartments.

B. Number of Regions of Interest (ROIs) from spatial whole transcriptome (GeoMx) passed after QC from both incidental and cancer group. ROIs were collected from epithelial (PanCK+) and adjacent stroma compartments (PanCK-). In total, 567 ROIs are in **Supplementary File S2**. Of these, floating cancer (n=15) and STIL associated with cancer (n=10) were not used for downstream analysis. ROIs from STIL associated with cancer were not included in the final analysis due to fewer ROIs and the unavailability of incidental STIL cases. Floating cancers were not considered as part of invasive cancers and were excluded from downstream analysis for this study.

C. Principal component analysis (PCA1 to PCA5) plots all spatially resolved GeoMx transcriptomic ROIs based on cell types (epithelial or adjacent stroma).

**Supp Figure S3 (Related to Figure 1)**

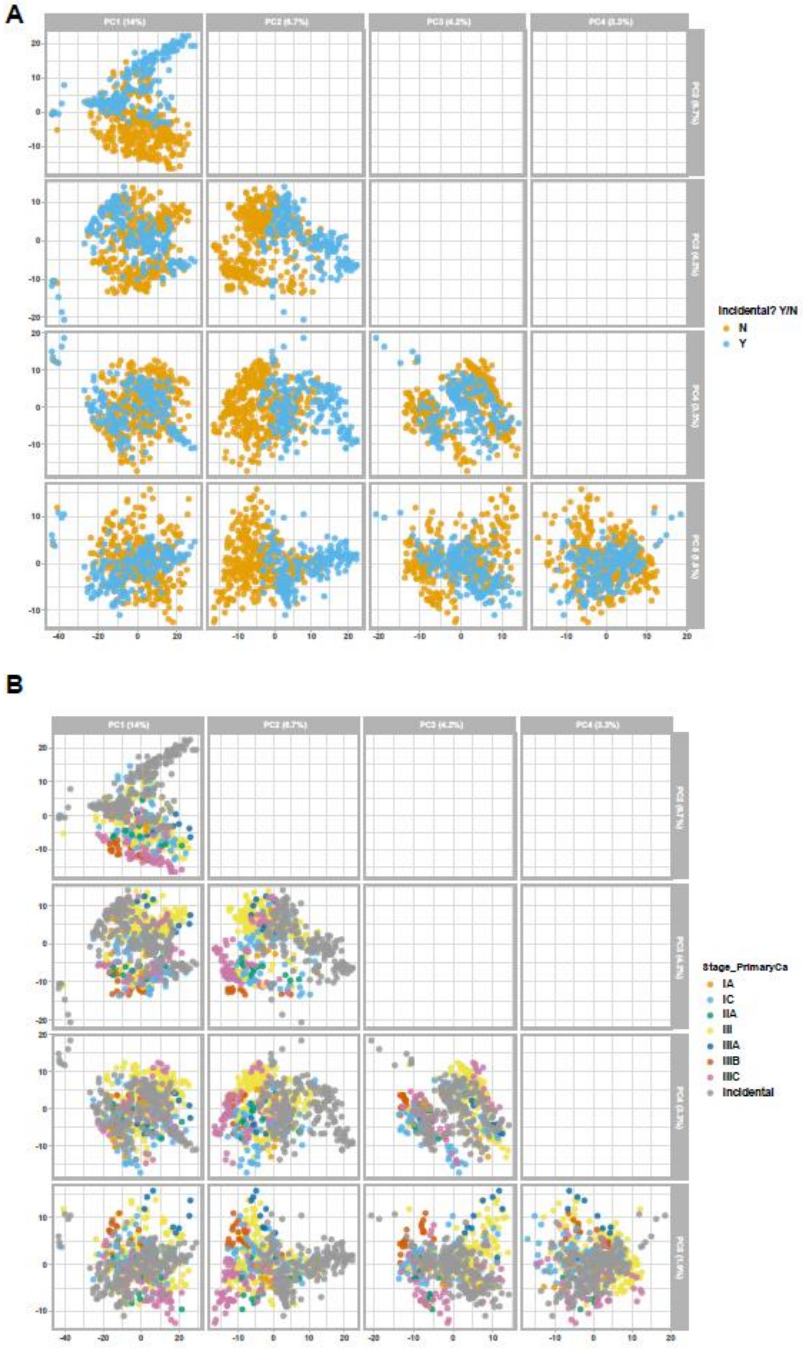


**Supp Figure S3. Additional PCA plots from GeomX data:** A-B. Incidental and cancer cases were processed for ROI collection, followed by sequencing on multiple batches. PC2 appears to separate the difference between incidental and cancer group (A) or incidental and end-stage disease (IIIC) (B).

### Supp Figure S4 (Related to Figure 2)

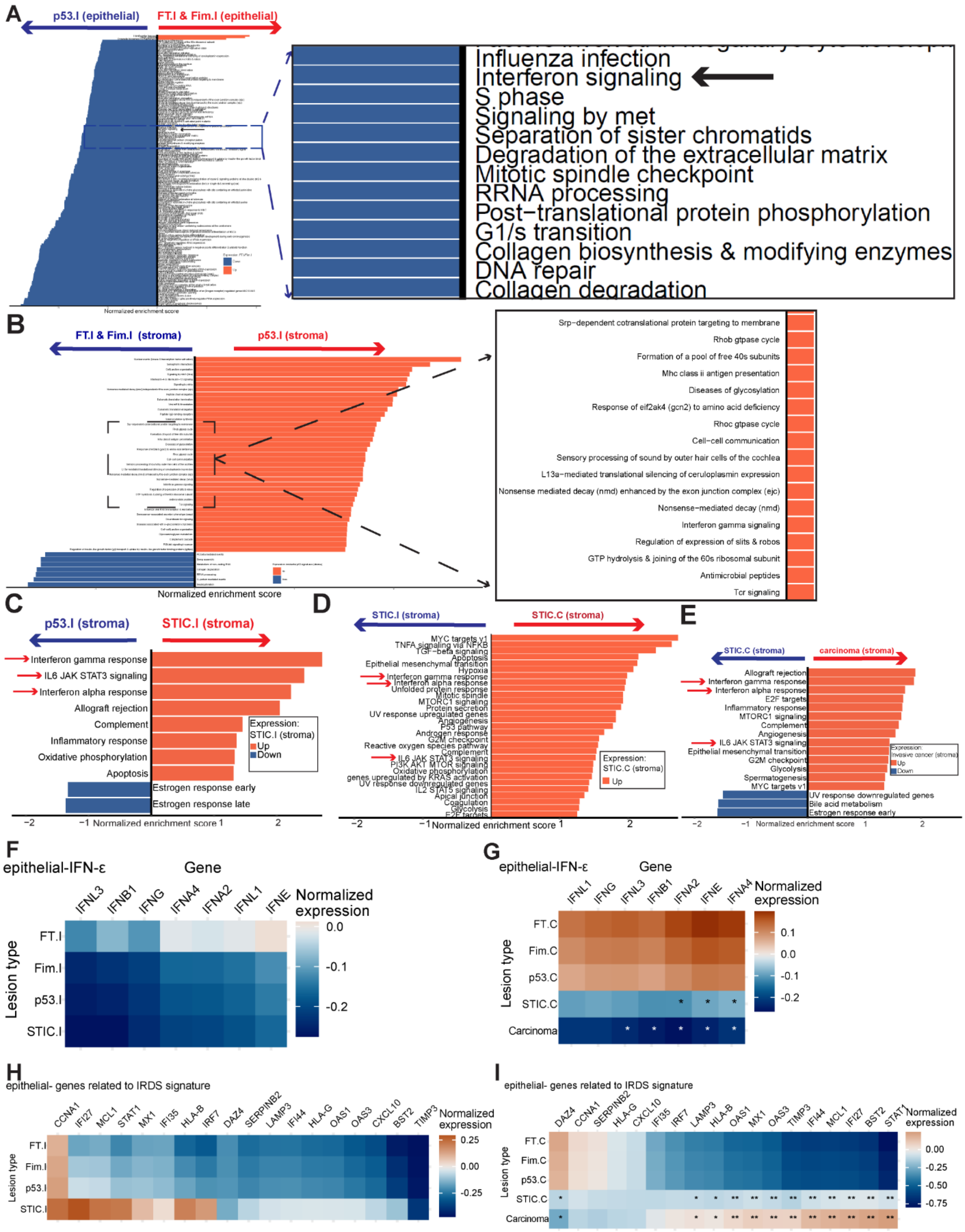

**Supp Fig S4. Temporal evolution of HGSOC from its pre-cancer lesions shown by spatial transcriptomics:**

A-B. Gene Set Enrichment Analysis (GSEA): Reactome pathway enrichment showing pathways associated with disease progression between epithelial of FT.I and Fim.I and p53.I. FT.I and Fim.I were combined as one group and compared to p53.I for differential gene expression. Progression of p53.I was predominantly associated with interferon (IFN) and proliferative gene sets both in epithelium (A) and the adjacent stroma (B). Some of these pathways are highlighted in insert. IFN pathway was indicated with a black arrow in the epithelium.

C-E. MsigDB Cancer Hallmark gene sets associated with disease progression in the adjacent stromal compartments between (C) p53.I (n=13) and STIC.I (n=20), (D) STIC.I (n=20) and STIC.C (n=36), (E) STIC.C (n=36) and invasive carcinoma (n=41). Progression of STIC.I (C) was predominantly associated with IFN; STIC.C (D) was predominantly associated with EMT, TGF-β and hypoxia gene sets and all of these pathways seem to be further upregulated in invasive carcinoma (E). IFN related pathways were shown with red arrows. A. Ranking of the pathways was based on NES>2 as well as on adjusted p-value <0.001. B. Ranking of the pathways was based on adjusted p-value <0.9. C-E. Ranking of the pathways was based on adjusted p-value <0.05. B-E. “stroma” indicates adjacent stroma to the epithelium as shown in **Supplementary Fig. S2**. F-I. In order to look into more details of IFN pathway and how it changes from normal FT to STIC.I or STIC.C to carcinoma, in addition to the IFN-α and IFN-ɣ gene sets shown in **Figure 2**, Bayesian regression modeling was applied to genes related to IFN-epsilon(ε) and IRDS gene sets.

F-G. Heatmap showing normalized expression of genes in the epithelia related to IFN-epsilon(ε), indicating, only in STIC.C and cancer group showed a downregulation of genes related to IFN- ε. IFNE is the transcript for IFN- ε (part of Type-I IFN), and other transcripts shown here are cytokines that are part of Type I and Type II IFNs, as previously published.

H-I. Heatmap showing normalized expression of genes in the epithelia related to IRDS, an indication of chronic IFN activation, namely observed in STIC.C and tumor. Median of the posterior distribution was shown in heatmaps. F-I. Columns correspond to individual genes, and rows correspond to types of lesions. Bayesian modeling was applied to see the relative gene expression changes in the incidental or cancer group compared to the matched FT. Median of the posterior distribution was shown in heatmaps. Significance testing used the proportion of the 95% highest density interval (HDI) within the Region of Practical Equivalence (ROPE, 0.05 times the standard deviation). Comparisons with >95% of the HDI outside the ROPE were significant (*); >99% were very significant (**)

**Supp Figure S5 (Related to Figure 2)**

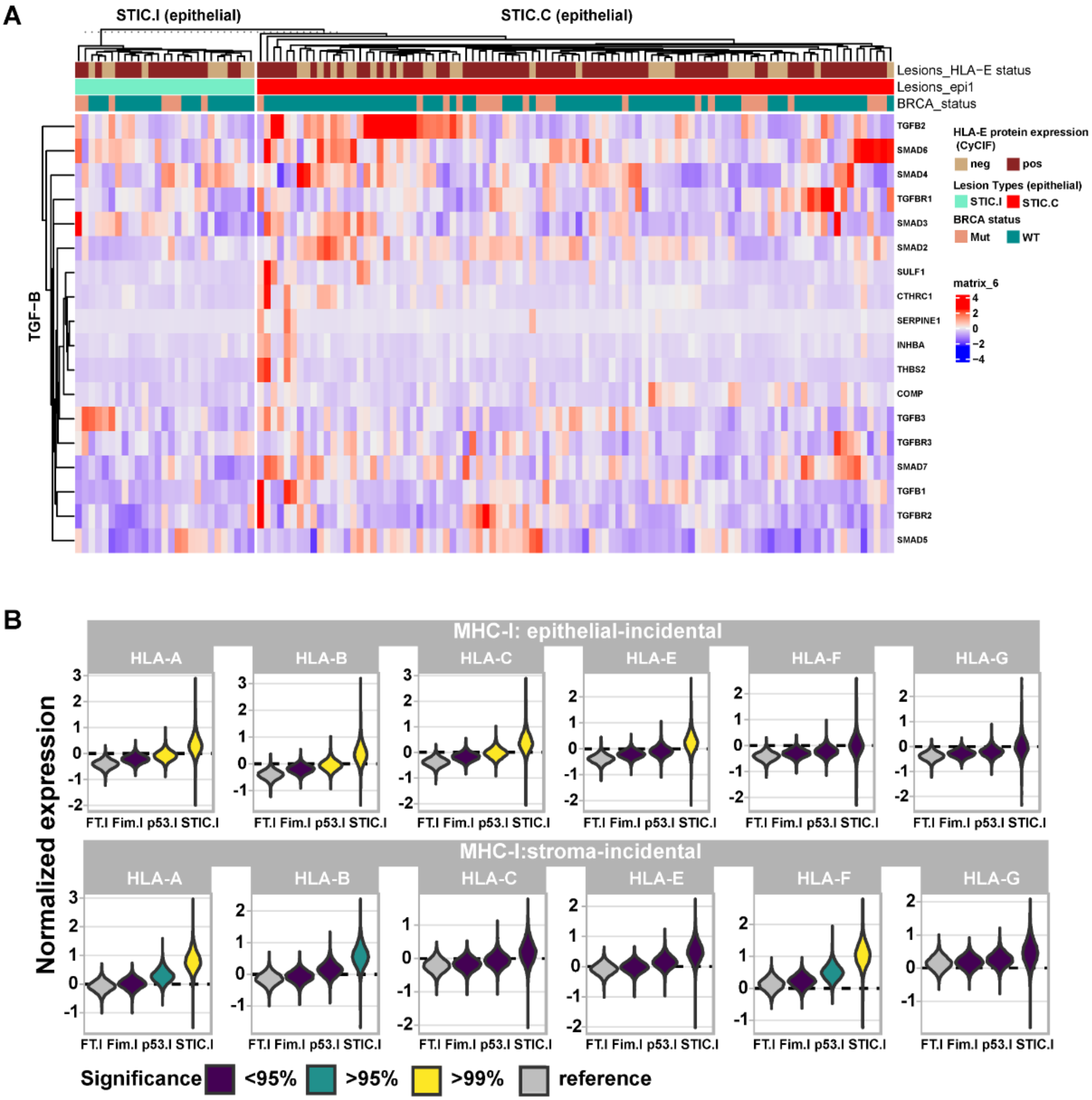

**Supp Figure S5.** A. Heatmaps showing expression of the genes related to TGF-β pathway activation and comparing between STIC.I and STIC.C epithelium with a supervised clustering. GeoMx expression counts were Q3 normalized to account for sequencing depth and log10 transformation to stabilize variances. To account for differences in expression levels across different genes, the log-transformed values were further normalized by scaling to a mean of zero and variance of one (z-transform). Columns correspond to individual ROI collected for GeoMX.

B. Violin plots showing normalized expression of genes in the epithelia related to both classical (HLA-A/B/C) and non-classical (HLA-E/F/G) MHC-class I in incidental cases both in the epithelia and the adjacent stroma. Bayesian modeling was applied to see the relative gene expression changes in the incidental or cancer group compared to the matched FT. Median of the posterior distribution was shown in heatmaps. Significance testing used the proportion of the 95% highest density interval (HDI) within the Region of Practical Equivalence (ROPE, 0.05 times the standard deviation). Comparisons with >95% of the HDI outside the ROPE were significant (*); >99% were very significant (**).

**Supp Figure S6 (Related to Figure 3)**

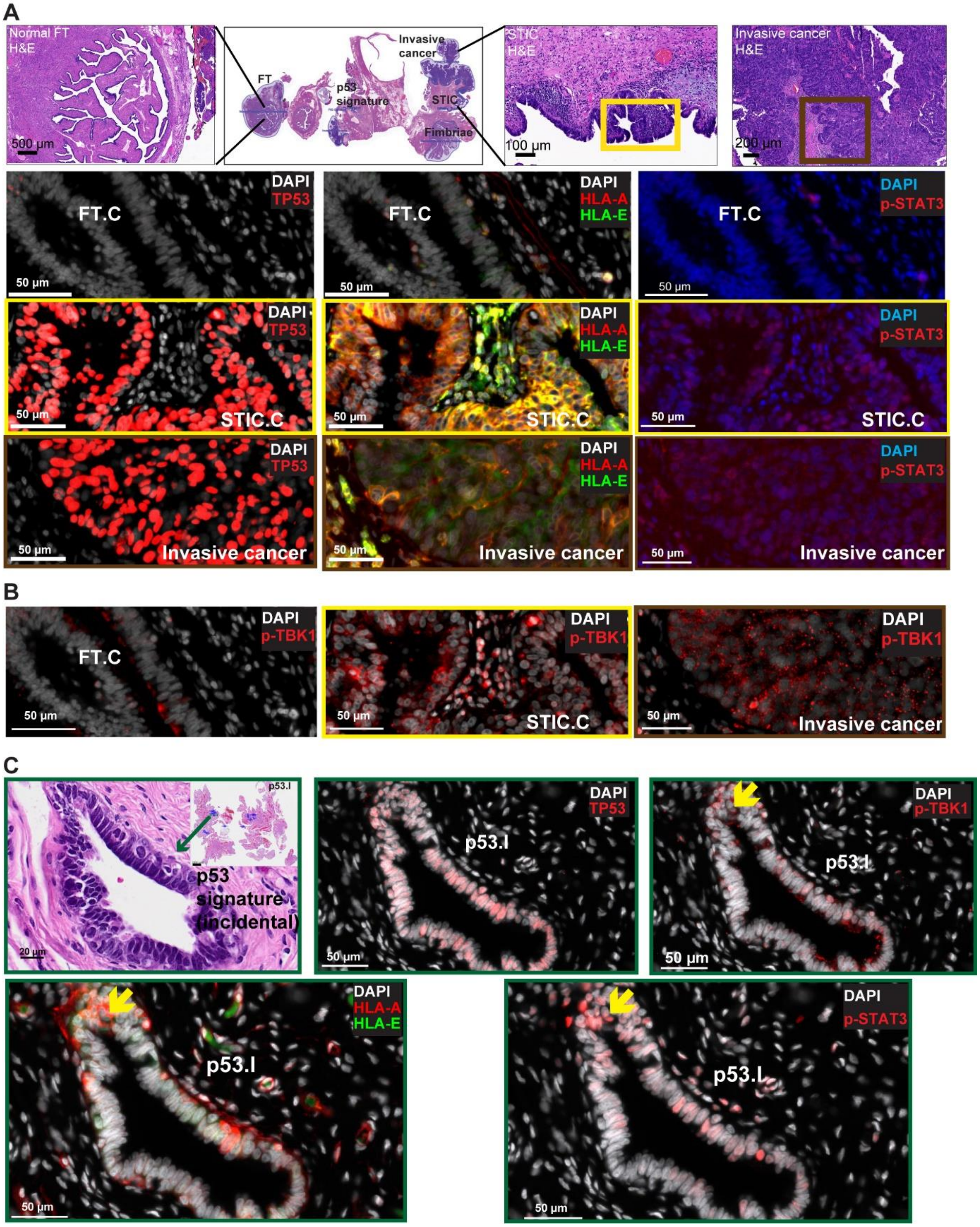

**Supp Fig S6. IFN pathway activation was not only restricted to the very early stage of HGSOC progression, suggesting a positive selection:**

Since IFN signaling pathway was upregulated in p53.I and STIC.I stage, we validated few key upstream and downstream of IFN signaling pathway in STIC.C and cancer specimens by CyCIF (protein level). A. CyCIF images representing downstream of IFN signaling pathway activation, such as overexpression of MHC-Class I (both HLA-A and HLA-E) and p-STAT3 in STIC.C and invasive carcinoma component compared to FT.C. A representative case of cancer group is shown here, also shown in **Figure 1**, STIC with concurrent HGSOC (Case RD-23-002, patient ID 9, BRCA2 mutant, Stage IC HGSOC). The H&E specimen (top) shows different histology on the specimen, such as FT.C, STIC.C, and invasive cancer. The CyCIF image (bottom) shows increased p53 mutant, both HLA-A+ and HLA-E+ and p-STAT3+ epithelial cells with disease progression in cancer group.

B. CyCIF images showing an increased p-TBK1+ epithelial cells (cytosolic/*punctate* expression) in STIC.C and tumor compared to matched FT.C.

A-B. STIC.C was outlined with a yellow box on H&E and tumor was outlined with a brown box on H&E.

C. H&E-another representative case of p53.I is shown suggestive of activation of upstream and downstream of IFN pathway activation, outlined with a green box. Yellow arrows were used to show examples of cells expressing all IFN-activation markers.

Supp Figure S7 (Related to Figure 3)

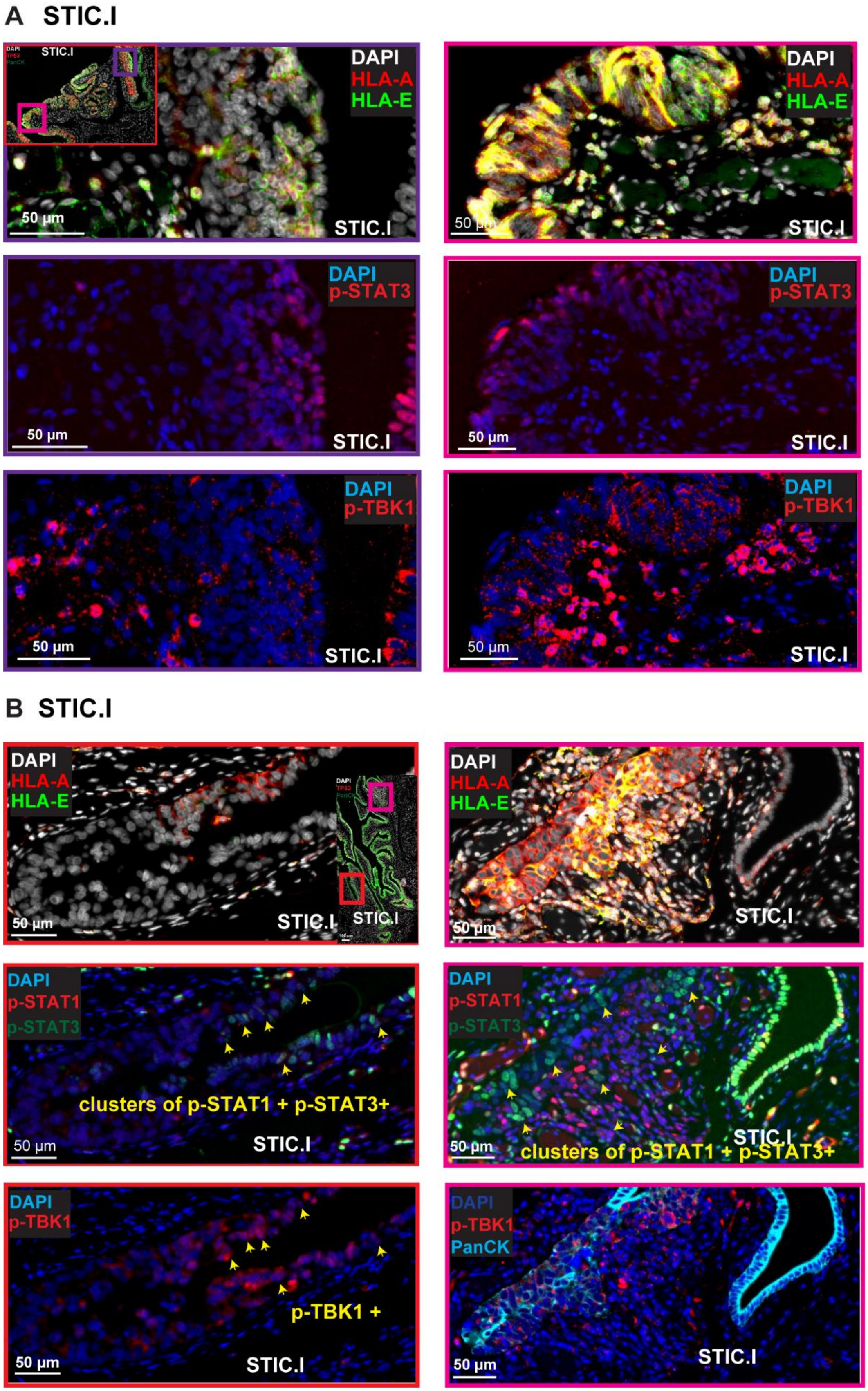

**Supplementary Fig. S7: Intra lesion heterogeneity of STIC.I:**

Even though incidental STICs overall overexpress HLA-A and HLA-E compared to matched FT.I, heterogeneity within the same STIC lesions has been observed. This figure depicts the intra-lesion heterogeneity of incidental STIC specimens in terms of the overexpression of genes downstream and upstream of IFN, such as HLA-A and HLA-E.

A. An example of STIC.I, outlined with a red box (case CD302.03(706), patient ID 38, BRCA1 Mut). The corresponding CyCIF image shows only STIC.I lesion, highlighting two areas of heterogeneity, outlined with purple and magenta boxes. The first region of STIC.I was highlighted with a purple box showing only some cells overexpressing HLA-A, HLA-E, p-STAT3 and p-TBK1. The second region of this STIC.I outlined with a magenta box suggestive of almost all cells over-expressing HLA-A, HLA-E, p-STAT3 and p-TBK1 (cytosolic/*punctate*).

B. Another example of STIC.I (case CD302.8(7923), patient ID 43, BRCA WT). The corresponding CyCIF image shows only STIC.I lesion, highlighting two areas of heterogeneity, outlined with red and magenta boxes. The first region of STIC.I, outlined with a red box shows very few cells overexpressing HLA-A, p-STAT3, p-STAT1 and p-TBK1. Cells co-expressing p-STAT1, p-STAT3 and p-TBK1 were showed with yellow arrow heads.

The second region of this STIC.I outlined with a magenta box showing almost all cells are HLA-A+, HLA-E+, both p-STAT3+ p-STAT1+ and p-TBK1+. Yellow arrow heads indicate cells co-expressing p-STAT1 and p-STAT3, indicative of cells with potential IFN activation. Overall, even though there was an increased median number of cells that showed a higher level of expressed proteins related to downstream of IFN activation, shown in Figure 3, such as HLA-A and HLA-E, heterogeneity for the presence of these proteins within the same lesion was observed.

**Supp Figure S8 (Related to Figure 3)**

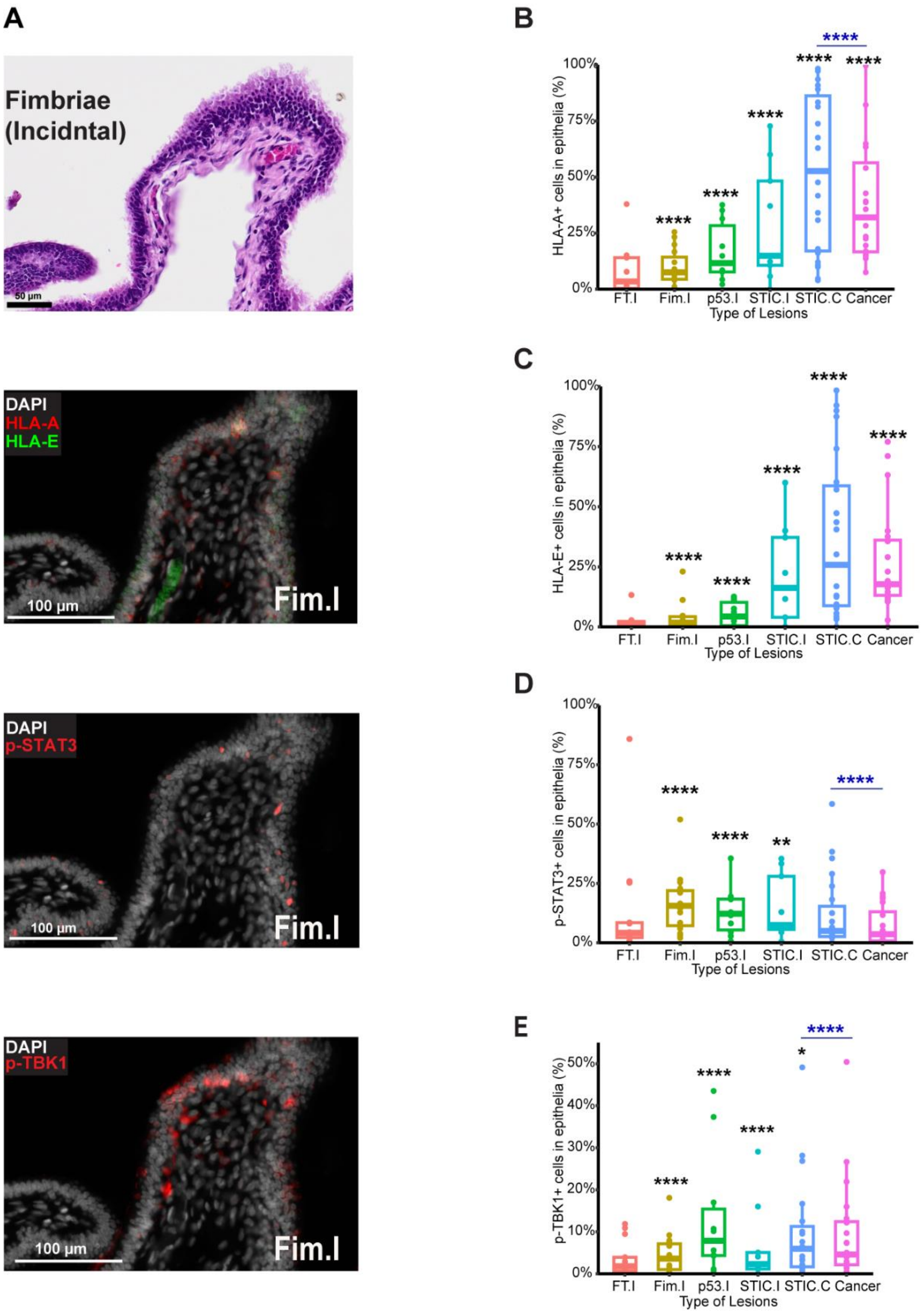

**Supp Figure S8. Single-cell analysis of multiplex tissue imaging related to IFN pathway activation:** Since IFN signaling pathway was upregulated even in p53.I stage, we validated few key genes upstream and downstream of IFN signaling pathway by multiple tissue imaging, CyCIF. A. CyCIF images representing downstream and upstream of IFN signaling pathway activation, such as overexpression of MHC-Class I (both HLA-A and HLA-E) and p-STAT3 in Fim.I. Very few, but when they express, mostly co-expressing HLA-A and HLA-E, indicating an upregulation of MHC-class I. This Fim.I case was matched from a p53.I case, shown in **Figure 3** (case C21-22 patient ID 28, BRCA1 Mut).

B-E. Box plots with individual data points show the comparison of (B) HLA-A+, (C) HLA-E+, (D) p-STAT3+, (E) p-TBK1+ epithelial cells across disease stages, from tissue imaging, expressed as a percentage. Number of specimens per group as follows: FT.I (n=13), Fim.I (n=15), p53.I (n=10), STIC.I (n=9), STIC.C (n=23) and Cancer (n=20). This data indicates an increased number of epithelial cells overexpressing potential downstream and upstream genes of IFN activation, especially at the initial stages of the disease. The solid line indicates the median within the interquartile range, with whiskers extending to a maximum of 1.5 times the interquartile range beyond the box. Black asterisks indicate significant differences in stages compared to the FT.I; blue asterisks indicate significant differences between groups; *$p<0.05$, ****$p<0.0001$. Binomial GLMMs taking patient ID as random effect (summary statistics in **Supplementary File S5, S6**).

**Supp Figure S9 (Related to Figure 3)**

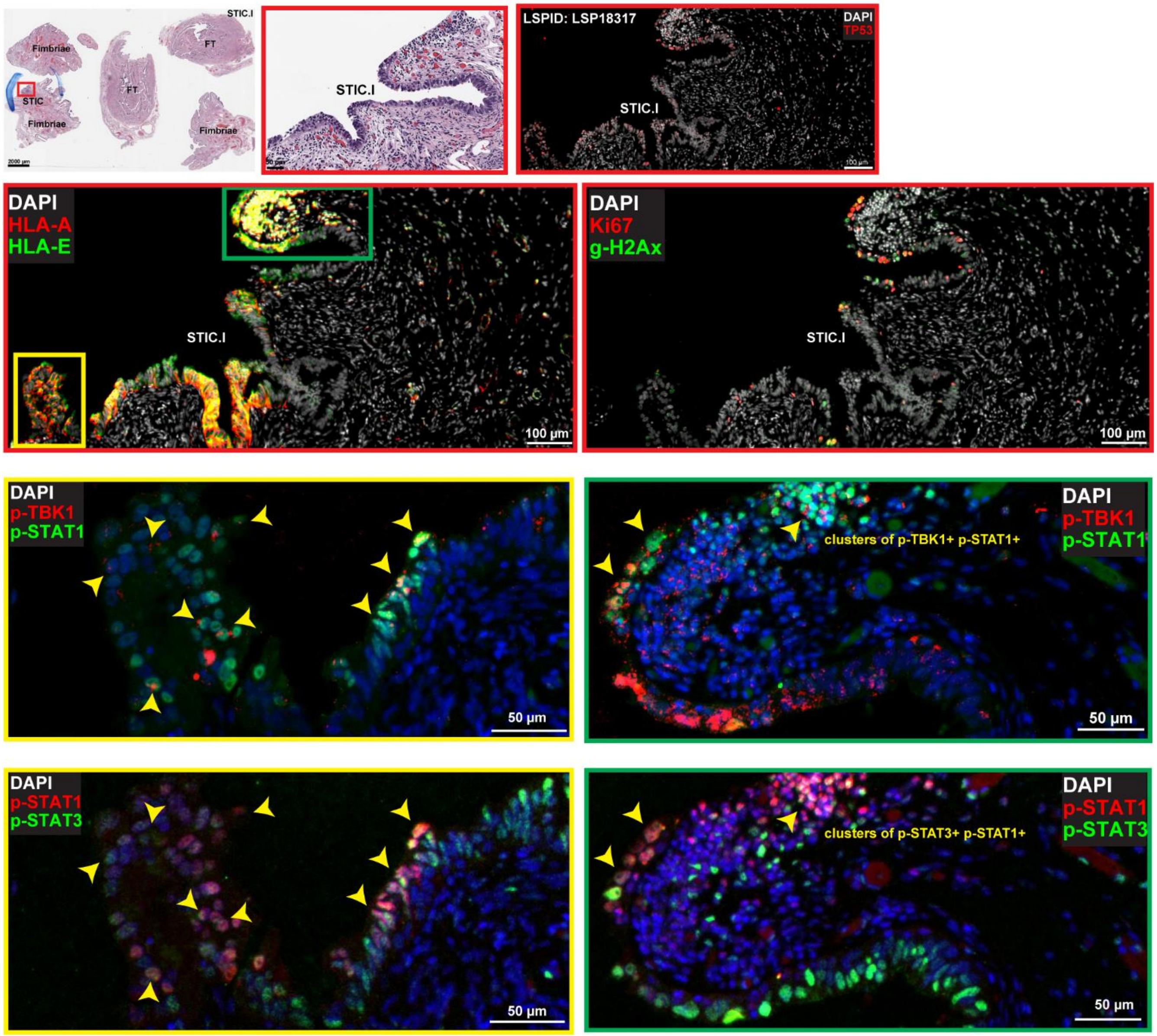


**Supp Figure S9.** A representation of an incidental STIC, outlined with a red box on H&E, (case CD302.05(964), patient ID 39, BRCA1 Mut, STIC.I). This CyCIF image shows an example of intra-lesion heterogeneity observed in terms of the IFN-activation and HLA-E overexpression followed by cells co-expressing p-STAT1 and p-STAT3 or p-STAT1 and p-TBK1 (cytosolic/*punctate*). For instance, some regions of the same STIC have no IFN activation (no HLA-A/E expression). Co-expression of these markers on the same cell was shown with yellow arrowheads. These areas were outlined with a yellow and green box. IFN pathway seems to be activated in the epithelium and some adjacent stromal components.

**Supp Figure S10 (Related to Figure 3)**

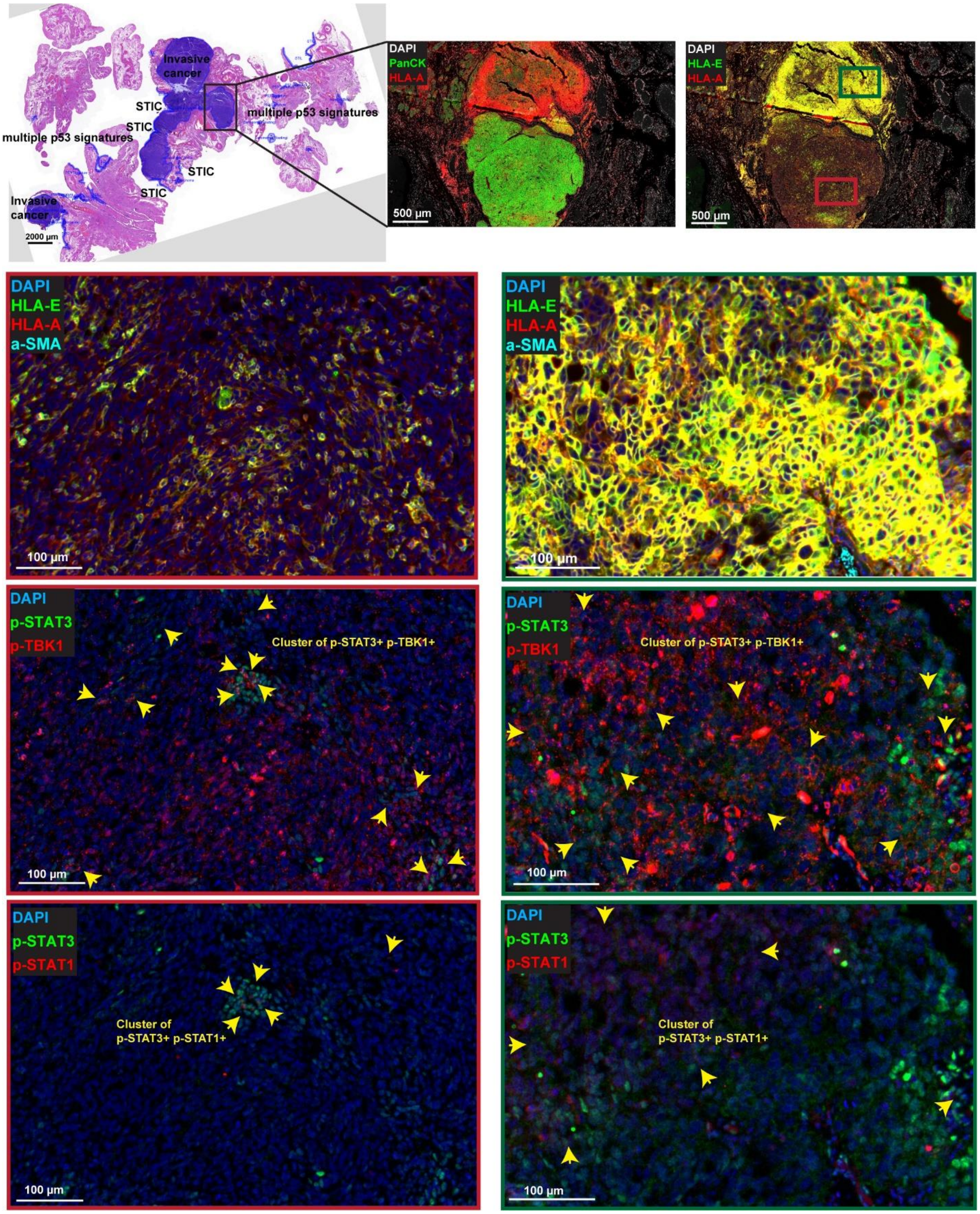

**Supp Figure S10.** A representation of an invasive cancer co-existing with STIC (case RD-23-005, patient ID 12, BRCA1 Mut, Stage IC HGSOC). The heterogenous tumor clones were shown on H&E with a black outline box. This CyCIF image depicts an example of heterogeneity observed in terms of the intensity of HLA-E overexpression followed by cells co-expressing p-STAT1 and p-STAT3 or pSTAT3 and p-TBK1 (cytosolic/*punctate*) (yellow arrows). Co-expression of downstream or upstream of potential activation of IFN- related genes are more frequently observed in the clones with high HLA-E (i.e. most cells expressing HLA-E with a high intensity), outlined with a green box and vice versa (red box). This figure represents intra tumoral heterogeneity in terms of IFN pathway.

**Supp Figure S11 (Related to Figure 3)**

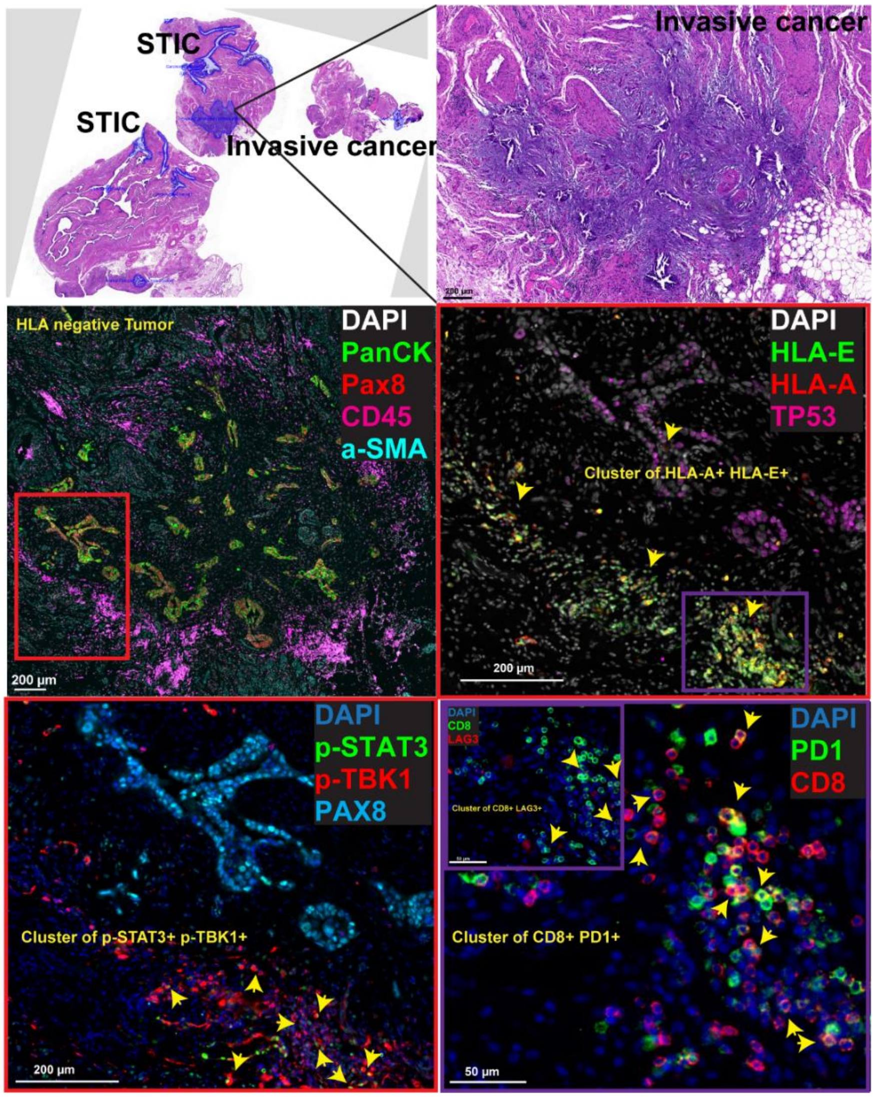


**Supp Figure S11.** A representation of invasive cancer co-existing with STIC (case RD-23-008, patient ID 14, BRCA WT, Stage IIIC HGSOC). This CyCIF image depicts an example of "inter" patient heterogeneity observed in the HLA-E overexpression followed by cells co-expressing pSTAT3 and p-TBK1 (cytosolic/*punctate*). This is the sole tumor specimen whereby, in the epithelium, almost no HLA-A and HLA-E expression was observed, meaning HLA-E negative tumor. No p-TBK1 or p-STAT3 expression was observed in the epithelium either, although cGAS was shown to be colocalized with BAF+ MN ruptured DNA in this area (not shown). This may suggest this particular tumor regulates IFN pathway downstream of cGAS-BAF colocalization. The IFN pathway seems to be activated only in the adjacent stroma of the tumor epithelium (i.e., tumor margin). The adjacent stroma area with HLA-E expression was outlined with a purple box. Expression of LAG3+ or PD1+ on CD8+ T cells was observed and shown with yellow arrows. This figure represents inter-patient heterogeneity regarding IFN pathway activation in the tumor context. STIC.C in this specimen has both HLA-E positive and negative epithelium (not shown).

**Supp Figure S12 (Related to Figure 3)**

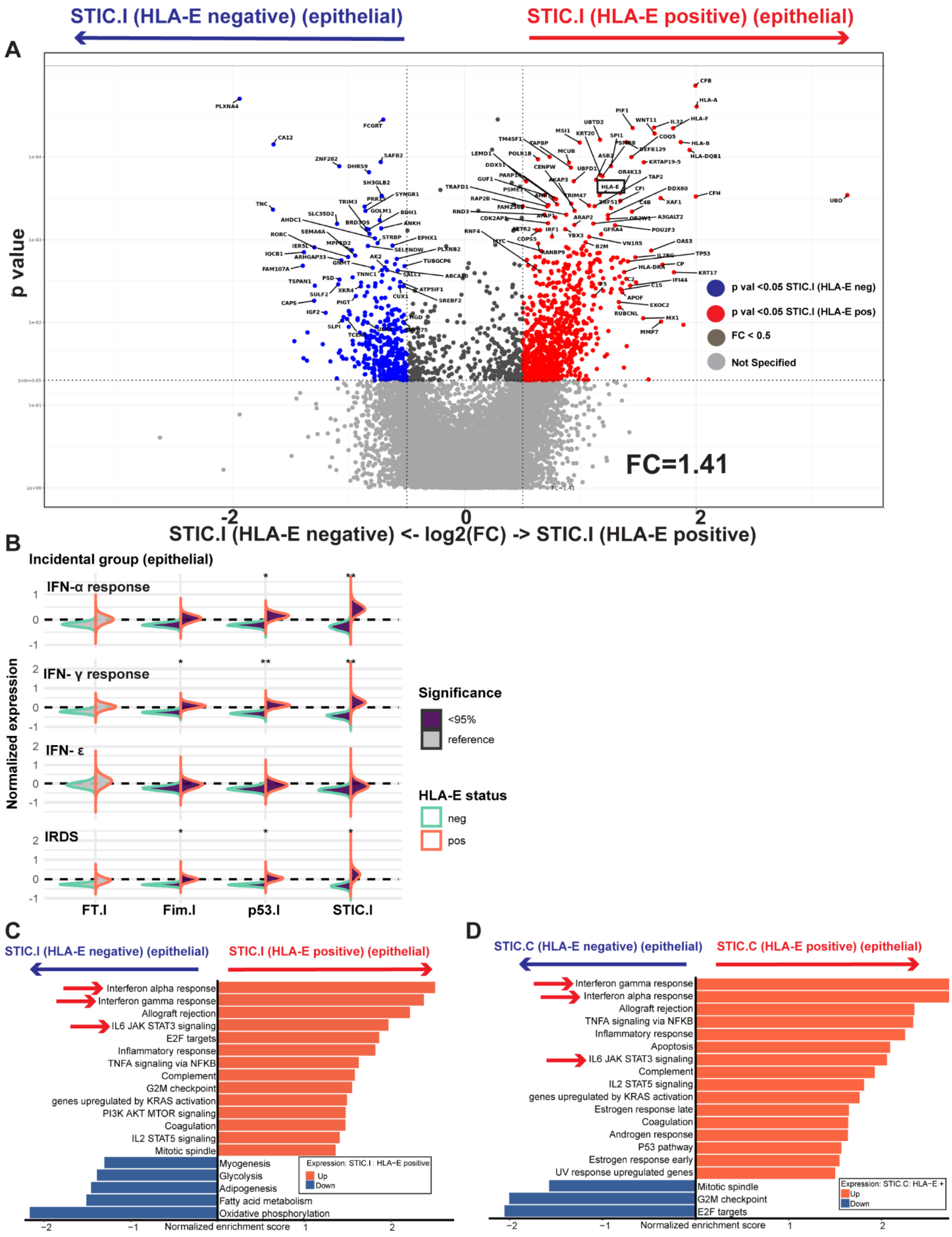

**Supp Fig S12: IFN activation was predominantly observed in HLA-E positive cells.**

Since we have observed heterogeneity for protein expression related to or indicative of IFN pathway activation, such as HLA-E within the same STIC lesions and cancer, we have collected both HLA-E positive (+) cells and HLA-E negative (-) cells for spatial transcriptomics, as shown in Supplementary Fig.S2.

A. Differential gene expression between HLA-E + and HLA-E - clones of STIC.I in epithelial compartments. Considering that overall, more HLA-E+ epithelial cells were observed in tissue imaging with disease progression, the number of ROIs or epithelial segments analyzed here were 9 (HLA-E -) and 18 (HLA-E +). Volcano plot showing HLA-E + cells observed by tissue imaging also overexpress *HLA-E* in RNA level compared to HLA-E – cells (outlined with a black box). HLA-E + cells were also predominantly associated with genes related to but not limited to MHC-class I, such as *HLA-A, HLA-B*; the response of IFN-α and IFN-ɣ, such as *MX1*, cell growth, such as *MYC*. FC, fold change. Linear Mixed Modeling, taking patient ID as a random effect were used for differential gene expression.

B. Bayesian modeling taking patient ID as a random effect and HLA-E status (+ or - cells observed by tissue imaging) as a fixed effect confirmed the association of upregulation of selected genes related to the response of IFN-α and IFN-ɣ and IRDS, only in HLA-E + cells even at the stage of Fimbriae of the incidental group. To investigate the effect of HLA-E expression on other genes, we further modified the Bayesian model, shown in **Figure 2F**, to include a fixed effect coefficient serving as a binary indicator for the presence or absence of HLA-E in the CyCIF image of the ROI (gene_expression ~ mo(stage) * hlae_e + (1 + mo(stage) * hlae_e | patient_id * gene). Columns correspond to types of lesions of incidental group, and rows correspond to gene sets consisting of selected genes of IFN Hallmark set or IRDS. Bayesian modeling was applied to see the trend of any changes in gene expression in incidental precursor lesions compared to the matched FT.I (reference) as well as within the same group between HLA-E + (orange colored border) and HLA-E- (green color bordered) cells. Significance testing used the proportion of the 95% highest density interval (HDI) within the Region of Practical Equivalence (ROPE, 0.05 times the standard deviation). Comparisons with >95% of the HDI outside the ROPE were significant (*); >99% were very significant (**) (See Methods for the details of the modeling, **Supplementary Figure S13**). The same test was applied either comparing to FT.I (reference) or comparing HLA-E + and HLA-E – cells within the same group.

C-D. GSEA was performed on STIC.I and STIC.C. MsigDB Cancer Hallmark gene sets confirmed the association with HLA-E + cells namely response in IFN-α and IFN-ɣ, inflammation such as IL-6 JAK STAT3 pathway and TNF- α signaling via NFKB. Some of these IFN-related pathways are shown with red arrows. Ranking of the pathways was based on adjusted p-value <0.05. C. The number of STIC.I ROIs or epithelial segments analyzed here were 9 (HLA-E -) and 18 (HLA-E +). D. The number of STIC.C ROIs or epithelial segments analyzed here were 28 (HLA-E -) and 68 (HLA-E +).

**Supp Figure S13 (Related to Figure 2, 3)**

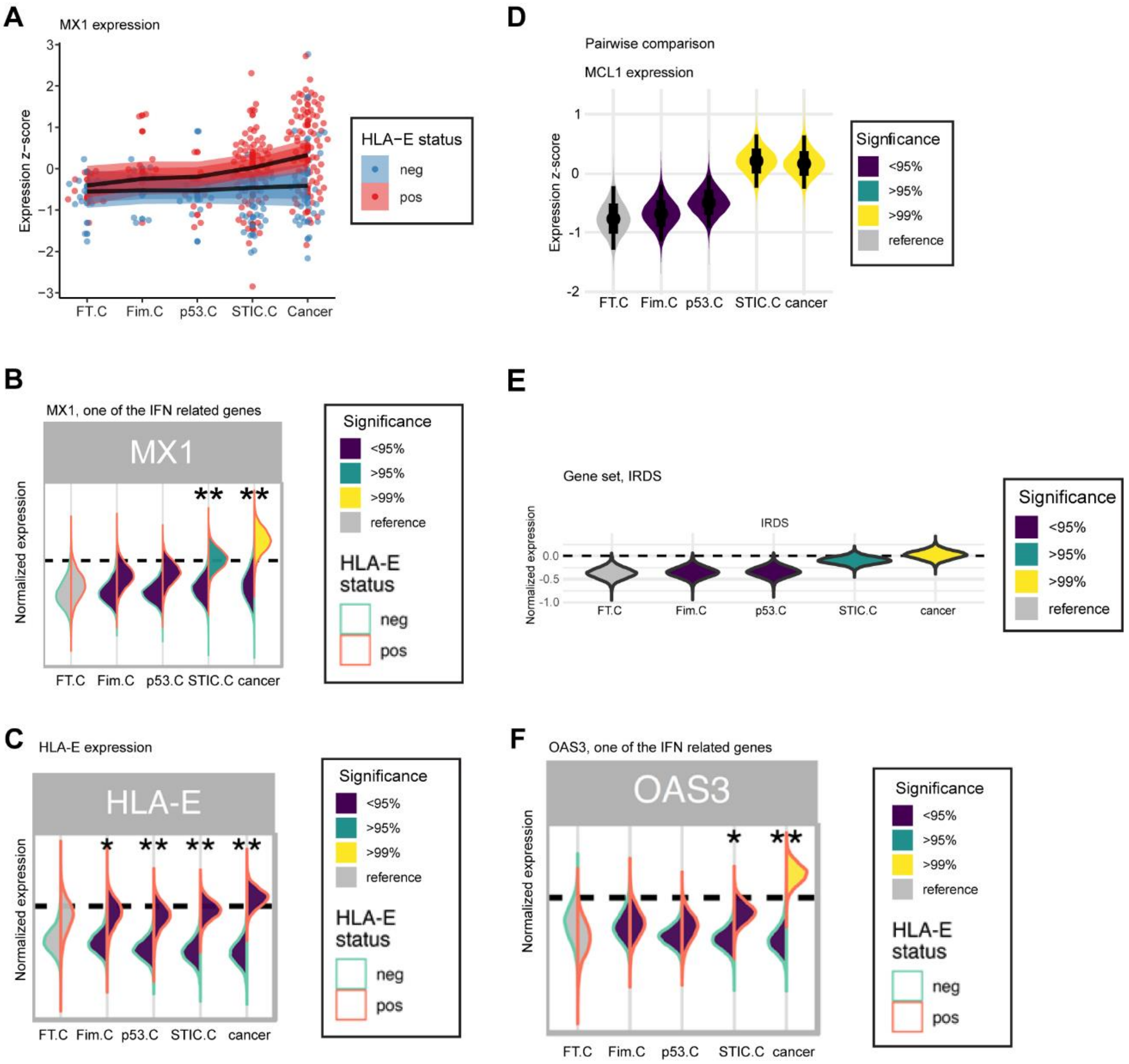


stage parameter (average expression change between two stages):
**-0.05** [ -0.20, 0.09] 95% CI **p 0.24**

hlae parameter (average difference between HLA-E positive vs negative):
**0.42** [ 0.12, 0.98] 95% CI **p 0.03**

**G brms model using HLA-E as a fixed effect**

```
model <- brm (
expression ~ mo(stage) *hlae + (1 + mo (stage) * hlae | patient * gene),
data = input_data,
prior = c(
   set_prior ( “normal (0,2) ”, class = “b”),
   set_prior ( “normal (0, 2) ”, class = “Intercept”)
)
)
```

**Supp Figure S13. Bayesian regression model with further modification:**

A. To investigate the effect of HLA-E expression on other genes, we modified the Bayesian model to include a fixed effect coefficient serving as a binary indicator for the presence or absence of HLA-E in the CyCIF image of the ROI. Using ordinal effect model, Bayesian regression modeling was applied to our own dataset, *MX1* from cancer group, as an example across stages of the disease.

B. Many IFN-related genes are sensitive to HLA-E status, such as *MX1*. Each column corresponds to each stage of the disease.

C. Applying the modified model confirms that the protein and RNA levels of HLA-E match, meaning HLA-E positive ROIs in the CyCIF image have higher HLA-E RNA expression than HLA-E negative.

D. Pairwise expression is possible using an example of *MCL1* from our dataset.

E. Progression of individual genes as a gene set is possible, such as IRDS.

F. Many IFN-related genes are sensitive to HLA-E status, such as *OAS3*.

B-F. Significance testing used the proportion of the 95% highest density interval (HDI) within the Region of Practical Equivalence (ROPE, 0.05 times the standard deviation). Comparisons with >95% of the HDI outside the ROPE were significant (*); >99% were very significant (**).  Matched FT.C was chosen as a reference.

G. In summary, to investigate the effect of HLA-E expression on other genes, we further modified the model: (gene_expression ~ mo(stage) * hlae_e + (1 + mo(stage) * hlae_e | patient_id * gene).

### Supp Figure S14 (Related to Figure 3)

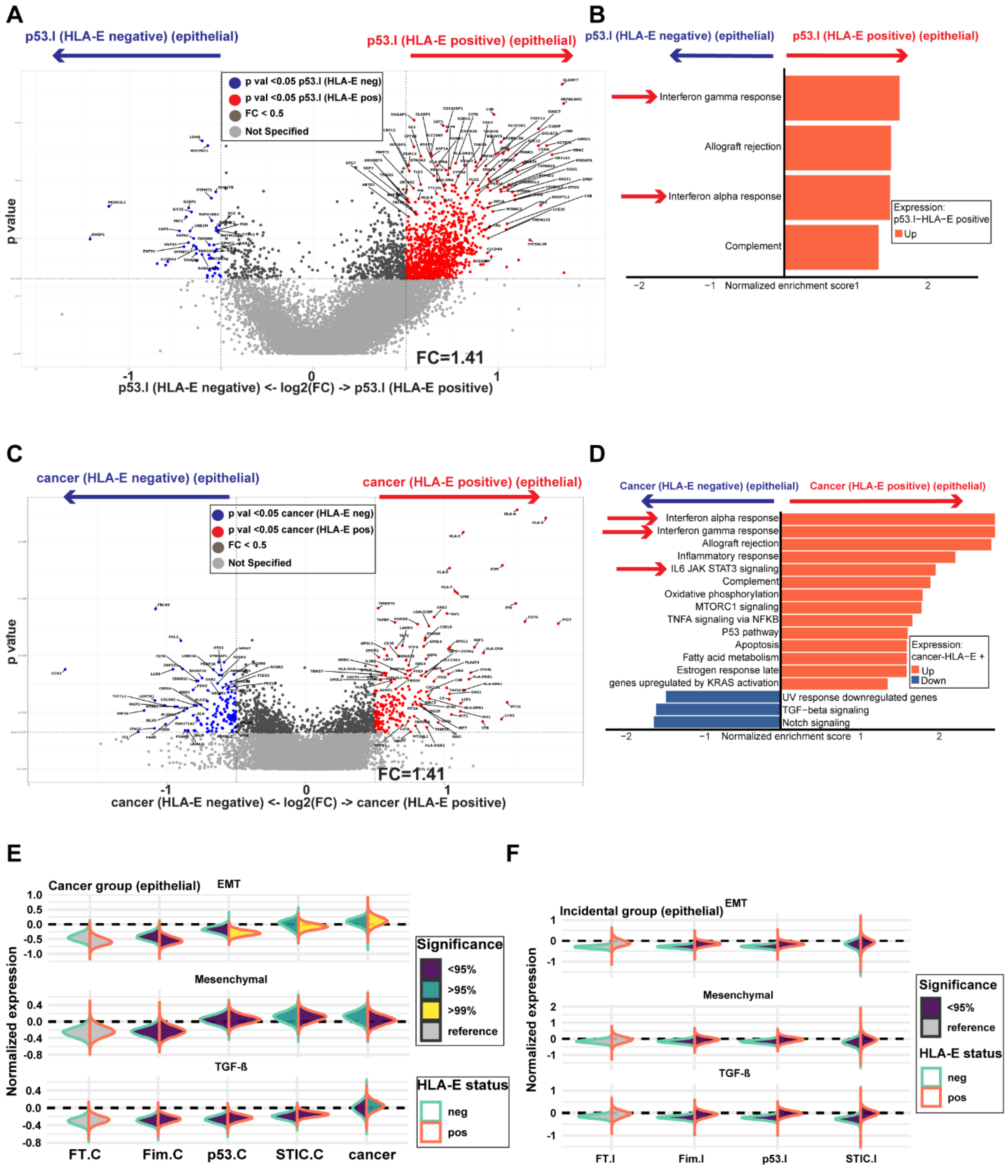

**Supp Figure S14. Association of HLA-E with IFN and other cancer hallmark pathways.** Differential gene expression and GSEA on MsigDB cancer hallmark pathways were performed for HLA-E positive and negative clones of the epithelium of p53.I (A-B), cancer (C-D). The number of p53.I ROIs or epithelial segments analyzed here were 24 (HLA-E -) and 15 (HLA-E +). The number of carcinoma ROIs or epithelial segments analyzed here was 27 (HLA-E -) and 78 (HLA-E +). IFN related pathways were shown with red arrows.

E-F. Bayesian modeling was performed in the epithelia of the cancer group or incidental group to see the association of HLA-E and genes related to selective genes of EMT or mesenchymal or TGF-β pathway as a gene set. Bayesian modeling was applied, taking patient ID as random effect but HLA-E status from tissue imaging as a fixed effect (**Supplementary Fig S13**).

**Supp Figure S15 (Related to Figure 3)**

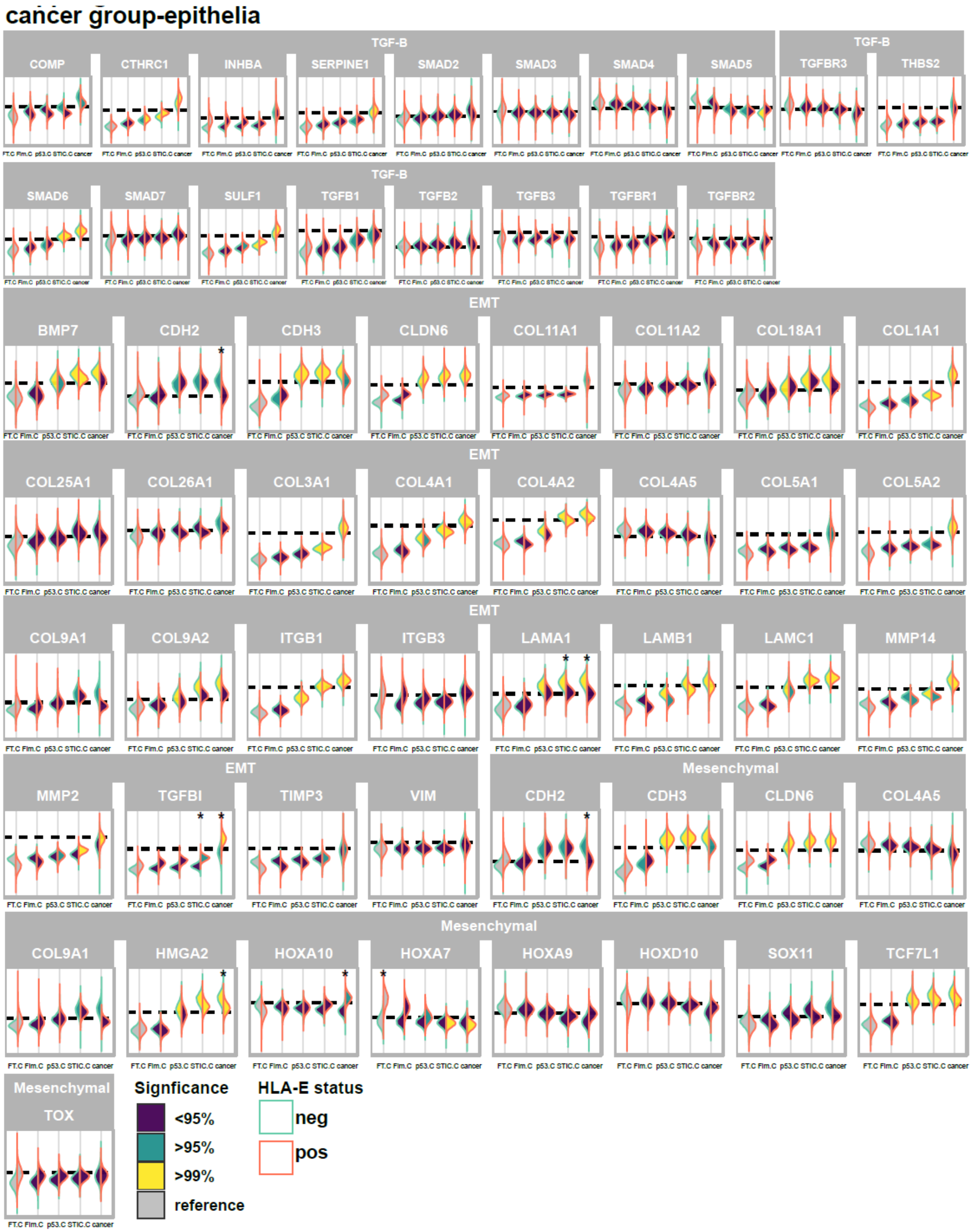

**Supp Figure S15.** Selected gene set from HGSOC cohort (published), genes related to TGF-beta, Epithelial mesenchymal transition (EMT) and mesenchymal phenotype (C5/PRO subtype) of HGSOC are not dependent on HLA-E status from CyCIF, meaning that they are potentially independent of IFN signaling pathway. Bayesian modeling was applied to the individual gene list, not as a gene set, taking patient ID as random effect but HLA-E status from tissue imaging as a fixed effect (**Supplementary Figure S13**). Y axis is normalized expression.

**Supp Figure S16 (Related to Figure 4)**

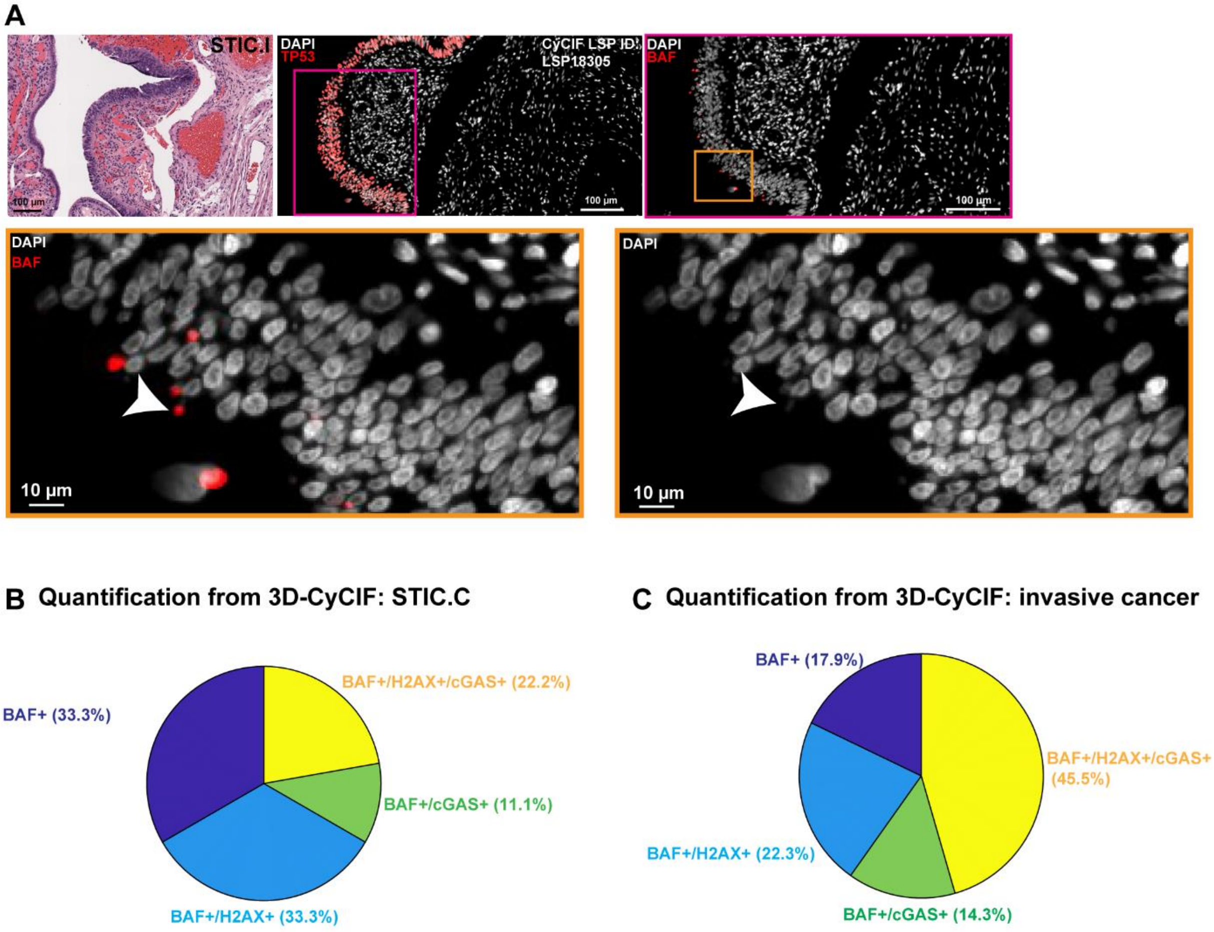


**Supp Figure S16. Multiplex tissue imaging revealed that micronucleus rupture might be one of the mechanisms of IFN signaling pathway activation through cGAS-STING pathway:** Since one of the potential upstream activators of the IFN pathway was p-TBK1, meaning cGAS-STING pathway might be activated at the precancer lesions. BAF (*BANF*), a marker of Micronuclei (MN) rupture, potentially indicates the presence of cytosolic DNA due to MN rupture, which may activate cGAS-STING pathway. A. Top: H&E of a representative case of STIC.I, also shown in Figure 3 (case CD302.04(939), patient ID 40, BRCA WT). CyCIF imaging showing BAF positive MN rupture, outlined with an orange box. Bottom: CyCIF image showing with white arrow head indicates the BAF+MN rupture. DAPI (i.e. DNA) only staining confirms the micronucleus in regions of STIC.I outlined with an orange box.

B-C. Pie charts showing the number of BAF+ MN or BAF+cGAS+ɣ-H2Ax+ MN or BAF+cGAS+ MN in STIC.C and invasive components from 3D reconstruction of a case shown in Figure 4, expressed as percentage.

**Supp Figure S17 (Related to Figure 5)**

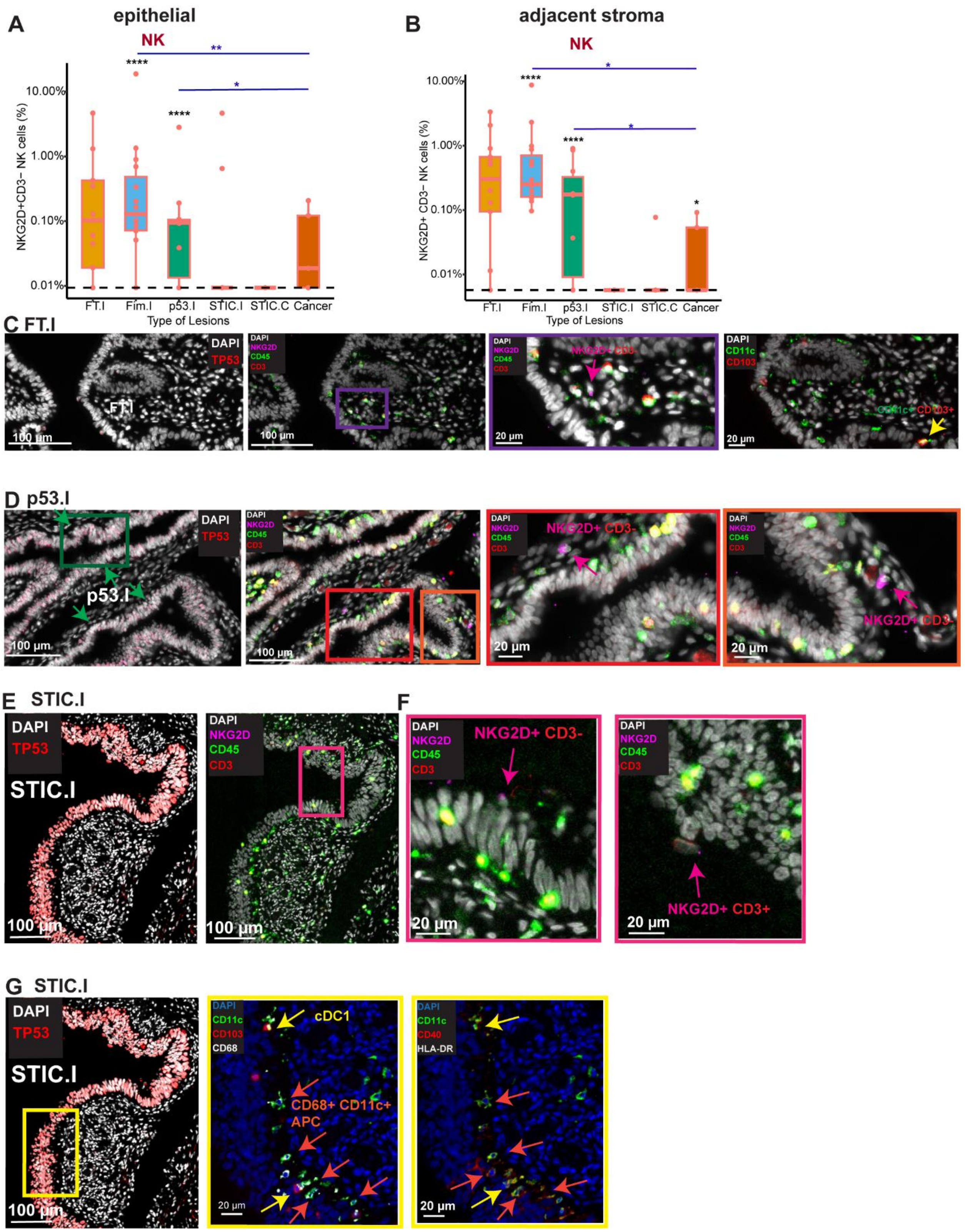

**Supp Fig. S17: Lower abundance of NK cells with HGSOC progression:**

A-B. Box plots with individual data points showing the NK cells observed across disease stages in the epithelium (A) and in the adjacent stroma (B), expressed as a percentage. Number of specimens per group for NKG2D antibody as follows: FT.I (n=13), Fim.I (n=15), p53.I (n=10), STIC.I (n=9), STIC.C (n=7) and Cancer (n=5). Black asterisks indicate significant differences in stages compared to the FT.I and blue lines with asterisks indicate comparison between groups; *$p<0.05$,****$p<0.0001$, binomial Generalized Linear Mixed-effects Models (GLMM). Y axis is represented as log10 scale, the dashed line represents values with zero.

C. CyCIF image showing from a representative case of a FT.I from a matched STIC.I, also shown in Figure 3 (case CD302.04(939), patient ID 40, BRCA WT). The Magenta arrowhead indicates an NKG2D+CD45+ CD3- immune cell in the adjacent stroma, suggestive of an NK cell. Yellow arrowhead indicates a CD11c+ CD103+ immune cell close to the epithelium, suggestive of a cDC1, potentially acting as the primary antigen presenting cells for early presentation to naïve CD4+T cells.

D. CyCIF image showing from a representative case of a p53.I, also shown in Figure 3 (case C21-22 patient ID 28, BRCA1 Mut). Green box and green arrows on p53.I indicate the layer of epithelial cells representing "p53signatures". Magenta arrowheads in 2 regions (orange and red box) indicate an NKG2D+CD45+ CD3- immune cell in the adjacent stroma close to the epithelium, suggestive of the presence of NK cells in the "p53 signatures".

E-F. CyCIF image showing a representative case of STIC.I, also shown in Figure 3 (case CD302.04(939), patient ID 40, BRCA WT). CyCIF images outlined with magenta box representing both NK cell (NKG2D+ CD45+ CD3-) shown with magenta arrowheads. Just to compare NKT cell (NKG2D+ CD45+ CD3+) was also shown with magenta arrow heads.

G. The Yellow box (G) represents activated APC (CD40+), in this case both cDC1 (CD11c+ CD103+ CD68-) (shown with yellow arrows) and CD68+ macrophage-derived APCs (CD11c+ CD103- CD68+) (shown with orange arrows) on the same STIC.I specimen, shown in panel E.

### Supp Figure S18 (Related to Figure 5)

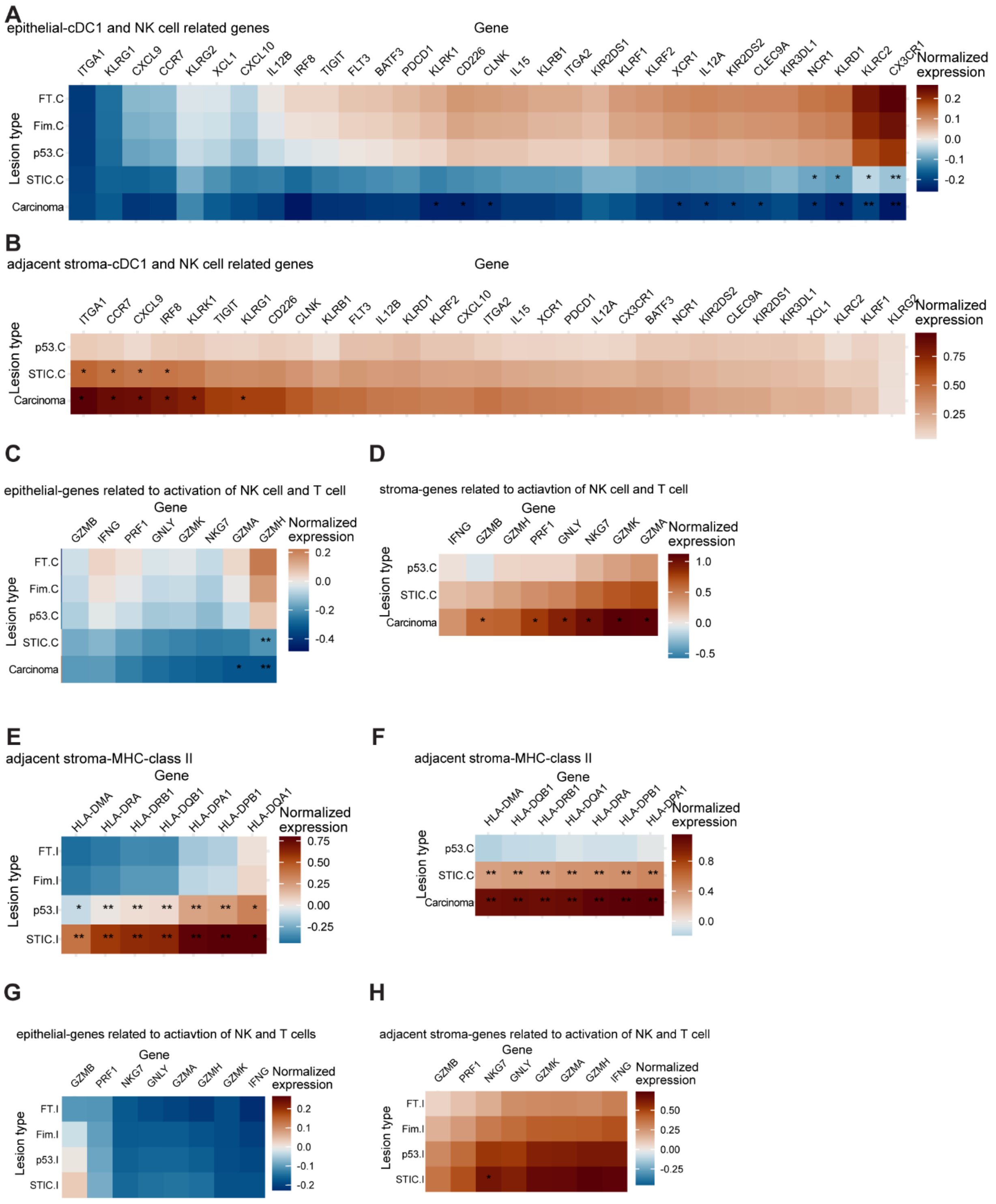

**Supp Fig S18: Expression of genes related to NK-cDC1 axis during HGSOC progression:**

A-D. Heatmaps showing normalized expression of selected published genes in the epithelia (A, C) and adjacent stroma (B,D) related to cDC1 and receptors of NK cells (A-B) as well as degranulation/activation of NK and T cells (C-D) from cancer group compared to their matched FT.C (A,C) or p53.C (B,D). Due to the unavailability of sufficient ROIs of stroma from FT.C/Fim.C in the cancer group, relative expression was compared with the stroma adjacent to p53.C. Significant downregulation of a few genes related to NK cell activation and inhibitory receptors and cDC1 followed by degranulation or activated NK/CD8+ T cells only observed in the tumor epithelium. Columns correspond to individual genes, and rows correspond to types of lesions.

E-F. Heatmaps showing normalized expression of genes in the adjacent stroma related to MHC-class II from incidental (E) and cancer groups (F) compared to their matched FT.I and p53.C, respectively from spatial transcriptomics.

G-H. Heatmaps showing normalized expression of selected published genes in the epithelia (G) and in the adjacent stroma (H) of incidental group related to degranulation/cytotoxicity of NK and T cells from cancer group compared to their matched FT.I. A trend or increased in few genes including *GZMB, IFNG, NKG7* was observed in both epithelium and adjacent stroma. Columns correspond to individual genes, and rows correspond to types of lesions.

A-H. Bayesian modeling was applied to see the relative changes in gene expression in either the incidental or cancer group compared to the matched FT or p53 signature (only in tumor-stroma). Median of the posterior distribution was shown in heatmaps. Significance testing used the proportion of the 95% highest density interval (HDI) within the Region of Practical Equivalence (ROPE, 0.05 times the standard deviation). Comparisons with >95% of the HDI outside the ROPE were significant (*); >99% were very significant (**).

**Supp Figure S19 (Related to Figure 5)**

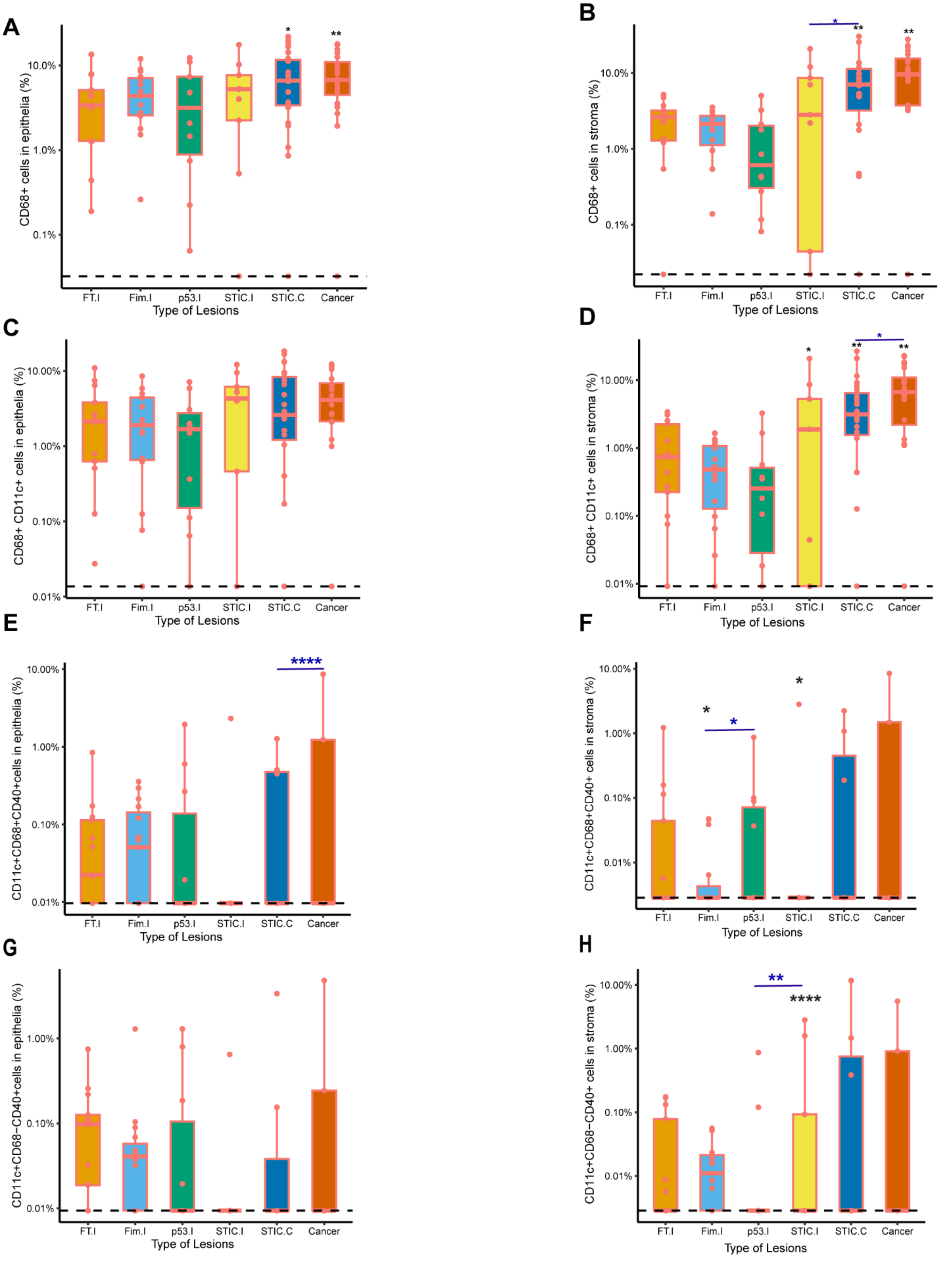

**Supp Figure S19**: **Increased macrophage and macrophage derived APC populations with HGSOC progression:**

A-H. Box plots with individual data points showing CD68+ macrophages, CD68+CD11c+, representing potential macrophage derived APCs, activated macrophage derived APC cells (CD11c+ CD68+ CD40+) (CD40 antibody was used only on a subset of cases) and activated dendritic cells (CD11c+ CD68- CD40+), expressed as a percentage. Black asterisks indicate significant differences in stages compared to the FT.I and blue lines with asterisks indicate comparison between groups; *$p<0.05$,****$p<0.0001$, binomial Generalized Linear Mixed-effects Models (GLMM). Y axis is represented as log10 scale, the dashed line represents values with zero.

A-D. Number of specimens per group as follows: FT.I (n=13), Fim.I (n=15), p53.I (n=10), STIC.I (n=9), STIC.C (n=23), and Cancer (n=20).

E-H. Number of specimens per group for CD40 antibody as follows: FT.I (n=13), Fim.I (n=15), p53.I (n=10), STIC.I (n=9), STIC.C (n=7) and Cancer (n=5).

**Supp Figure S20 (Related to Figure 5)**

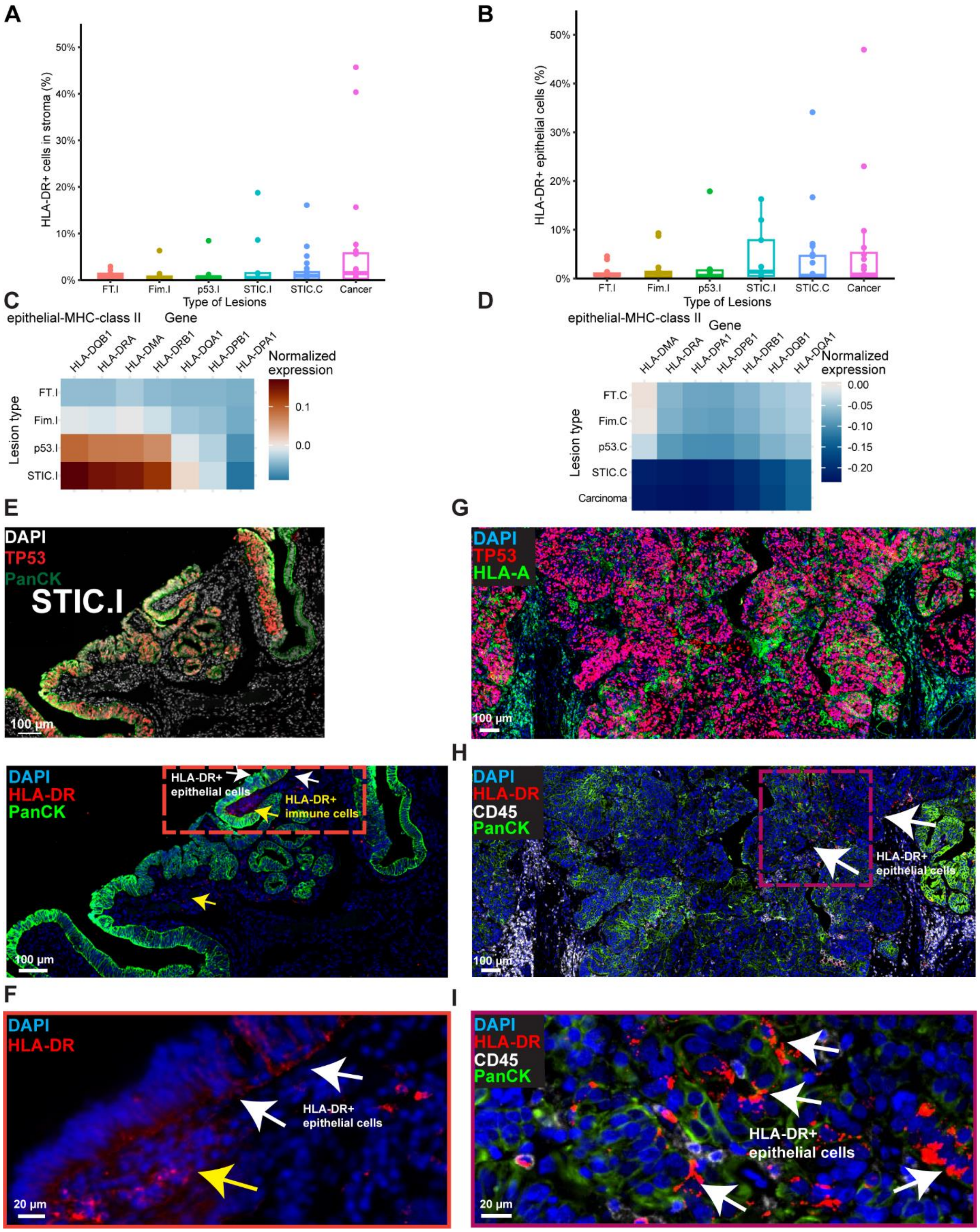

**Supp Figure S20. MHC-II class expression in HGSOC:**

A-B. Box plot shows the comparison of HLA-DR+ cells in stroma (A) or in the epithelia (B) across disease stages, from tissue imaging, expressed as a percentage. Number of specimens per group as follows: FT.I (n=13), Fim.I (n=15), p53.I (n=10), STIC.I (n=9), STIC.C (n=23), and Cancer (n=20). FT.I from the incidental group was chosen as a control group. The solid line indicates the median within the interquartile range, with whiskers extending to a maximum of 1.5 times the interquartile range beyond the box. Outliers were shown as dots. Black and blue asterisks indicate significant differences in stages compared to the FT.I and between groups, respectively. Binomial GLMMs, with patient ID and observational level random effect.

C-D. Bayesian modeling on genes related to MHC-class II on epithelium compared to their matched FT.

E-I. A representative CyCIF image for STIC.I case (case CD302.03(706), patient ID 38, BRCA1 Mut) (E), suggests very few HLA-DR+ epithelial cells outlined with an orange box. White arrows indicate HLA-DR+ epithelial cells and yellow arrow indicates HLA-DR+ immune cells (F). G-I. A representative CyCIF image of a tumor, also shown in **Fig. 1F**, (Case RD-23-002, patient ID 9, BRCA2 mutant, Stage IC HGSOC) (G). Some HLA-DR+ epithelial cells were observed, outlined with a magenta box (H). White arrows indicated HLA-DR+ (MHC-class II) epithelial cells (I). In comparison, HLA-A+ (MHC-class I) tumor epithelial cells were shown in G.

**Supp Figure S21 (Related to Figure 5)**

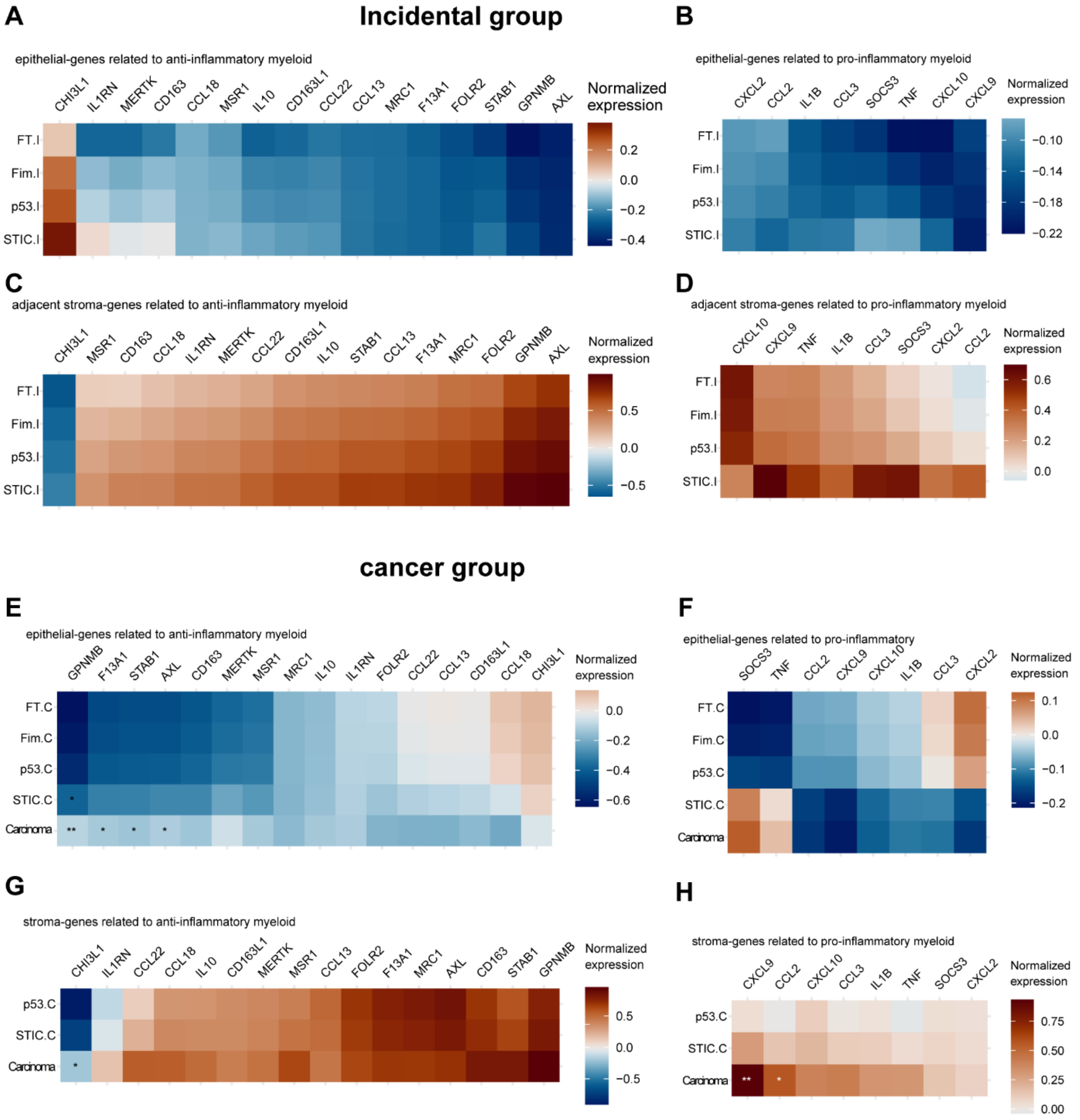

**Supp Figure S21. Expression of genes related to pro-inflammatory and anti-inflammatory myeloid populations across disease stages:**

A-D. Heatmaps showing expression of the genes related to anti-inflammatory and pro-inflammatory myeloid populations in incidental cases in the epithelial (A-B) and the adjacent stroma (C-D).

E-H. Heatmaps showing expression of the genes related to anti-inflammatory and pro-inflammatory myeloid populations in cancer group in the epithelial (E-F) and the adjacent stroma (G-H).

A-H. Columns correspond to individual genes, and rows correspond to types of lesions. Bayesian modeling was applied to see the relative gene expression changes in the incidental or cancer group compared to the matched FT or p53.C (stroma-cancer group). Median of the posterior distribution was shown in heatmaps. Significance testing used the proportion of the 95% highest density interval (HDI) within the Region of Practical Equivalence (ROPE, 0.05 times the standard deviation). Comparisons with >95% of the HDI outside the ROPE were significant (*); >99% were very significant (**).

**Supp Figure S22 (Related to Figure 6)**

A

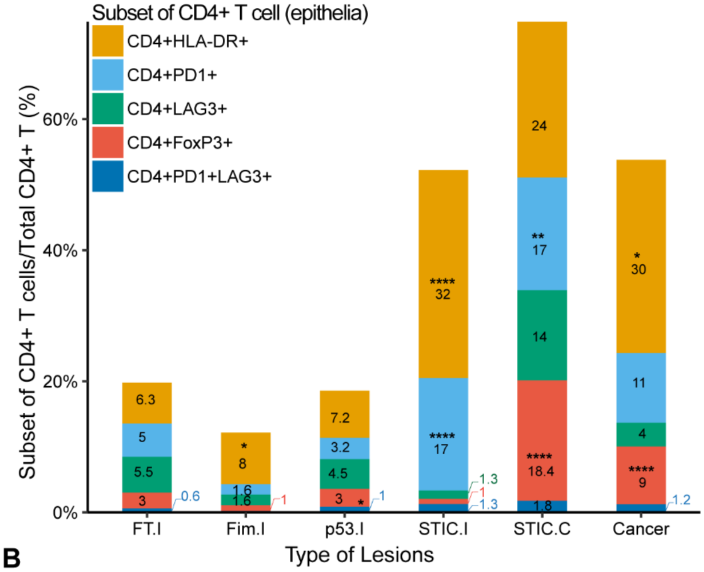


B

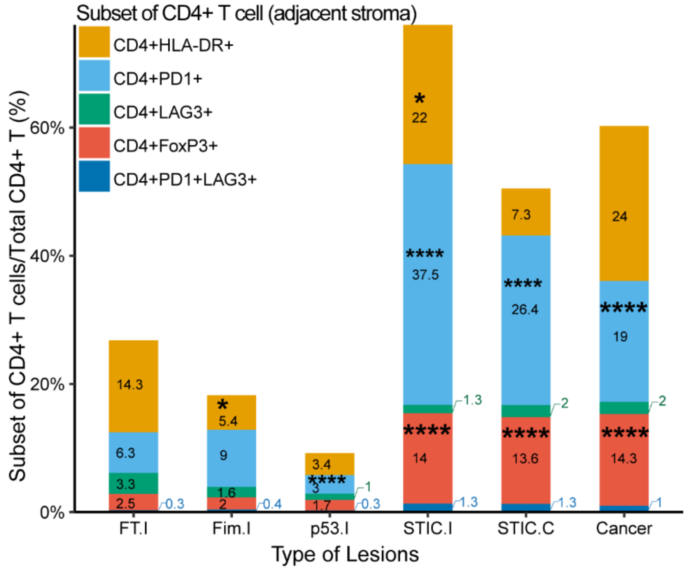


**Supp Figure S22. Increased CD4+ T cell infiltration during HGSOC progression:**

A-B. Stacked bar plots showing the proportion of CD4+ T cells in total CD4+ T cells from single cell-CyCIF analysis in the epithelia (A) and in the adjacent stroma (B) across disease stages. Number of specimens per group as follows: FT.I (n=13), Fim.I (n=15), p53.I (n=10), STIC.I (n=9), STIC.C (n=23) and Cancer (n=20). Black asterisks indicate significant differences in stages compared to the FT.I; *p<0.05, **p<0.01, ****p<0.0001. Binomial GLMMs, with patient ID as random effect. Average proportions were rounded up to the next whole number when applicable and shown for each cell state across lesion types.

**Supp Figure S23 (Related to Figure 5)**

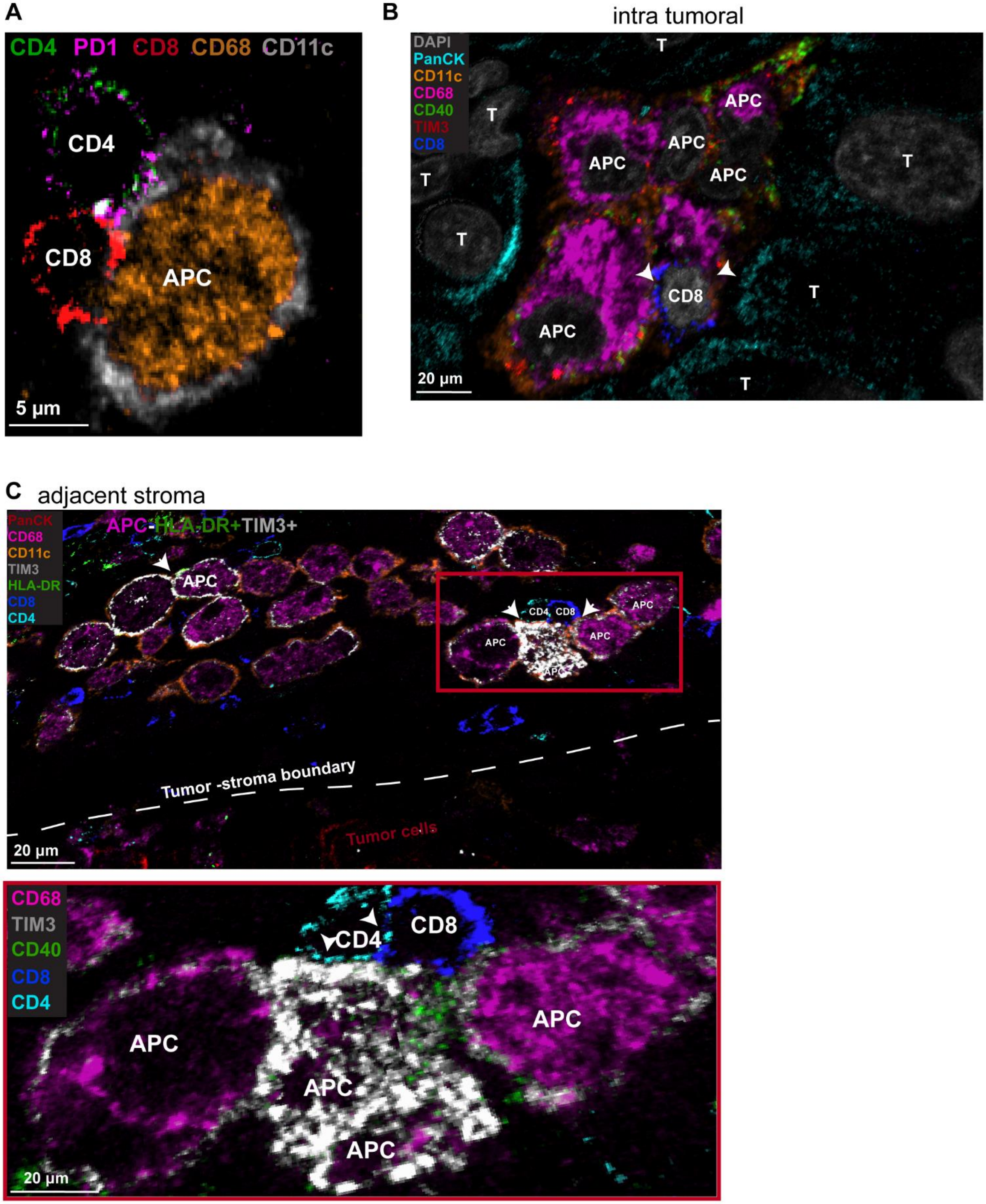

**Supp Fig. S23: Interactions between antigen presenting cells (APCs) and T cell in 3D CyCIF:**

3D reconstruction of a tumor from a case of STIC with concurrent cancer is shown in **Figure 1F**, (Case RD-23-002, patient ID 9, BRCA2 mutant, Stage IC HGSOC). One of the major immune cell subtypes found in 2D CyCIF was macrophage derived APCs (CD11c+ CD68+), increasing with HGSOC. These APCs often expressing HLA-DR, indicative of their antigen presentation function (activated).

A. By using 3D CyCIF, we can visualize the potential interactions between different immune cell populations. For instance, CD11c+CD68+ APCs are found to be clustered together with CD4 and CD8, indicating a potential interaction shown in (A). We also observed CD4+ T cells often express PD1 (activated CD4+) while interacting with CD11c+CD68+ APCs (A). These interactions were observed intra-tumoral and in the tumor stroma boundary (adjacent stroma).

B-C. Visualization with TIM3 antibody suggested most of these APCs co-expressing HLA-DR/CD40 (potential activation) and TIM3, indicative of reduced functionality of APCs in intra tumoral (B) or in the adjacent stroma (C). These APCs were observed to be either isolated or clustered together with CD8 and CD4 outlined with the red box in the adjacent stroma. We can see a potential interaction of CD8+ and CD4+ T cells with APCs, as indicated in white arrows. Two out of four APCs expressing co-stimulatory molecules interacting with the T cells. However, all of these APCs (CD68+ CD11c+) express TIM3, indicating potential reduced functionality or dysfunction.

**Supp Figure S24 (Related to Figure 6)**

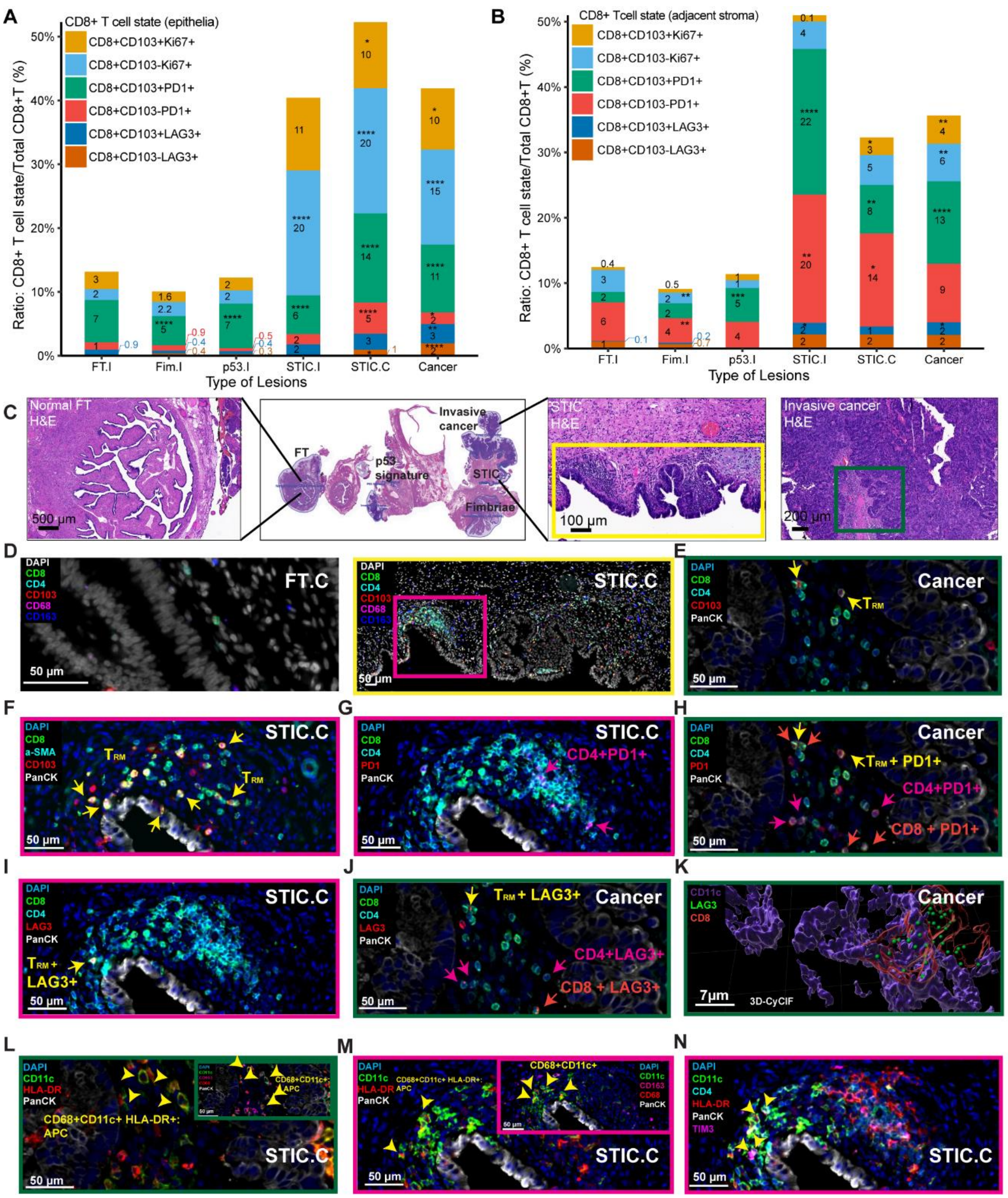

**Supp Fig. S24. Increased exhausted T population in the later stage of precancer and tumor:**

A-B. Stacked bar plots showing the proportion of CD8+ T cell state in total CD8+ T cells from single cell-CyCIF analysis in the epithelia (A) and in the adjacent stroma (B) across disease stages. CD8+ T cells are divided into CD8+ CD103- (i.e. cytotoxic, CTL) and CD8+ CD103+ T (i.e. Tissue resident memory T cells, $T_{RM}$ , also expressing CD45RO). Number of specimens per group as follows: FT.I (n=13), Fim.I (n=15), p53.I (n=10), STIC.I (n=9), STIC.C (n=23) and Cancer (n=20). Black asterisks indicate significant differences in stages compared to the FT.I; *p<0.05, **p<0.01, ****p<0.0001. Binomial GLMMs, with patient ID as random effect. Average proportions were rounded up to the next whole number when applicable and shown for each cell state across lesion types.

C. H&E image showing one of the STIC.C (STIC with concurrent HGSOC), also shown in **Figure 1F** (Case RD-23-002, patient ID 9, BRCA2 mutant, Stage IC HGSOC) with different histology.

D-E. CyCIF image showing major immune populations on matched FT.C, STIC.C and cancer components, namely in the adjacent stroma including $T_{RM}$. STIC.C was outlined with a yellow box and cancer was outlined with a green box. $T_{RM}$ was indicated by yellow arrows (E). F. CyCIF image showing CD8 populations, including $T_{RM}$ (yellow arrows), both intra epithelial and in adjacent stroma in STIC.C, outlined with the magenta box in panel D.

G. One of the other subtypes of T cells, CD4+ helper T cells co-expressing PD1+ (magenta arrows) in the adjacent stroma of STIC.C, indicating an activation state of CD4.

H. CyCIF image showing CD8+ (orange arrows) or $T_{RM}$ (yellow arrows), CD4+ T cells co-expressing PD1 (magenta arrows), indicating an activation state of these T cell populations in cancer.

I-J. Both in STIC.C (I) and tumor-adjacent stroma (J), CyCIF images show that some of these T cell populations also expressing LAG3+, indicating an exhausted phenotype. T cell populations were shown as follows: CD8+ (orange arrows) or $T_{RM}$ (yellow arrows), CD4+ T cells co-expressing LAG3 (magenta arrows).

K. Due to LAG3's punctate structure, 3D CyCIF and render settings from the same specimen (adjacent stroma to the tumor) reconfirm the presence of LAG3+ CD8+. In this instance, CD8+ T cell was in close contact with an APC (CD11c+).

L-M. CyCIF images showed not only a T cell population present, but APCs were frequent in the adjacent stroma of the tumor (L) and STIC.C (M). Yellow arrowheads confirm these APCs are CD68+ CD11c+ HLA-DR+, indicating potential macrophage derived APCs presenting antigens.

N. CD11c+ HLA-DR+ TIM3+ populations were shown with yellow arrowheads in STIC.C, indicating some of these APCs potentially had reduced capacity of antigen presentation.

F, G, I, M, N. STIC.C was outlined with a magenta box shown in the panel D.

E, H, J-L. Adjacent stroma of the tumor was outlined with a green box, shown in panel C.

**Supp Figure S25 (Related to Figure 6)**

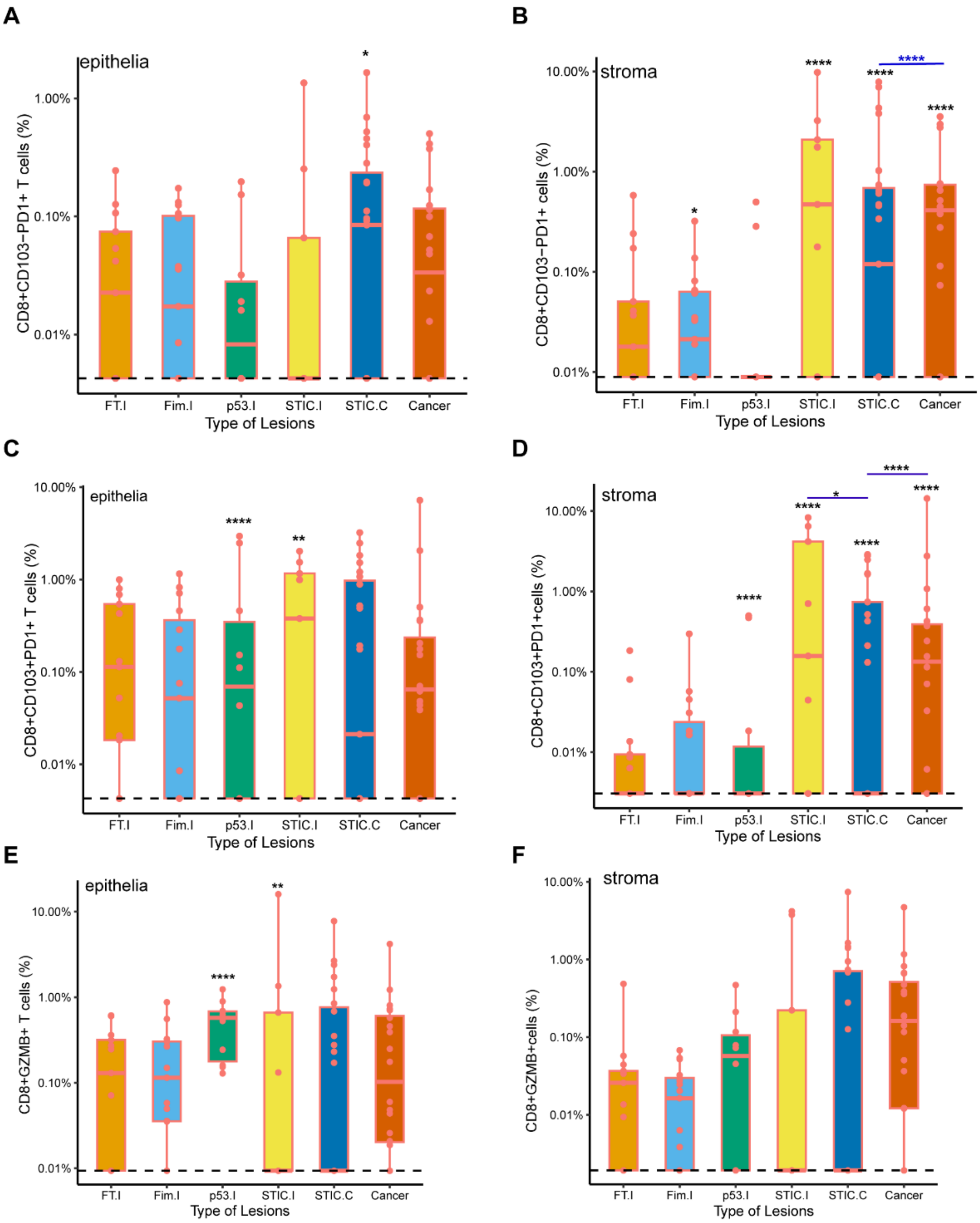

**Supp Figure S25. T cell subset:**

A-F. Box plots with individual data points showing CD8+ T cell subset with PD1 co-expression (A-D) and GZMB co-expression (E-F), expressed as a percentage. Number of specimens per group as follows: FT.I (n=13), Fim.I (n=15), p53.I (n=10), STIC.I (n=9), STIC.C (n=23) and Cancer (n=20). Black asterisks indicate significant differences in stages compared to the FT.I and blue lines with asterisks indicate comparison between groups; *p<0.05,**p<0.01****p<0.0001. GLMM, taking patient ID as a random effect. Y axis is represented as log10 scale, the dashed line represents values with zero. Stroma = adjacent stroma to the epithelium.

**Supp Figure S26 (Related to Figure 6)**

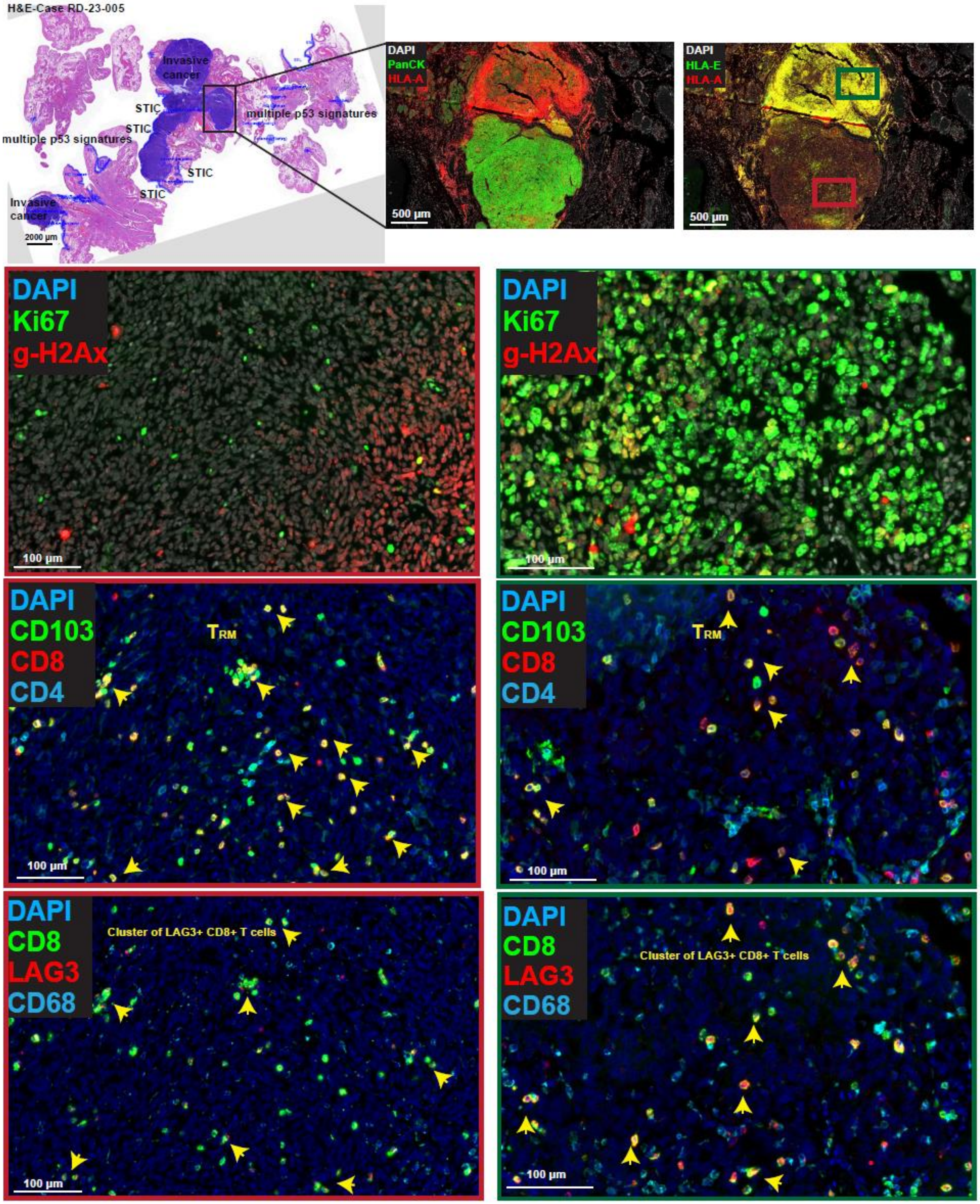

**Supp Figure S26**. A representation of invasive cancer co-existing with STIC (case RD-23-005, patient ID 12, BRCA1 Mut, Stage IC HGSOC), shown in **Supp Figure S10**. This CyCIF image illustrates heterogeneity in the intensity of HLA-E overexpression followed by proliferating cells and ɣ-H2Ax+ cells indicative of DNA-damage. This image also shows the presence of intra-tumoral $T_{RM}$ and cytotoxic T cells, co-expressing LAG3 (yellow arrows), indicating their exhausted status.
