## Supplementary material for "Multimodal Spatial Profiling Reveals Immune Suppression and Microenvironment Remodeling in Fallopian Tube Precursors to High-Grade Serous Ovarian Carcinoma": Supp Methods

**SUPPLEMENTARY METHODS**

Tissue Processing for CyCIF and GeoMx

Blocks of confirmed cases were freshly cut and immediately processed upon arrival for CyCIF. In general, after the sectioning for H&E, the second serial section was used for CyCIF immediately and the next adjacent section was kept for GeoMx at 4°C. The latter was also sectioned in an RNAse-free environment to reduce RNA degradation. All specimens for both assays were taken from adjacent sections except for 3 incidental p53 signatures (patients 31, 33, and 34). For those 3 cases, the section used for GeoMx was distant from the CyCIF section (~20um). Diagnostic H&E and/or off-bottom H&E images were annotated by board-certified pathologists (R.D, S.C, N.S). We had 10 incidental p53 signatures; however, it was not confirmed only by the H&E diagnosis. Due to the nature of histopathological features of p53 signatures, on H&E, they would look like layers of normal epithelium. Thirty BSO specimens were run as part of the first cycle of CyCIF with TP53, PanCK, and HLA-E antibodies. Then, based on both H&E and TP53 expression and intensity from CyCIF, p53 signatures, as well as Fimbriae and/FT, were annotated by a pathologist (S.C). Ten out of these 30 BSO specimens had confirmed p53 signatures based on the TP53 expression by CyCIF and were used for this study to proceed with further cycles of CyCIF. 3D CyCIF specimen was processed as 2D CyCIF, except a thicker (20µm) section was used compared to 2D (5 µm).

ROIs for GeoMX were chosen based on reviewing CyCIF images annotated by a pathologist (S.C). To avoid any RNA degradation, it is recommended to perform GeoMx within 2-3 weeks of the arrival of the fresh slides. Therefore, we ran 6-7 cycles of CyCIF (15-20 antibodies), then chose ROIs based on CyCIF imaging and processed ROI collection from GeoMx from the adjacent section. We completed the remaining CyCIF cycles up to 13 while preparing the libraries from collected GeoMx ROIs.

### Data Processing, quality control and downstream analysis from GeoMx spatial transcriptomics

The initial data was processed as recommended by NanoString using GeoMx DSP software, NanoString (v 3.1.0.221). The QC steps, and Quartile-3 (Q-3) normalization process was followed as recommended by NanoString. In brief, first total 603 segments or ROIs were collected including some immune control and necrotic areas. In total, 552 ROIs in total were chosen for further downstream analysis. 25/51 ROIs were removed based on very low raw read depth (<1000 raw read counts) and not being able to detect at least 5% of the genes detected above the limit of quantification. Overall, three strict criteria were followed to remove the segments as recommended: i. low raw read counts, ii. <80% of genes are aligned and iii. <50% sequence saturation (i.e. might not have sequenced deep enough). Following this initial QC, BioQC was followed as recommended, indicating a global probe QC. There are 2 set criteria: i. geomean probe in all segments (geomean probes within target <= 0.1) and ii. Fails Grubbs outlier test in >=20% of segments (if probe fails, remove only from the local segment). Following this, the Limit of Quantification (LOQ) was calculated using 2 standard deviations from the geomean of negative probes. This step is for counting gene-level data (count matrix file). Then we further filter segments whereby at least 5% of genes detected above LOQ. Furthermore, one “No template control” was usually recommended to be included in a 96 well plate collection, and No template control >1000 indicates RNAse in bench space. We did not have cases indicating there was RNAse contamination. <10 nuclei count was chosen to be flagged and manual inspection was performed for those segments. Due to the ROIs being smaller in the precancer area, the segments were not removed as NanoString recommended. Then the Q3-normalisation is performed using DSP software, normalizing every segment to its own Q3 counts (75% of gene percentile). The Q3-normalised file (count matrix file) is in **Supplementary File S8** and was used for downstream analysis. One of the STIC cases with concurrent HGSOC had severe tissue loss both in CyCIF and GeoMx (GeoMx slide ID: LSP16163). Hence, we removed that entire case (case ID RD-SS-009) from this study and did not include collected ROIs from that specimen (only normal FT were collected due to the tissue loss) in the ROI-annotation file. We then excluded those immune controls and necrotic ROIs since these were collected as a technical control for another project.

Following those excluded case and ROIs, in total, 567 ROIs passed QC that includes cancer (floating) (n=15) and STIL associated with cancer (n=10). The annotation file (**Supplementary File S2** contains all ROI (n=567) passed QC excluding the ROIs collected for another study/case with tissue loss) since the full data set will be publicly available. The Q3 normalized count matrix file contains all ROIs passed QC, including ROIs collected for another study, including the case with tissue loss. Due to data integrity, the count matrix file contains all ROI passed QC combining all batches of specimen processed.

Of those 567 ROIs, we included 542 for all the downstream analysis for this study. Floating cancer (n=15) was not considered part of invasive cancer. Furthermore, 10 ROIs from STIL lesions (cancer group) were not included in the downstream analysis due to small ROI numbers and no matched incidental STIL lesions (**Supplementary Fig S2**).

Then, Principal component analysis (PCA) was run to see the differences in gene expression between epithelial and stroma or incidental and cancer groups. Only six segments appeared to be cluster together due to a lower read depth in general, although they passed QC. Those ROIs were from incidental p53, fimbriae, or cancer stroma. Those 6 ROIs were kept in the analysis because LMM for differential gene expression, does take patient and ROI variability into account. Hence, since these 6 ROIs passed QC, it was recommended to keep them in downstream analysis by NanoString. We ran all these samples in four batches, but no batch effect was observed; therefore, no batch correction was performed.
